# Computational methods to develop potential neutralizing antibody Fab region against SARS-CoV-2 as therapeutic and diagnostic tool

**DOI:** 10.1101/2020.05.02.071506

**Authors:** Hemanth K. Manikyam, Sunil K. Joshi

**Affiliations:** Faculty of Science, North East Frontier Technical University, Arunachal Pradesh, India; Division of Haematology / Oncology & Bone Marrow Transplantation, Department of Pediatrics, University of Miami Miller School of Medicine, Miami, FL USA

**Keywords:** SARS-CoV-2, coronavirus, IMGT, Human antibodies, I-TASSER, IgBLAST, Molbiol, LYRA, Fab, Nabs and mAbs

## Abstract

SARS-CoV-2, a global pandemic originated from Wuhan city of China in the month of December 2019. There is an urgency to identify potential antibodies to neutralize the virus and also as a diagnostic tool candidate. At present palliative treatments using existing antiviral drugs are under trails to treat SARS-CoV-2.Whole Genome sequence of Wuhan market sample of SARS-CoV-2 was obtained from NCBI Gene ID MN908947.3.Spike protein sequence PDB ID 6VSB obtained from RCSB database. Spike protein sequence had shown top V gene match with IGLV1-44*01, IGLV1-47*02 and has VL type chain. Whole Genome sequence had shown top V gene match with IGHV1-38-4*01 and has VH type chain. VD chain had shown link to allele HLA-A0206 80%, HLA-A0217 80%, HLA-A2301 75%, HLA-A0203 75%, HLA-A0202 70% and HLA-A0201 55% of binding levels. Some conserved regions of spike protein had shown strong binding affinity with HLA-A-0*201, HLA-A24, HLA-B-5701 and HLA-B-5703 alpha chains. Synthetic Fab construct BCR type antibody IgG (CR5840) had shown Polyspecific binding activity with spike glycoprotein when compared with available Anti-SARS antibody CR3022.Thus we propose CR5840 Fab constructed antibody as potential neutralizing antibody for SARS-CoV-2. Based on germline analysis we also propose cytotoxic T lymphocyte epitope peptide selective system as effective tool for the development of SARS-CoV-2 vaccine.

## Introduction

The SARS-CoV-2 causing COVID-19 pandemic threatened global public health and economy. Neutralizing antibodies (NAbs) elicited by virus during infection may correlate to patient recovery. However characterisation of neutralizing antibodies elicited during viral infection has not been well studied in relation with clinical outcome in the patients. Currently, neutralizing antibodies (NAbs) versus this virus are expected to correlate with recovery and protection of this disease. However, the characteristics of these antibodies have not been well studied in association with the clinical manifestations in patients. NAbs play crucial role in controlling the viral infection. Monoclonal antibodies (mAbs) functional antigen-binding fragment (Fab) is the form of NAbs developed for SARS-CoV and MERS currently developed. These mAbs target Receptor binding domain and N terminal domains. mAbs thus inhibit membrane fusion and restrict viral host cell entry.

Palivizumab is a monoclonal antibody (mAb) produced by recombinant DNA technology used to reduce the risk of respiratory syncytial virus in children. Phage-display technology a new method of developing promising method in antibody development by mAbs that utilizes fragment antigen binding (Fab) molecules as an intermediate. This technology had proven the neutralization effect of Fabs against influenza virus. Phage display technology an alternative method of producing monoclonal antibodies that utilizes bacteriophages that display fragments of antibodies Fabs from viral surface. Fab fragment of an antibody contain light kappa chain and heavy chain that binds actively to specific regions of antigens. Human neutralizing mAbs like CR6261, Navivumab, Firivumab and Diridavumab are presently used to treat Influenza virus infections. Human neutralizing mAbs S230.15 and m396 isolated from SARS-COV-1 infected patients had shown competitive inhibition with ACE2 for binding RBD. Highly conserved Residue regions are found to key factors for cross reactivity.

Polyclonal antibodies from SARS-CoV-2-infected patients been used to treat SARS-CoV-2 infection, but no SARS-CoV-2-specific neutralizing mAbs or Fabs have been reported. Researchers around globe are working hard to identify monoclonal antibodies or their functional fragments Fabs as prophylactic, therapeutic and diagnostic agents against COVID-19 caused by SARS-CoV-2. Since SARS-CoV-2 closely related to SARS-Co-V-2 the conserved residues with epitopes may similar structural and functional properties. Hence there are chances of utilising mAbs or it active fragments Fabs against COVID-19 for neutralizing or cross neutralizing activity which to be proved through proper clinical studies.

As it takes time to identify proper mAbs against SARS-CoV-2, we suggest application of computational methods to predict the mAbs based on structural confirmations submitted like SARS-CoV-2 spike proteins. Once specific mAbs or Fab regions are predicted and their sequence been obtained, Phage display technology can be used to synthesize Fab fragments or Monoclonal antibody technology can be used to produce mAbs as a therapeutic and diagnostic agents against COVID-19.

## Methods

### IgBLAST and IMGT (Immunogenetics) Analysis

IgBLAST was developed by NCBI for analysis of variable domain sequences of immunoglobulin^1, 4, 5^. IMGT is a platform to access sequence, genome and structure Immunogenetics data based on gene ontology. After immunoglobulin germline prediction we compare the V-J and V-D-J junction sequence with the extremities of the corresponding “V, J and D “germline” genes (see IMGT Repertoire > Alignments of alleles).^8, 9^

### Binding Site and Ligand Prediction

ProBis a computer based tool to predict binding site and corresponding Ligands for submitted protein structure. Spike protein sequence PDB ID 6VSB was obtained from RCSB database and submitted to ProBis to predict specific Ligands.^6, 10^

### I-TASSER (Iterative Threading Assembly Refinement)

I-TASSER is a computational approach to predict protein structure and function. It is an most accurate protein structure and function prediction algorithm.^11, 12^ After IMGT and ProBis analysis predicted V-D-J and Fab Ligands were obtained and submitted to I-TASSER for best functional model of ligand which can be obtained as PDB file.

### Lymphocyte Receptor Automated Modelling (LYRA)

Predicted protein ligand sequences from ProBis algorithm obtained from RCSB and compared for best possible alignments and combinations using IgBLAST and LYRA predictions tools. LYRA server predicts either T cell or B cell receptors based on homology modelling. Based on BLOSUM score Framework templates are selected and complementary determining regions (CDR) are then selected if needed based on a canonical structure model and grafted onto the framework templates. The final output is a complete three-dimensional model of the target antibody.^14^

### Stereochemical quality of a protein structure

PROCHECK EMBL-EBI algorithm checks the Stereochemical quality of a protein structure and produces a number of postscript plots analysing residue-by-residue geometry ^13^

### FRODOCK and Pydock protein-protein docking analysis

Constructed Fab fragment using I-TASSER and LYRA algorithms in PDB format was studied for Structural prediction of protein-protein interaction using FRODOCK and Pydock algorithms.^15, 16^ Spike protein structure obtained from RCSB database PDB ID: 6VSB. See table 1 and 2.

**Table 1:** Protein - Protein interface statistics of heavy chain and kappa light chain of Fab region.

| Chain | No. of Interface Residues | Interface Area (Å<sup>2</sup>) | No. of Salt Bridges | No. of Disulphide Bonds | No. of Hydrogen Bonds | No. of Non-Bonded Contacts |
| --- | --- | --- | --- | --- | --- | --- |
| H | 11 | 692 | - | - | 1 | 78 |
| L | 11 | 737 |  |  |  |  |

**Table 2:** Top 10 energies and Rank predicted for *CR5840* interaction with SARS-CoV-2 Spike protein (PDB ID: 6VSB).

| Conf | Electrostatics | Desolvation | VdW | Total | Rank |
| --- | --- | --- | --- | --- | --- |
| 9721 | -15.031 | -24.564 | -1.536 | -39.748 | 1 |
| 7731 | -8.751 | -32.630 | 28.402 | -38.541 | 2 |
| 5986 | -15.974 | -18.586 | -17.699 | -36.330 | 3 |
| 2796 | -8.088 | -32.259 | 42.454 | -36.102 | 4 |
| 4113 | -16.412 | -24.967 | 56.981 | -35.681 | 5 |
| 3627 | -5.174 | -29.850 | -6.292 | -35.653 | 6 |
| 502 | -9.962 | -27.967 | 29.594 | -34.970 | 7 |
| 309 | -17.792 | -19.174 | 20.341 | -34.932 | 8 |
| 6896 | -3.943 | -34.699 | 38.134 | -34.828 | 9 |
| 414 | -14.218 | -21.921 | 15.601 | -34.579 | 10 |

## Background

Presently world is facing pandemic caused by SARS-CoV-2 and tool more than 60,000 lives as per existing records. Any infections caused by microorganisms develop human neutralising antibodies after certain period of time. Microbial genome contains antigenic sequences that can trigger certain type of IMGT antibodies in the host which can be useful to design vaccines and also as diagnostic tools. IgBLAST and Molbiol tools are useful in detecting top gene matches and their germline antibodies using genome sequence of pathogens. We conducted IgBLAST analysis of whole genome and also Spike protein of SARS-CoV-2 coronavirus in order to identify possible antigen neutralizing human antibodies. Immune Epitope Database (IEDB) tools used to identify antigen sequence epitopes. Lymphocyte receptor automated modelling (LYRA) and I TASSER tools are used to construct the BCR antibody.

## Results

### IgBLAST and IMGT (Immunogenetics) Analysis

Sequences producing significant alignments with germline gene and top gene alignment shown in figure 1&3 for whole genome and figure 2&4 for spike protein genome. Imgt Domain Classification was studies for given sequence. Minus strand of a V gene and has been converted to the plus strand during IgBLAST. V-D-J rearrangement summary of query sequence shown in figure 1 for whole genome and figure 2 for spike protein genome. Alignments with top V gene hits are shown in supplementary file 1, file 2 and file 3. Spike protein sequence had shown top V gene match with IGLV1-44*01, IGLV1-47*02 and has VL type chain. Whole Genome sequence had shown top V gene match with IGHV1-38-4*01 and has VH type chain. VD chain had shown link to allele HLA-A0206 80%, HLA-A0217 80%, HLA-A2301 75%, HLA-A0203 75%, HLA-A0202 70% and HLA-A0201 55% of binding levels. See supplementary file 3.Some conserved regions of spike protein had shown strong binding affinity with HLA-A-0*201, HLA-A24, HLA-B-5701 and HLA-B-5703 alpha chains.

**Figure 1.**
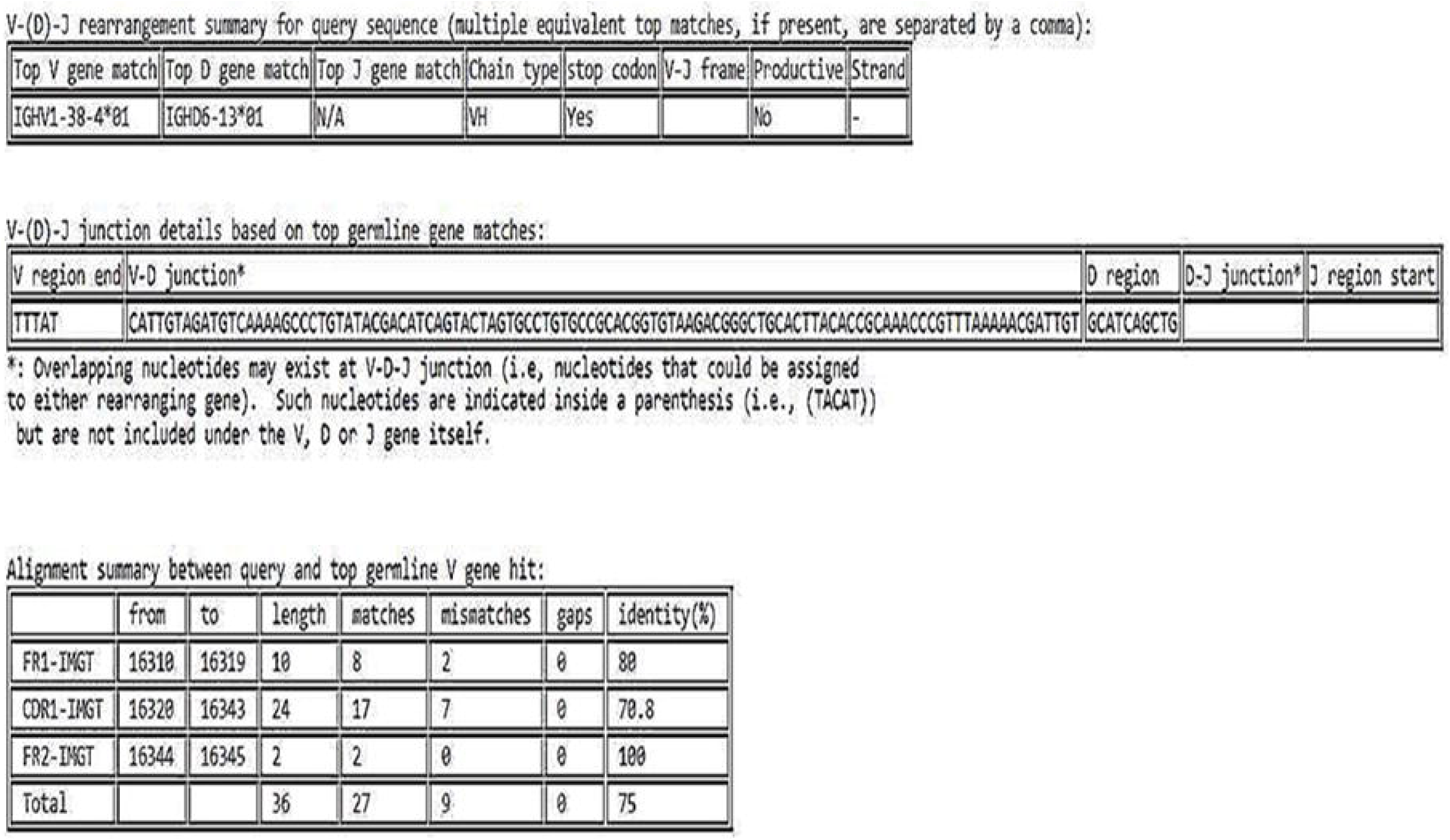
Predicted V-D-J junctions and top germline V gene hits for the query sequence of Gene ID MN908947.3

**Figure 2.**
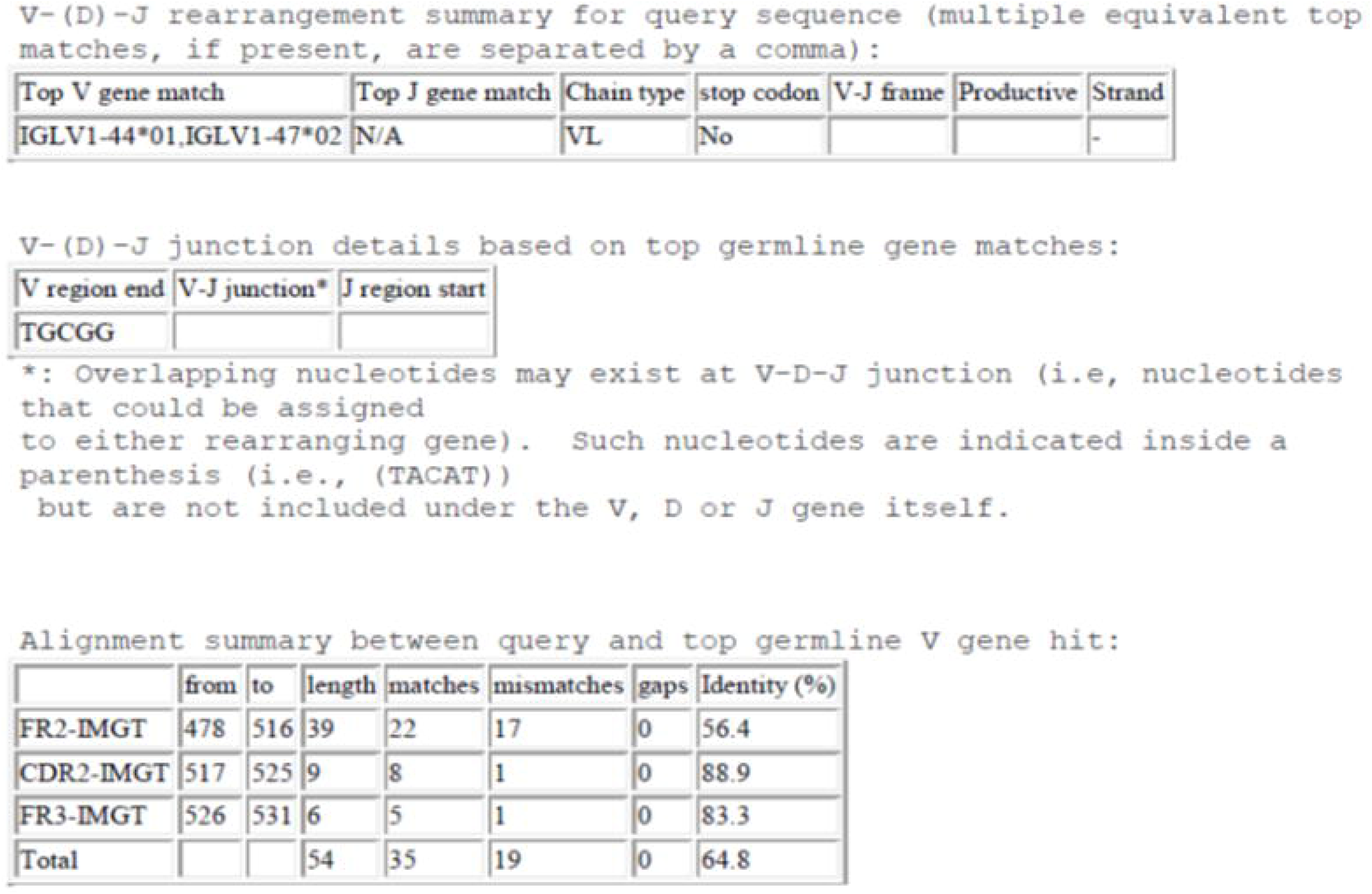
Predicted V-D-J junctions and top germline V gene hits for the query sequence of Spike protein side chain A PDB ID 6VSB

**Figure 3.**
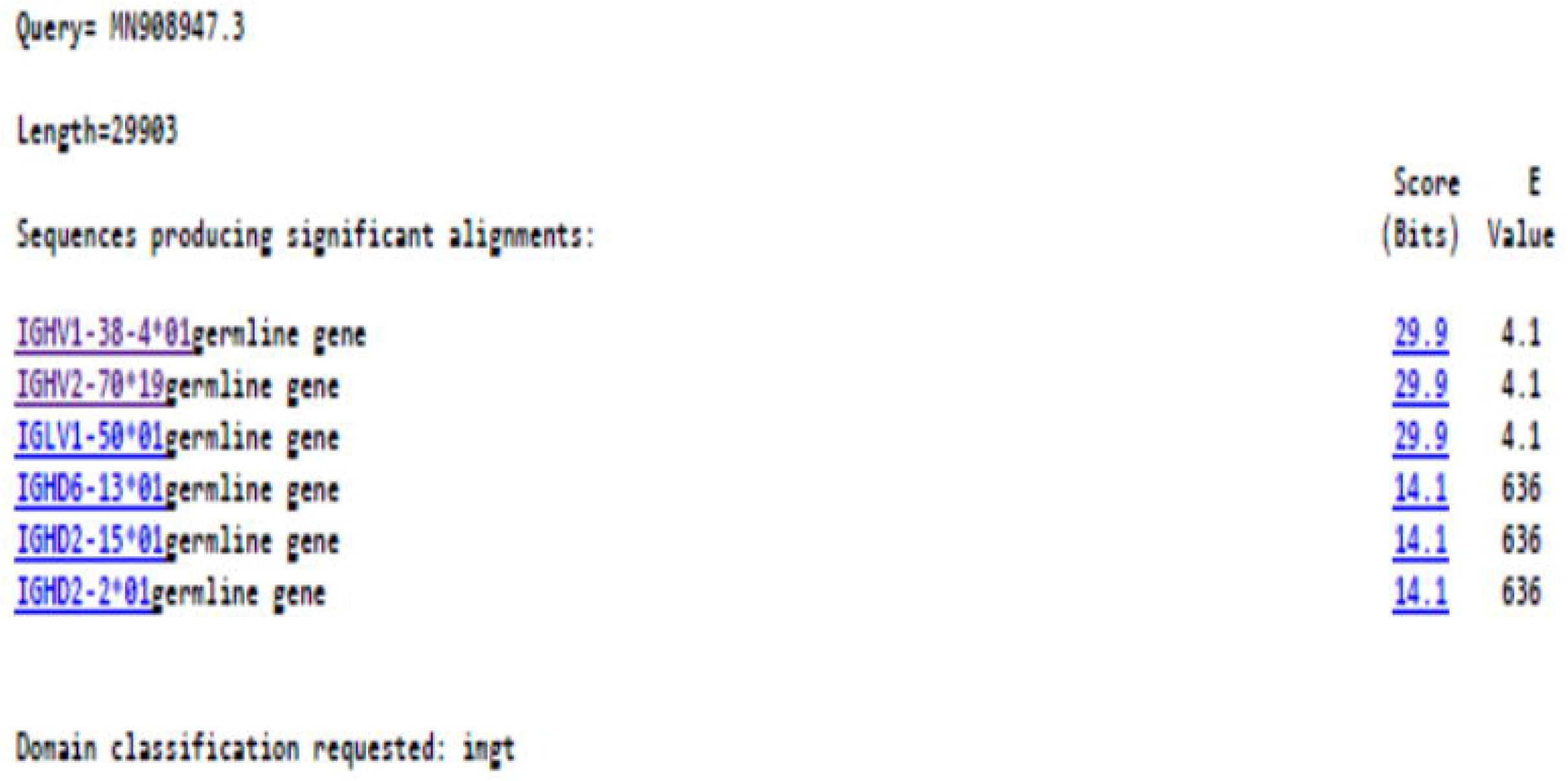
Sequences producing significant alignments for query sequence whole genome of SARS-CoV-2

**Figure 4.**
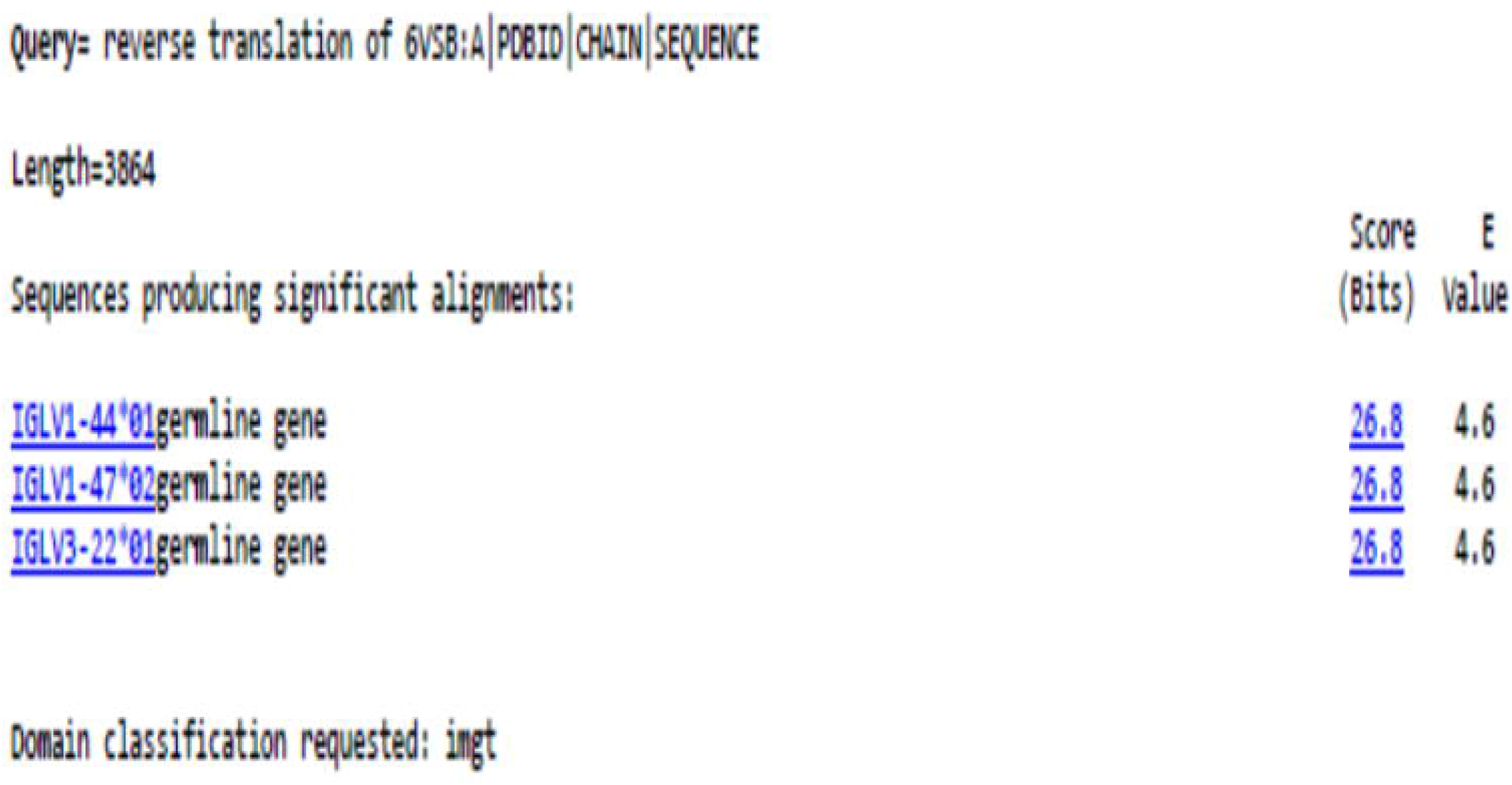
Sequences producing significant alignments for query sequence Spike protein chain A genome of SARS-CoV-2

### ProBis, I-TASSER and LYRA analysis

From epitope and germline data based on spike protein sequence, ProBis ligand prediction tools generated antibodies that are specific to spike glycoprotein. From the sequence of specific antibodies obtained, we generated synthetic IgG hybrid antibody Fab sequence. LYRA and I-TASSER tools predicted possible structural conformation from heavy chain and kappa light chain and generated final hybrid antibody CR5840 see figure 5 and 6

**Figure 5.**
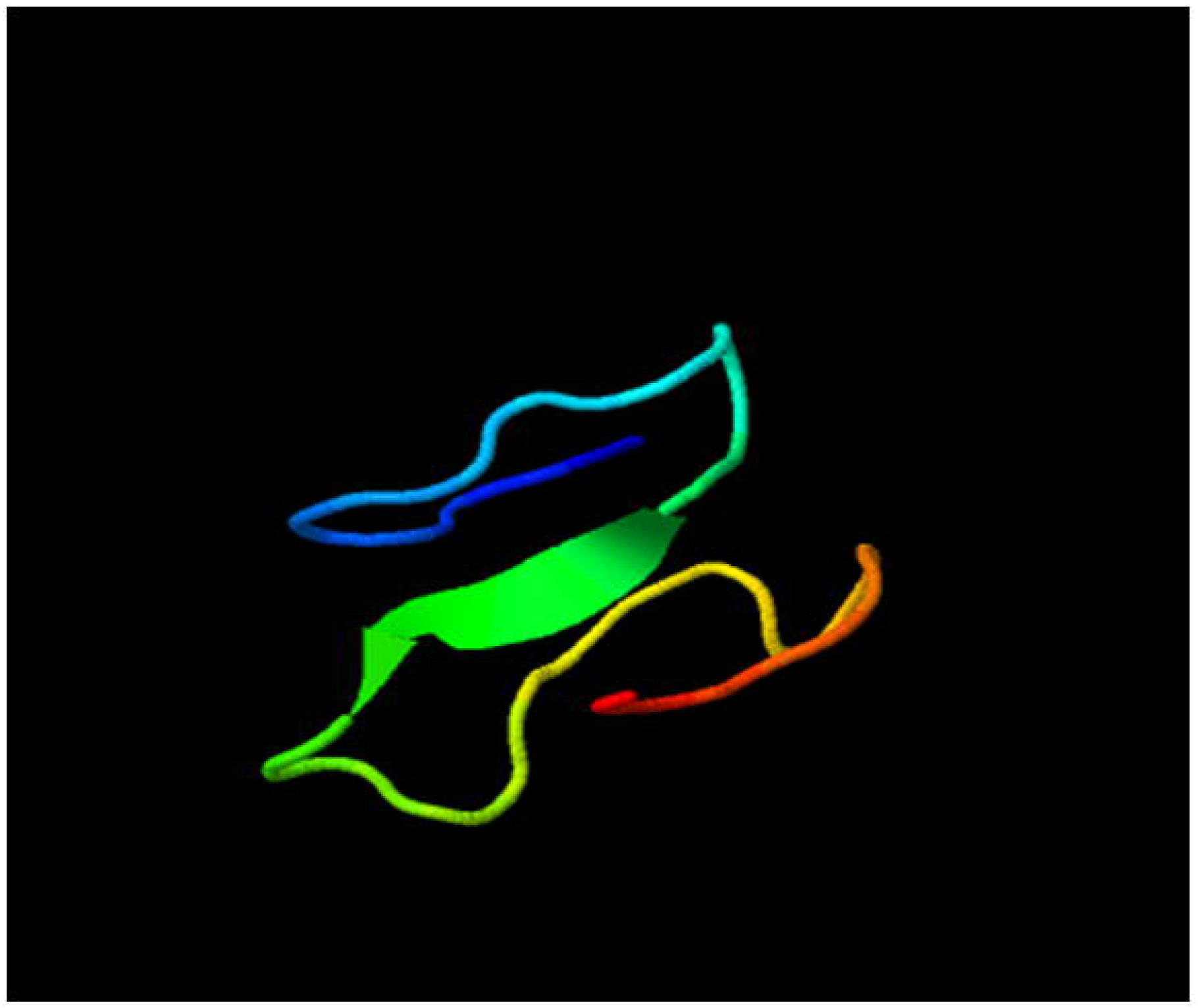
Top I-TASSER structural confirmation for VD chain for whole genome sequence of SARS-CoV-2

**Figure 6.**
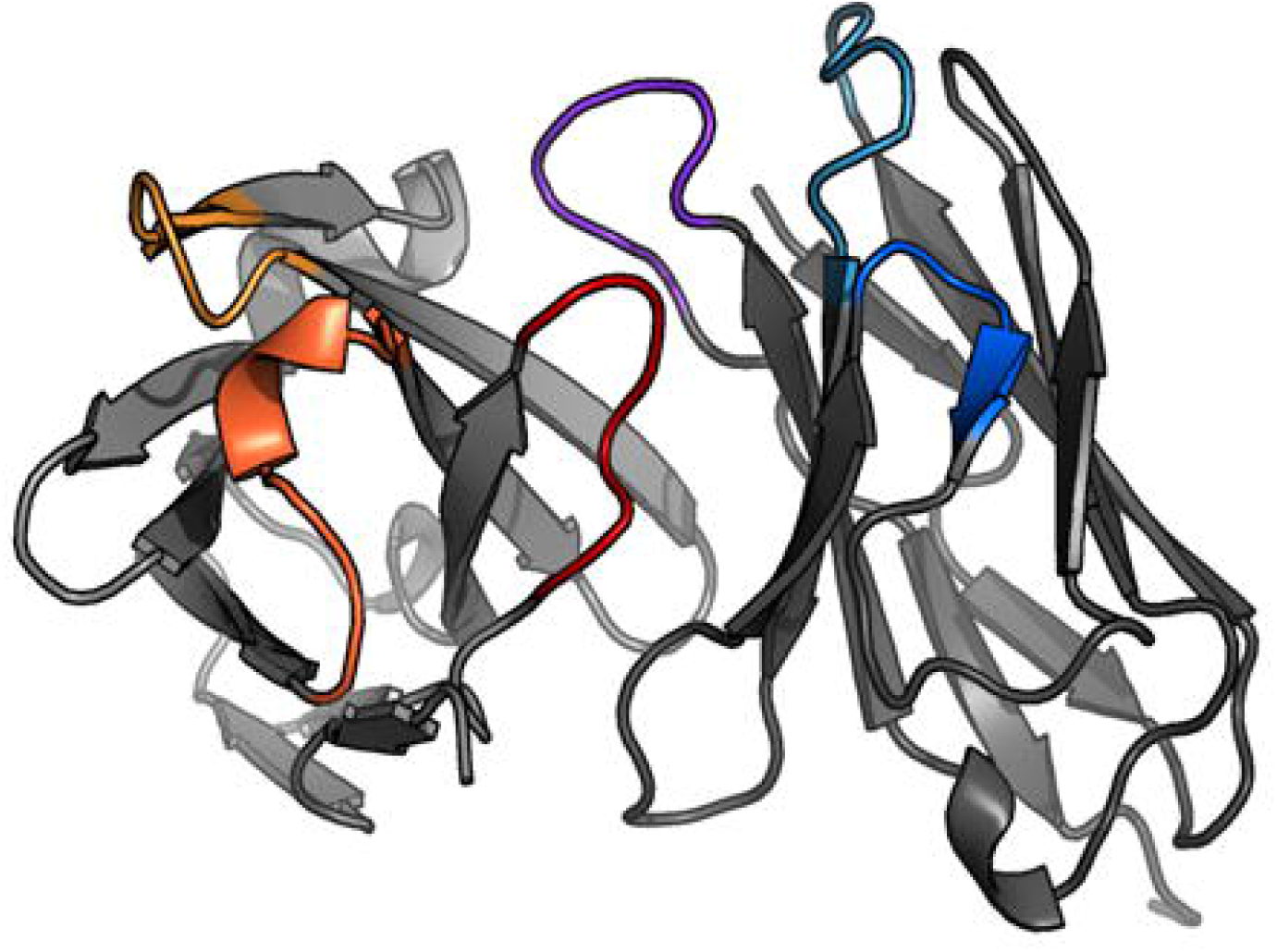
heavy and kappa light chains looped - final predicted structure (CR5840)

### Stereochemical quality of a protein structure

Quality analysis of structurally confirmed Fab fragment annotated as CR5840 was conducted using PROCHECK EMBL-EBI algorithm. Ramchandran plot, residue to residue analysis, RMS distance analysis with main chain bond length, angle analysis and parameters obtained see figure 9, 10 and 11. Protein-protein interface of heavy chain and kappa light chain of Fab fragment shown in figure 12. Interacting chains are joined by coloured lines, each representing a different type of interaction, as per the key above. The area of each circle is proportional to the surface area of the corresponding protein chain. The extent of the interface region on each chain is represented by the black wedge whose size signifies the interface surface area. Statistics for this interface are given Table 1. Residue interactions shown in figure13. The number of H-bond lines between any two residues indicates the number of potential hydrogen bonds between them. For non-bonded contacts, which can be plentiful, the width of the striped line is proportional to the number of atomic contacts see figure 13. Topology of Fab fragment shown in figure 8. Cleft, tunnel and Pores analysis had figure 14, 15, 16 and supplementary file 4. Binding site analysis shown in table. List of all matched Gene Ontology (GO) terms, Reverse template with certain matches (E-value < 1.00E-06) and list of predicted matches for CR5840 were presented in supplementary file 5. Most of the gene ontology predicted CR5840 structure for immunoglobulin IgG complex having cell surface interactions.

**Figure 7.**
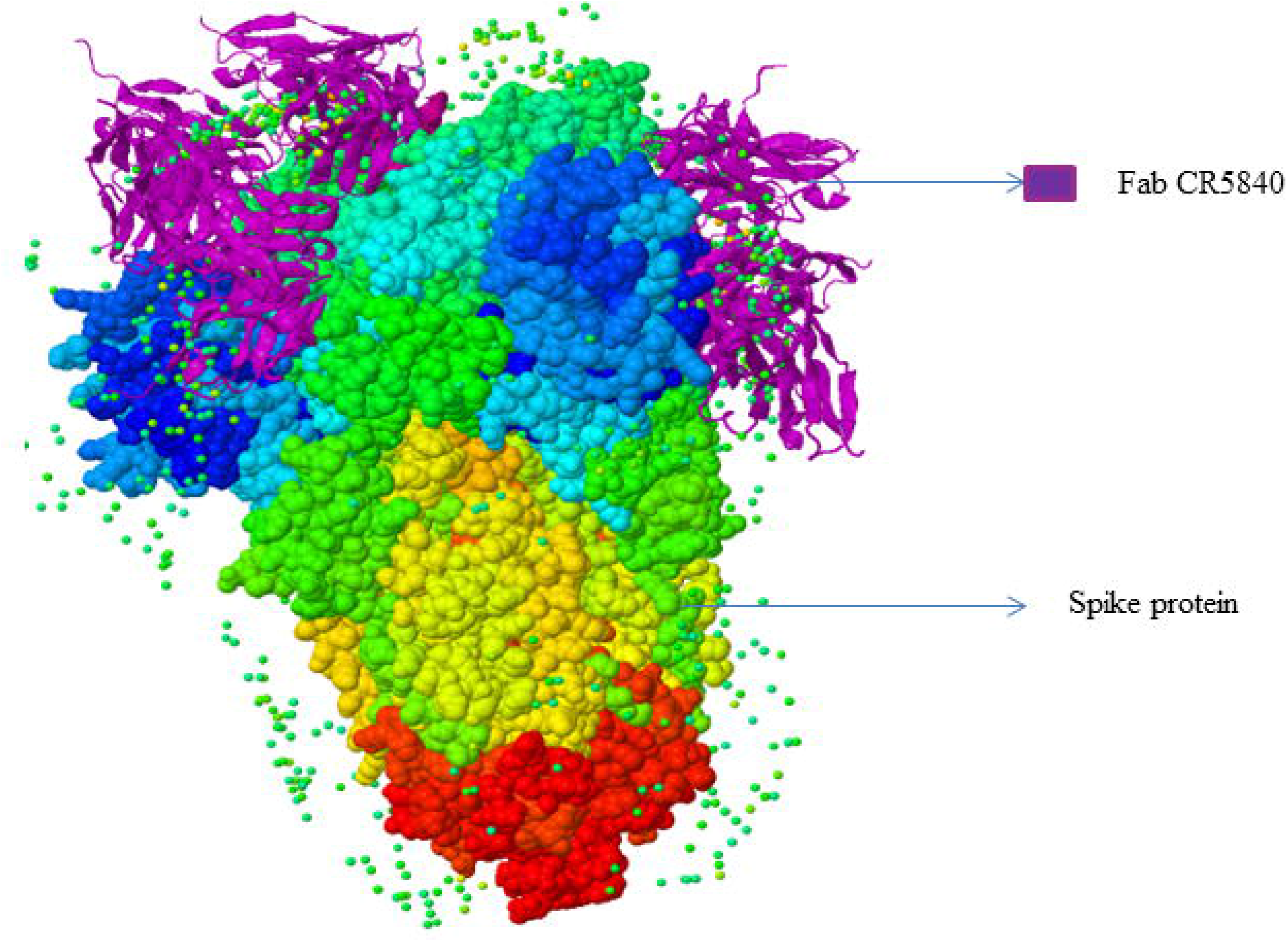
synthetic hybrid antibody confirmatory structures and interaction with spike glycoprotein of SARS-CoV-2

**Figure 8.**
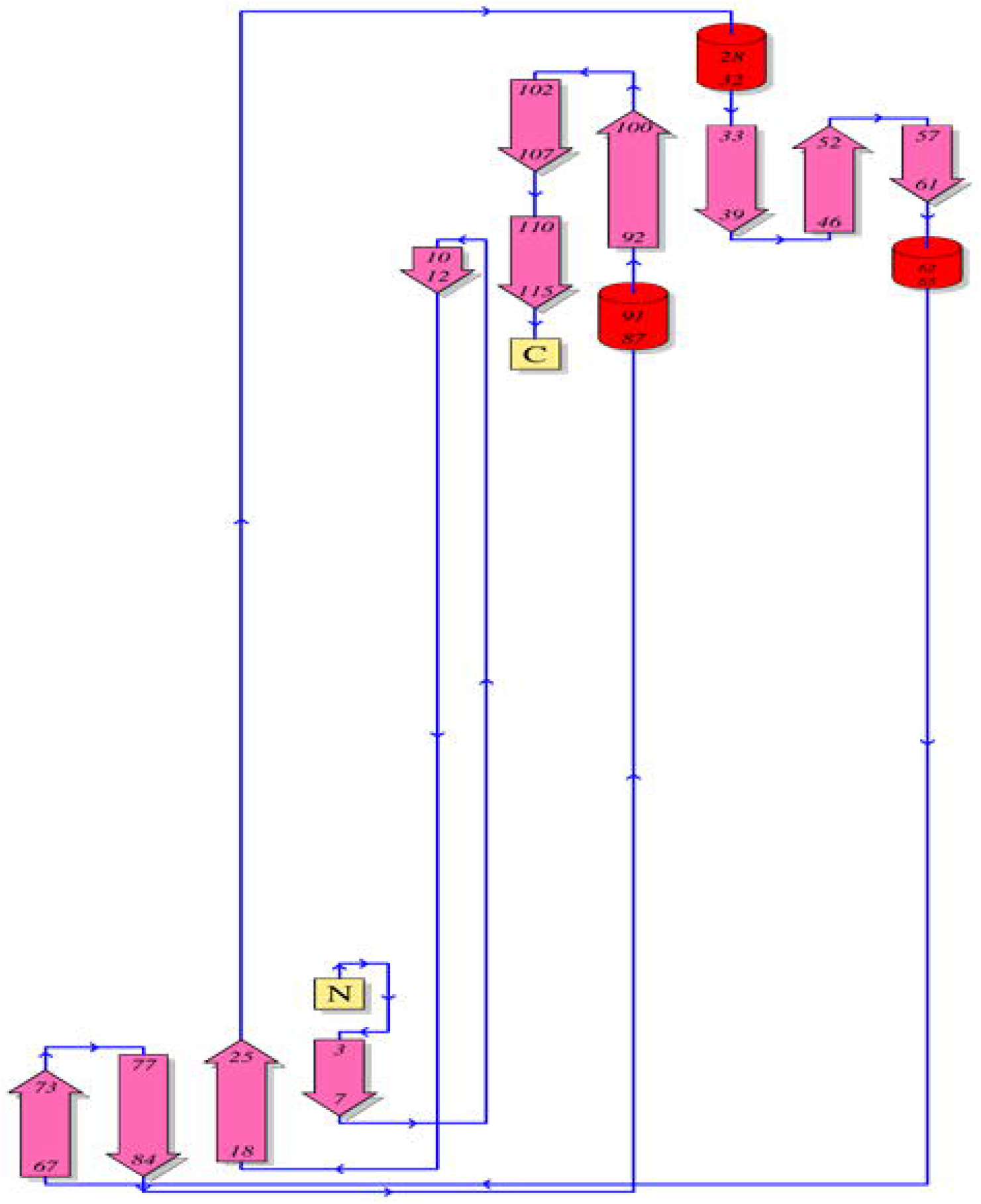
Topology of heavy and kappa light chain looping

**Figure 9.**
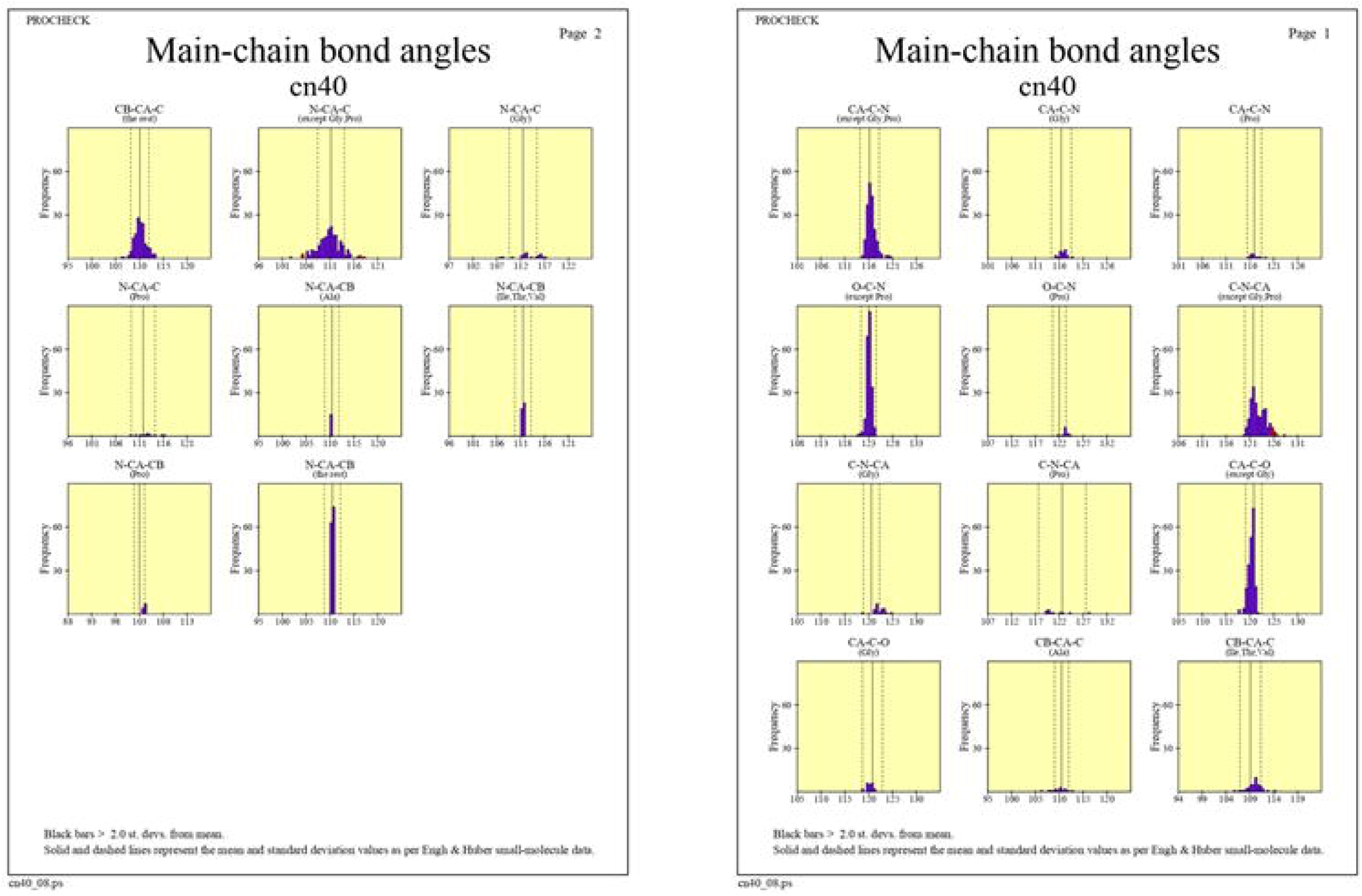
Main chain bond angles of CR5840 Fab region

**Figure 10.**
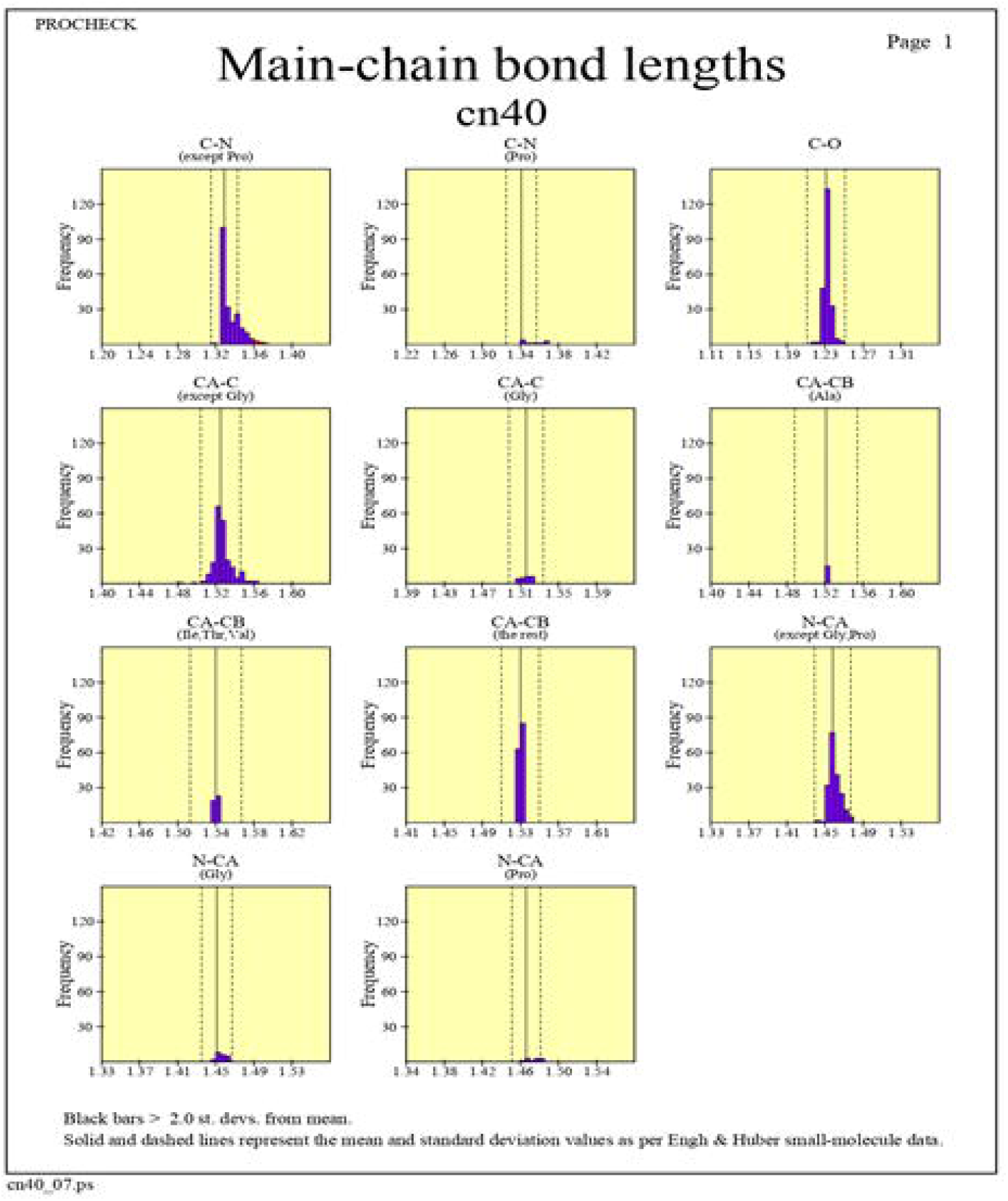
Main chain bond Lengths of CR5840 Fab region

**Figure 11.**
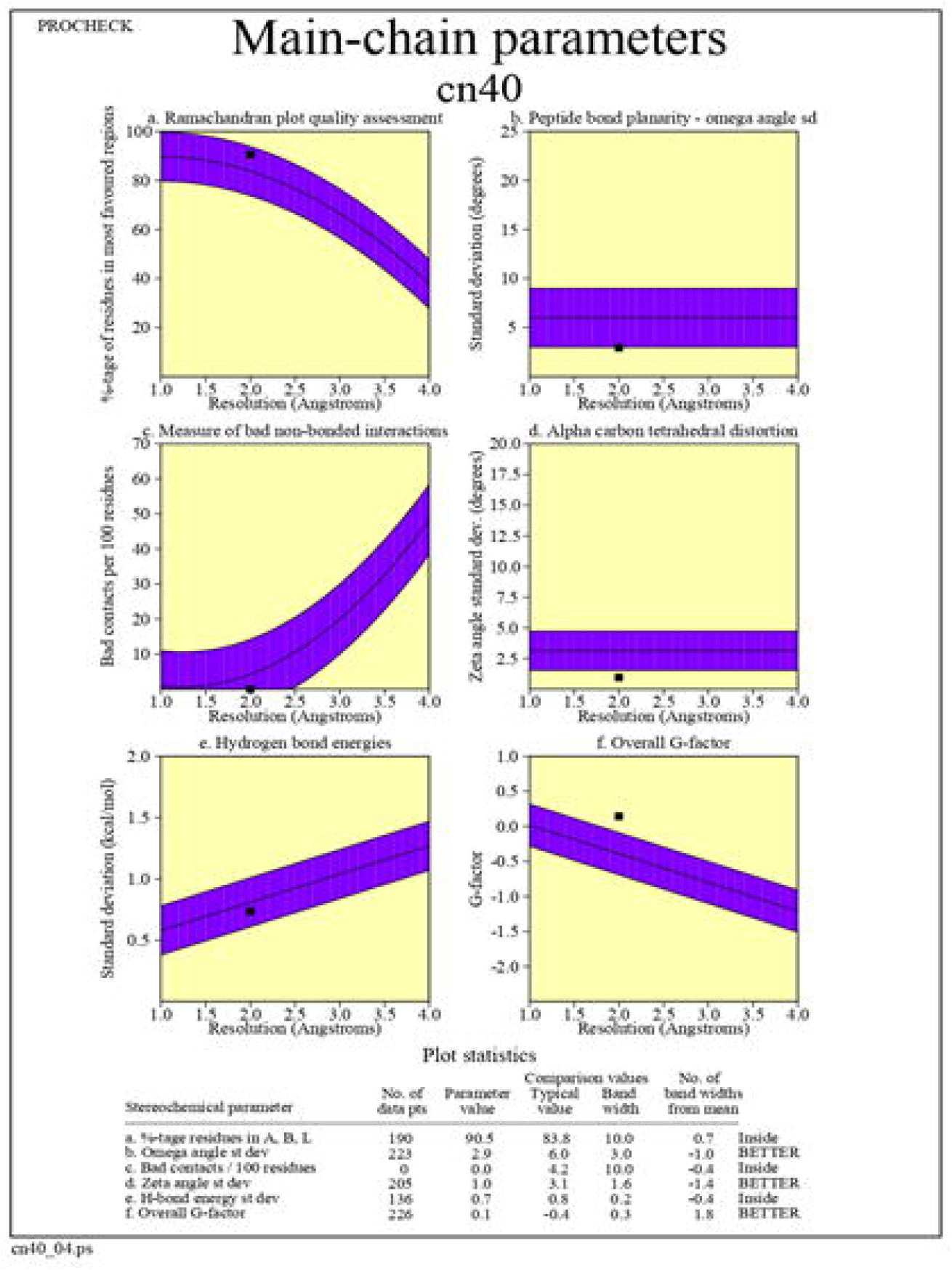
Main chain Parameters of CR5840 Fab region

**Figure 12.**
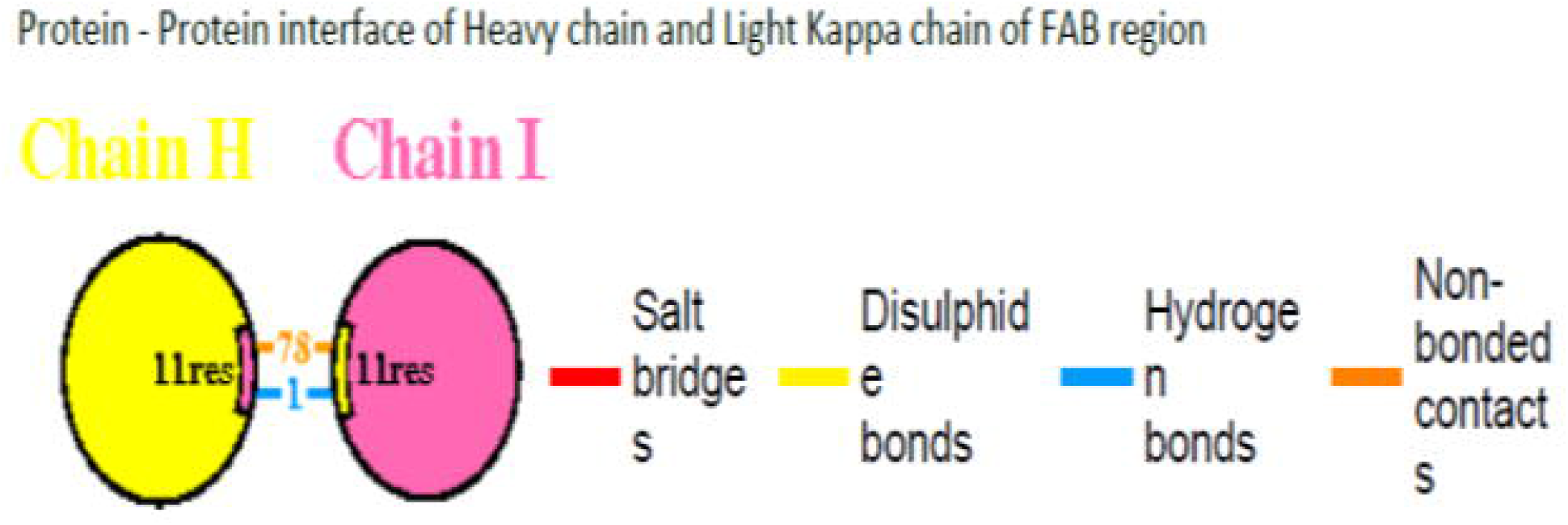
Protein-Protein interface of Heavy chain and Light Kappa chain Fab region

**Figure 13.**
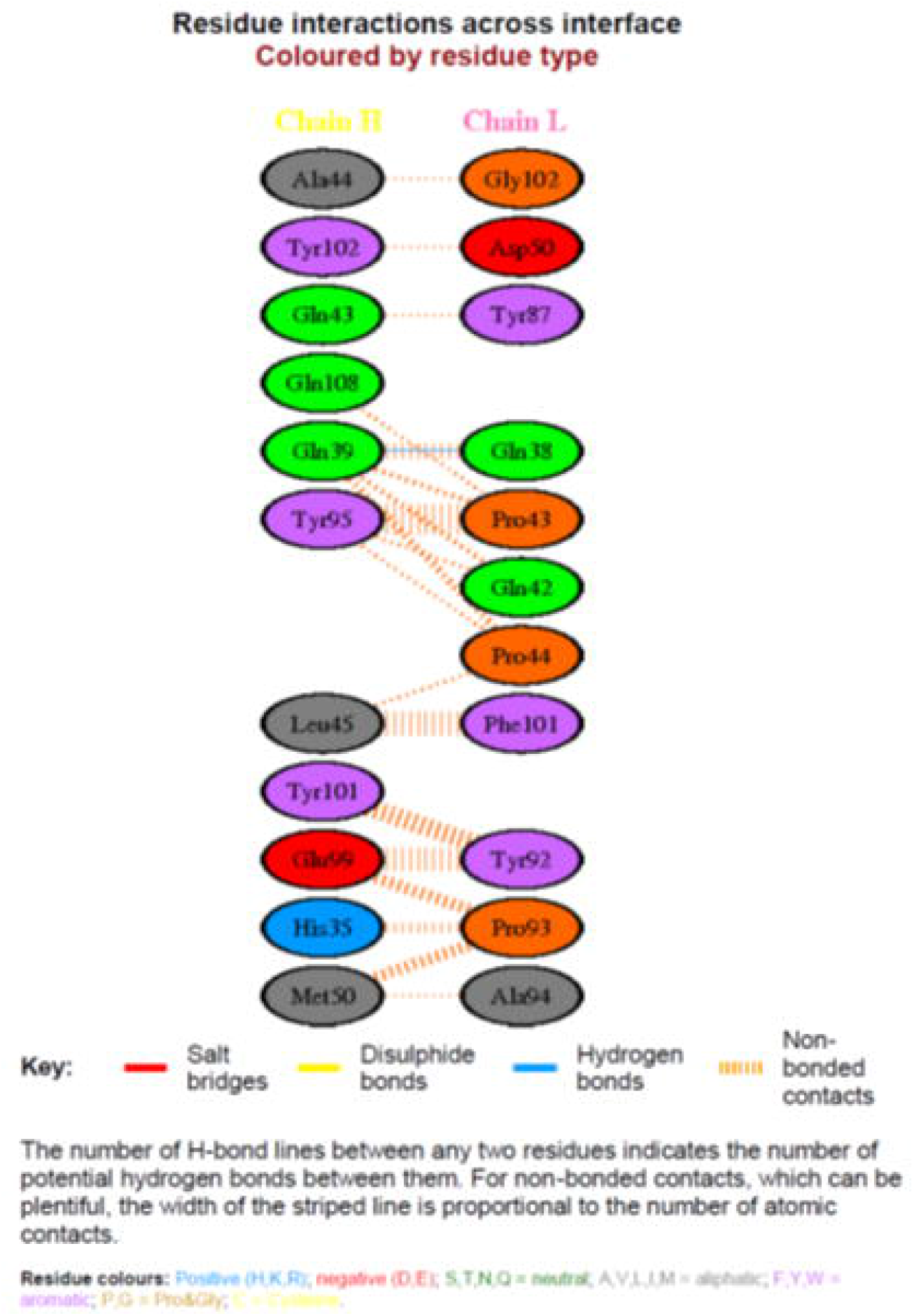
Residue interactions across intetface of heavy and light kappa

**Figure 14.**
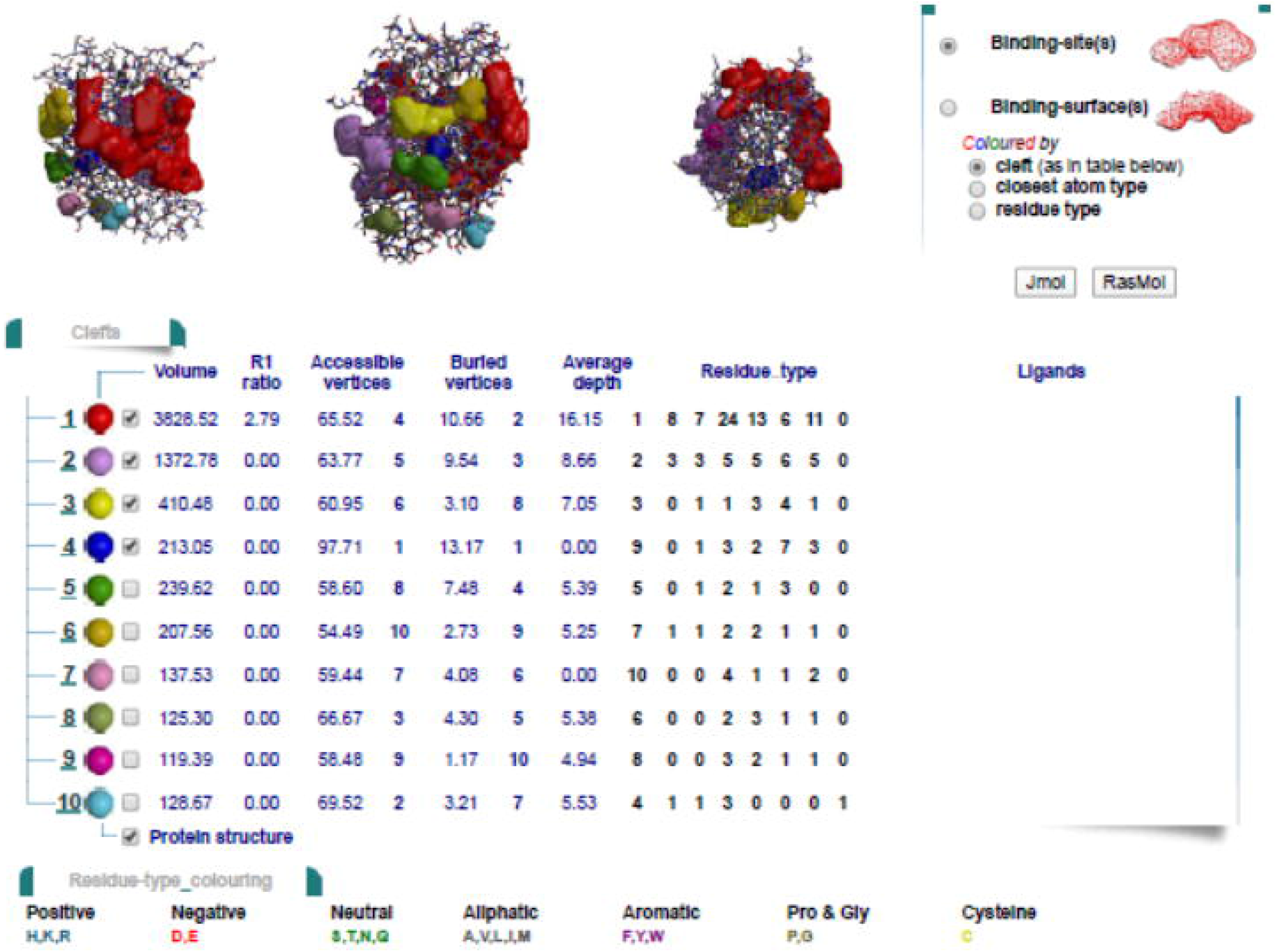
Cleft analysis

**Figure 15.**
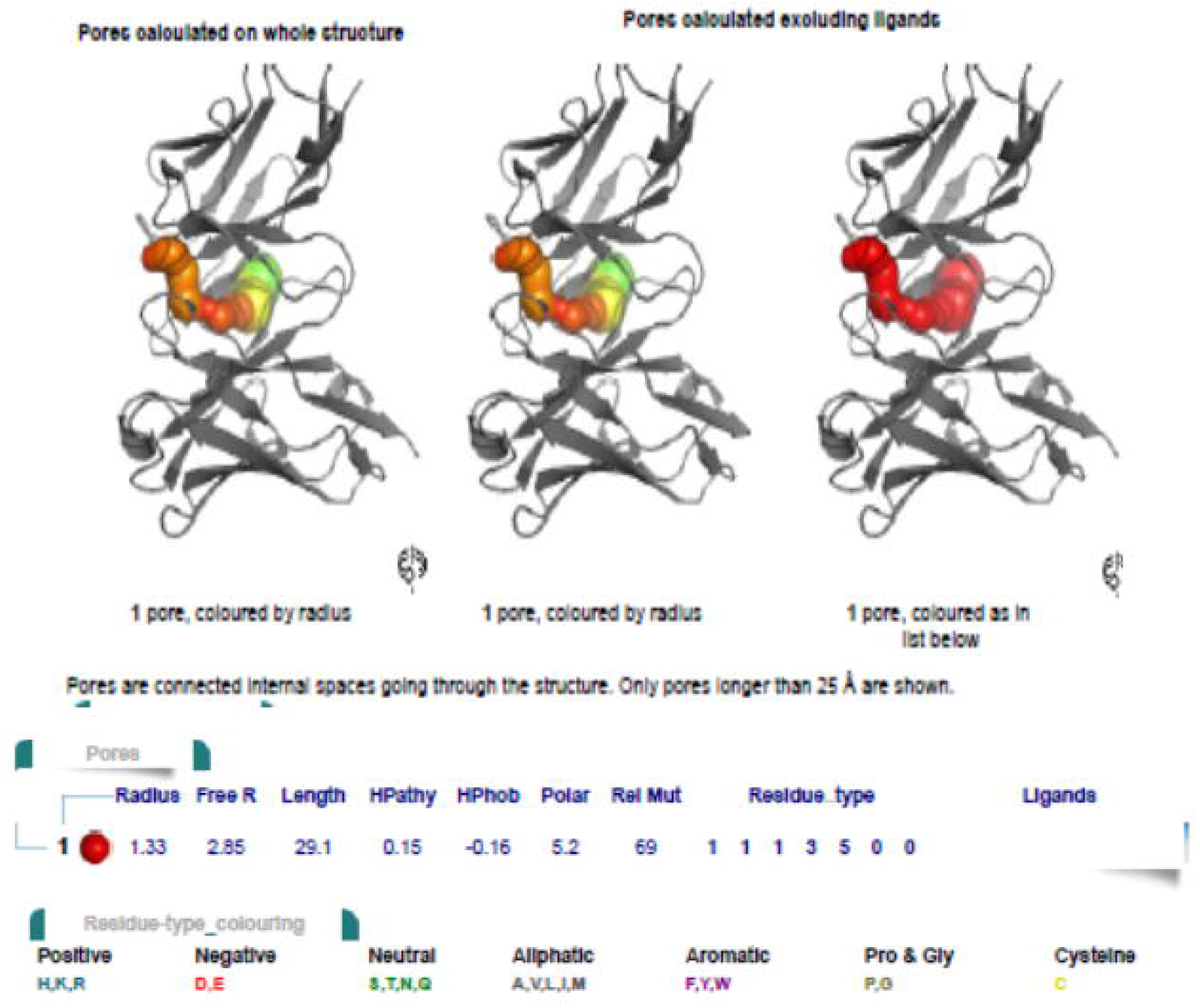
Pores analysis

**Figure 16.**
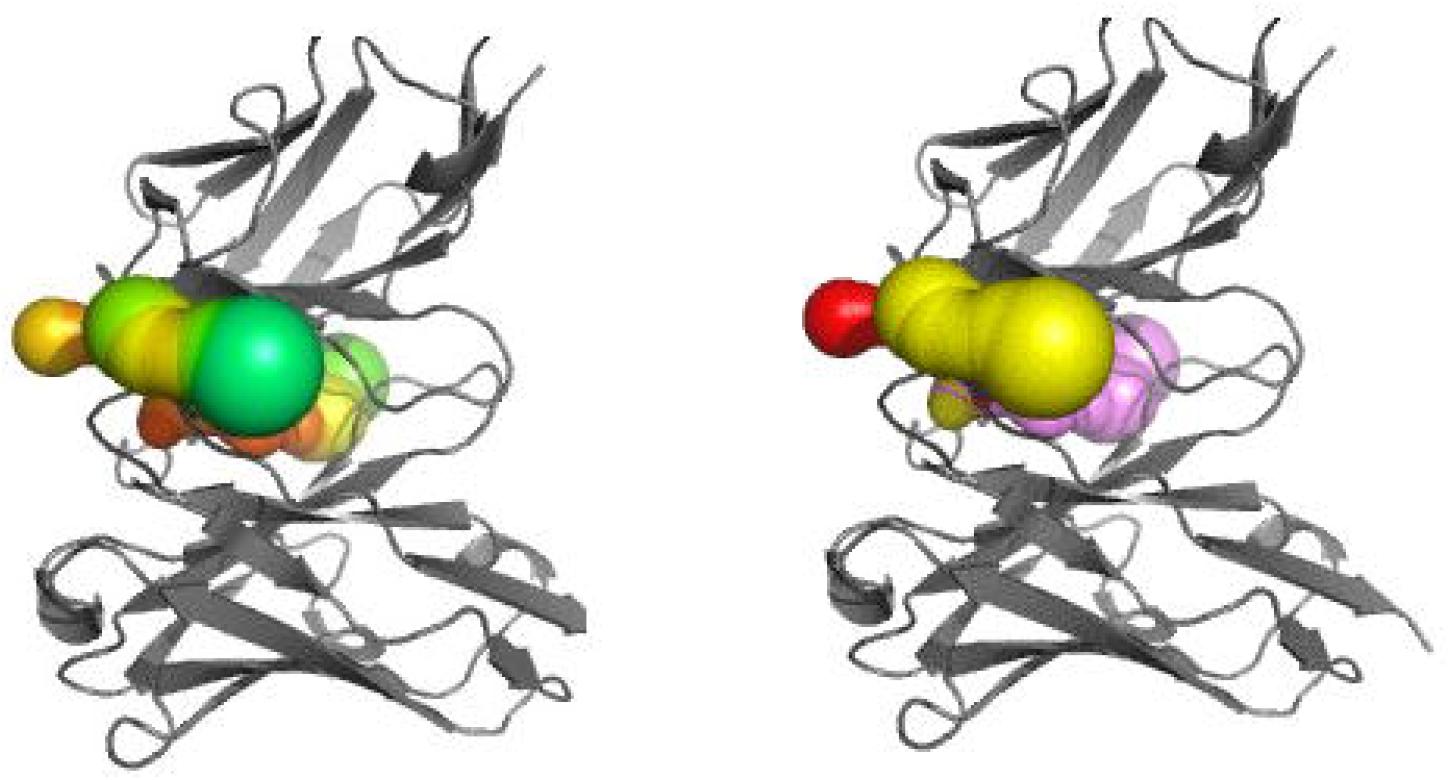
Tunnels in Fab chain

**Figure 17.**
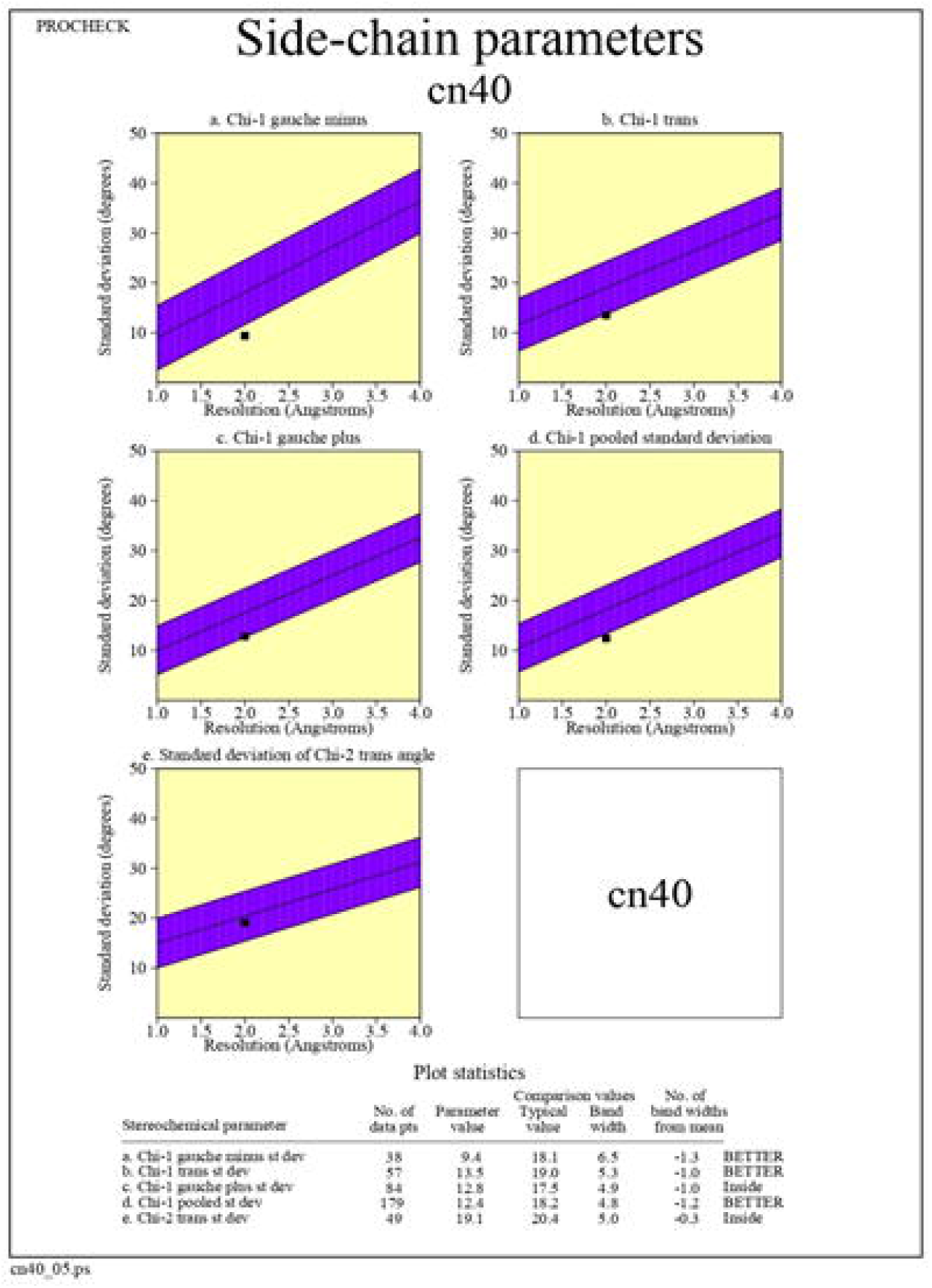
Side Chain parameters of Fab chain CR5840

**Figure 18.**
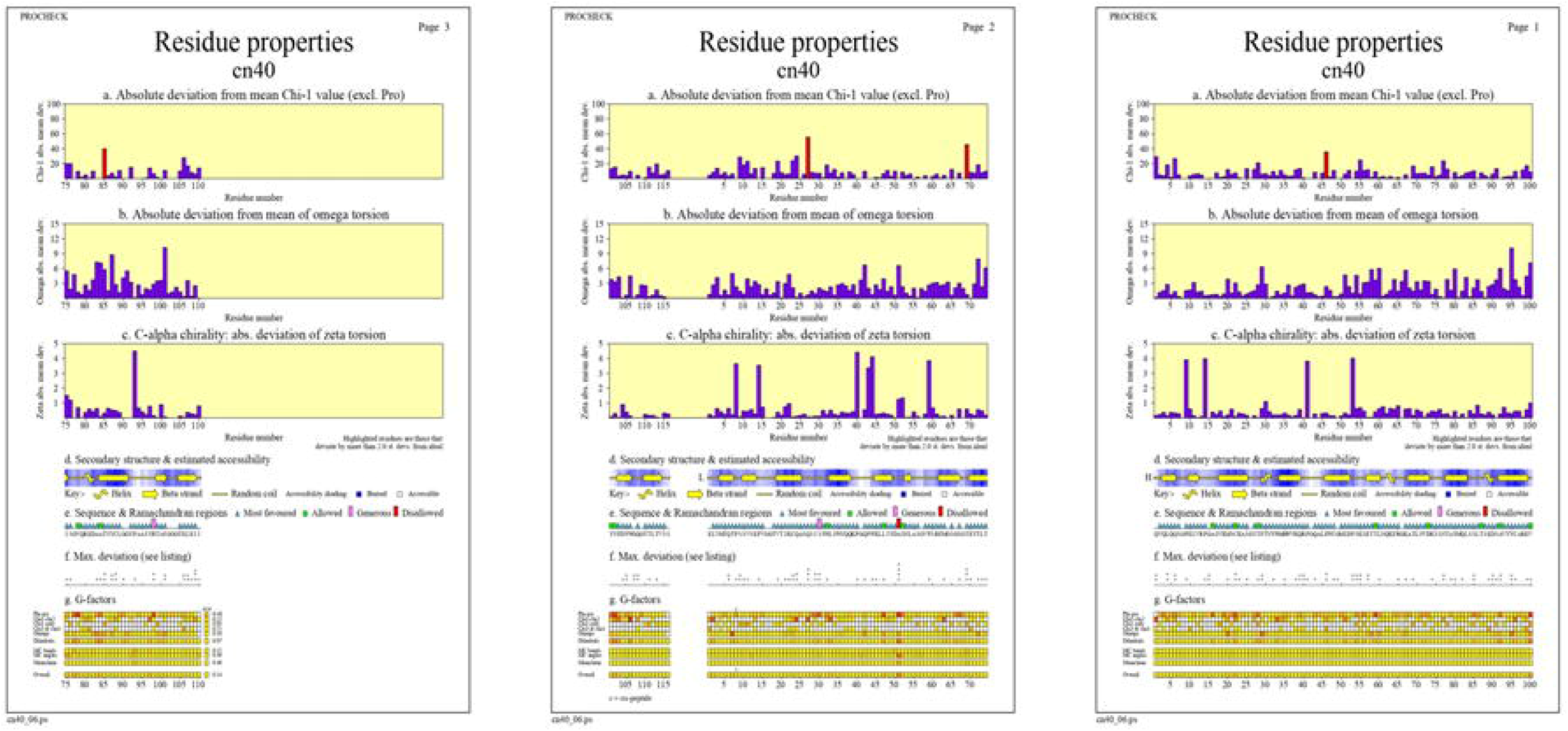
Residue properties of Fab chain CR5840

**Figure 19.**
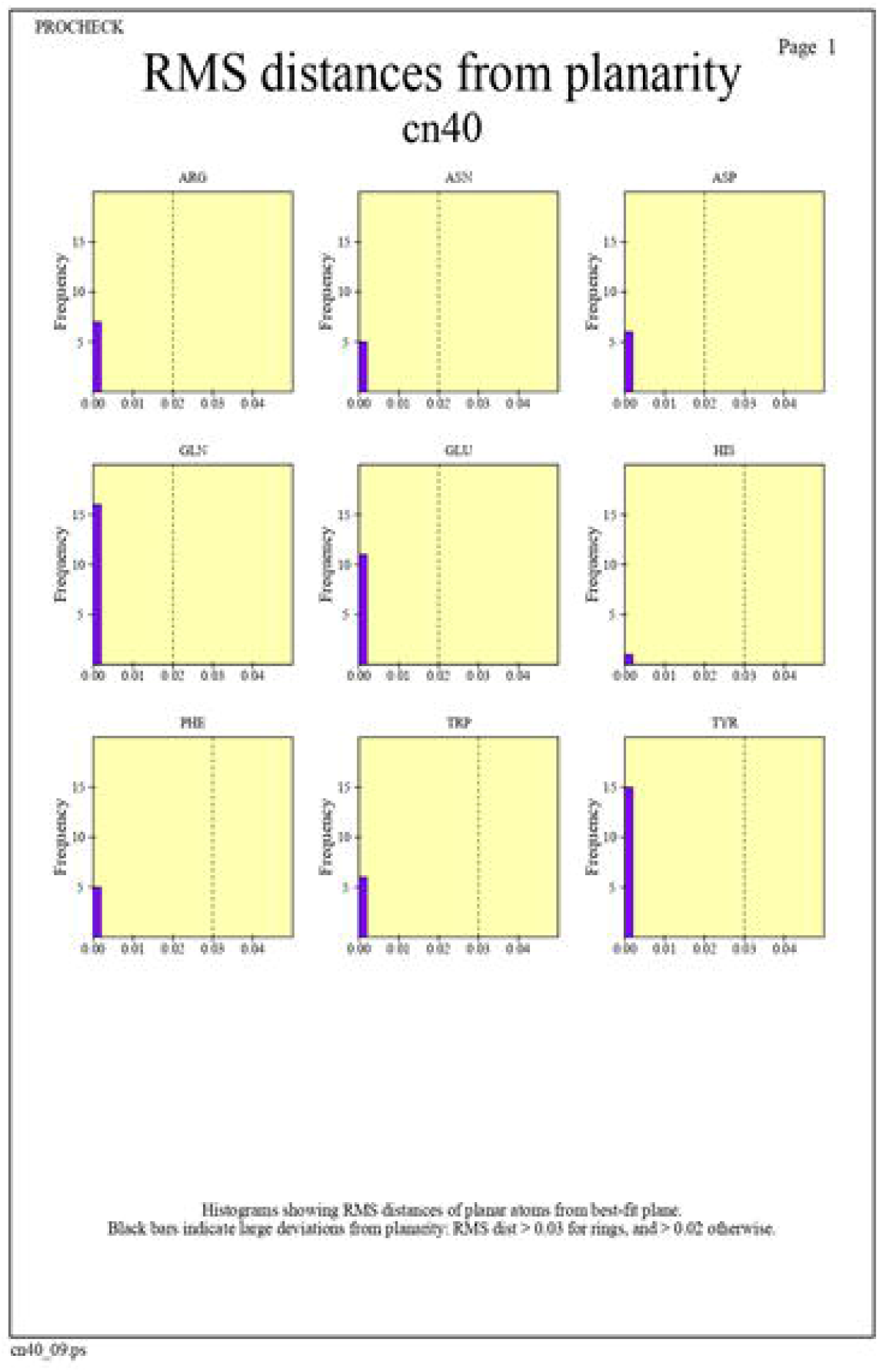
RMS distance of planarity of Fab chain CR5840

**Figure 20.**
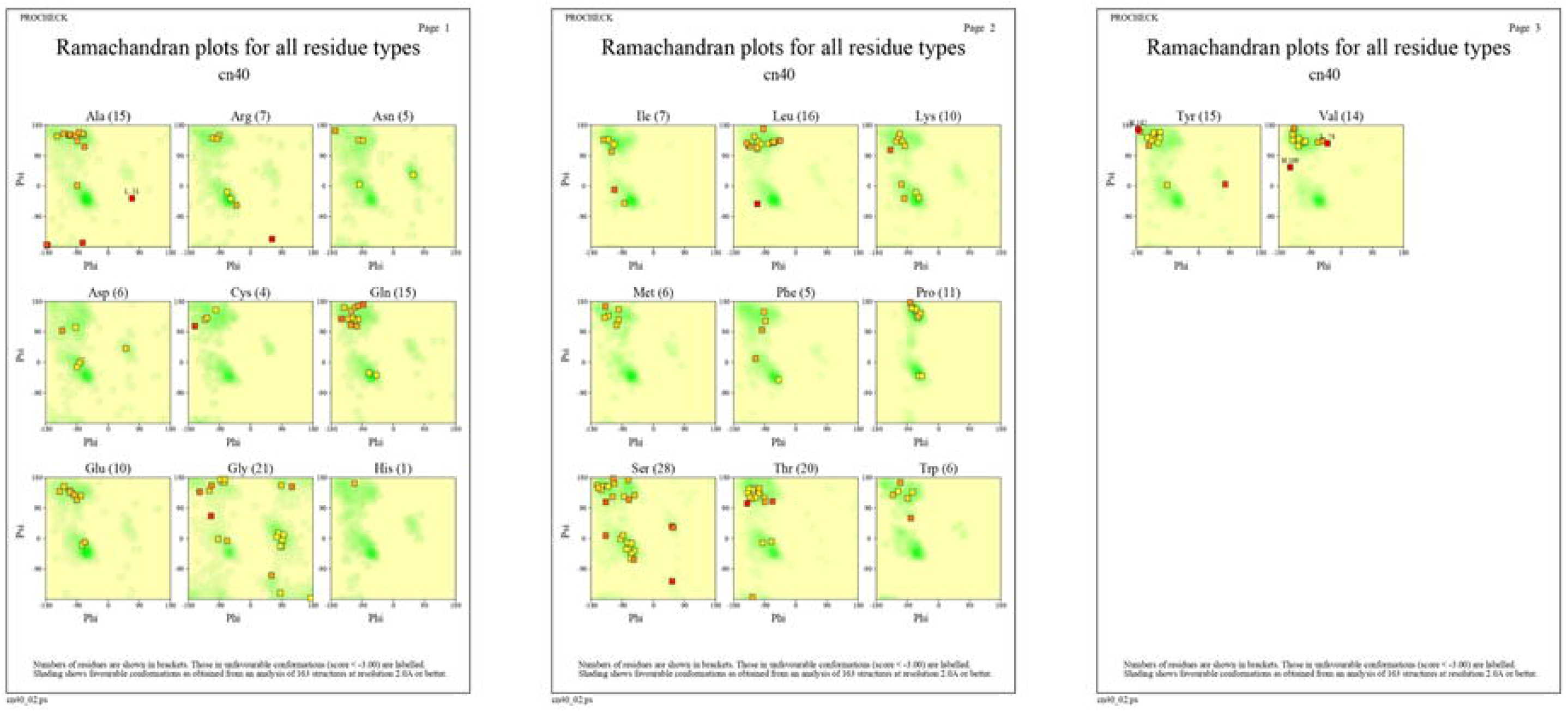
Ramachandran plots for all residue types of Fab chain CR5840

**Figure 21.**
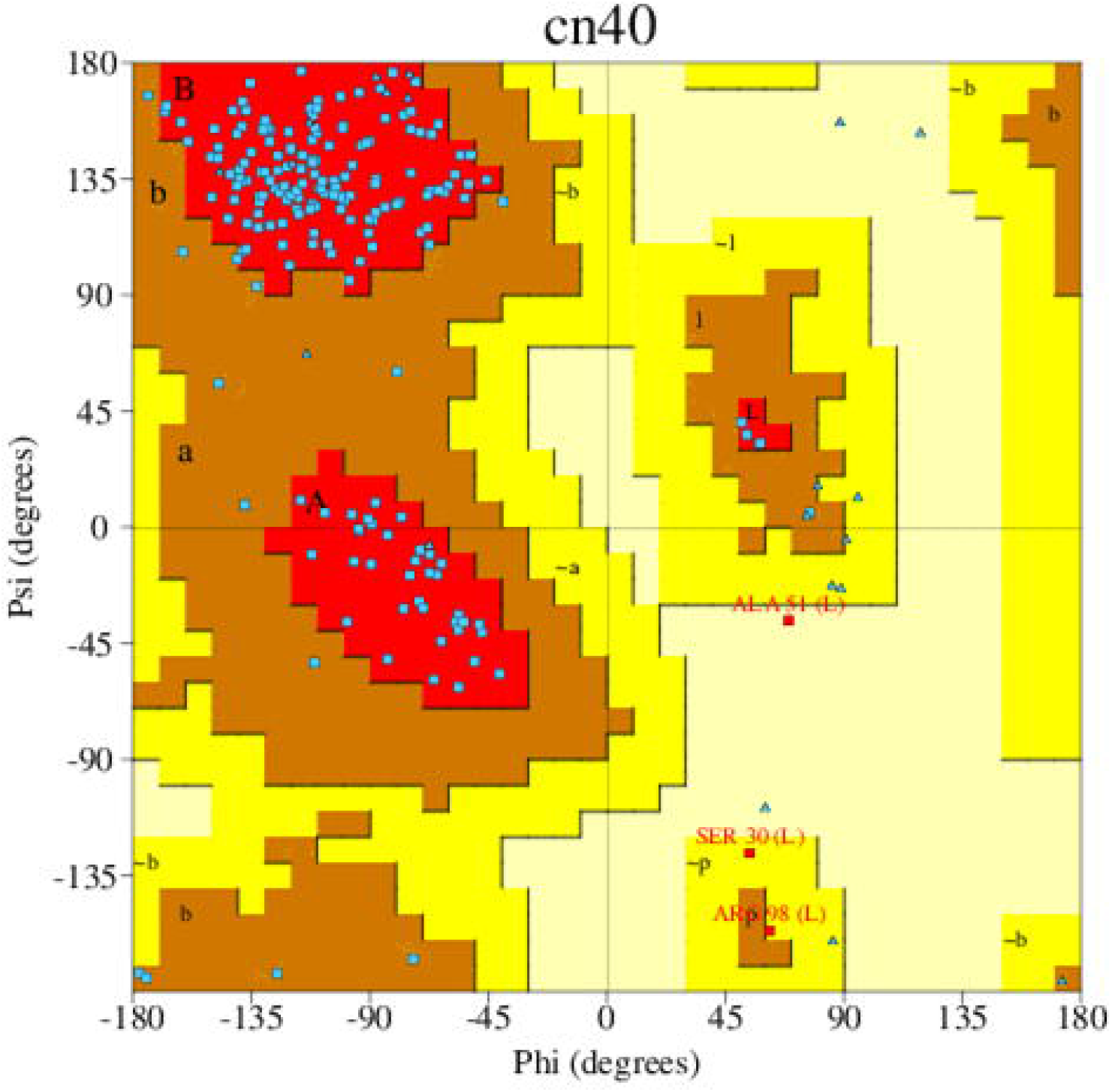
Ramachandran plots - Fab chain CR5840

### FRODOCK and Pydock protein-protein interaction

FRODOCK and Pydockweb tools are used to run protein-protein interaction CR5840 as ligand and Spike glycoprotein as Receptor (PDB ID: 6VSB) in comparison with available antibody CR3022. CR5840 had shown high binding affinity see figure 7 when compared to CR3022 against spike glycoprotein of SARS-CoV-2. Predicted rank order for top models of binding pockets shown in table 2 and 3.

**Table 3:**
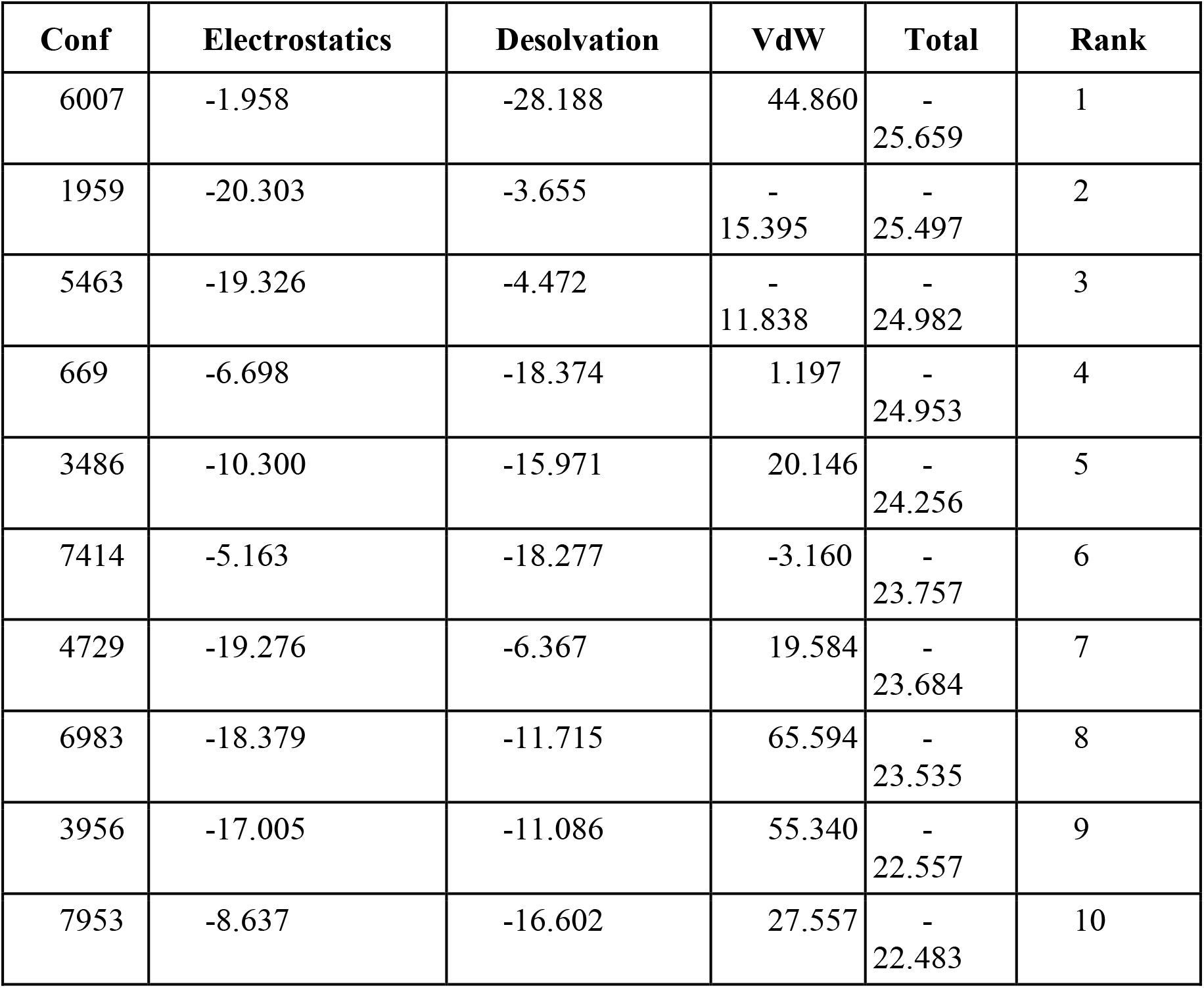
Top 10 energies and Rank predicted for *CR3022* interaction with SARS-CoV-2 Spike protein (PDB ID: 6VSB)

| Conf | Electrostatics | Desolvation | VdW | Total | Rank |
| --- | --- | --- | --- | --- | --- |
| 6007 | -1.958 | -28.188 | 44.860 | -<br>25.659 | 1 |
| 1959 | -20.303 | -3.655 | -<br>15.395 | -<br>25.497 | 2 |
| 5463 | -19.326 | -4.472 | -<br>11.838 | -<br>24.982 | 3 |
| 669 | -6.698 | -18.374 | 1.197 | -<br>24.953 | 4 |
| 3486 | -10.300 | -15.971 | 20.146 | -<br>24.256 | 5 |
| 7414 | -5.163 | -18.277 | -3.160 | -<br>23.757 | 6 |
| 4729 | -19.276 | -6.367 | 19.584 | -<br>23.684 | 7 |
| 6983 | -18.379 | -11.715 | 65.594 | -<br>23.535 | 8 |
| 3956 | -17.005 | -11.086 | 55.340 | -<br>22.557 | 9 |
| 7953 | -8.637 | -16.602 | 27.557 | -<br>22.483 | 10 |

## Discussion

There is an urgent necessary to address potential therapeutics for global pandemic created by SARS-CoV-2 infection. Viral antigens go through human innate immune system during initial or latent periods of incubation period and generate neutralizing antibodies against the antigens. Viral genome carries the coding for viral antigens which can be used to predict neutralizing human antibodies using different genomic and Highthroughput screening tools. Here used IgBLAST, I-TASSER and Molbiol tools to predict and design the neutralizing human antibodies against SARS-CoV-2 infection. Whole genome and spike protein genome sequence of SARS-CoV-2 were obtained from NCBI and RCSB database and IgBLAST studies carried out. Whole Genome sequence had shown top V gene match with IGHV1-38-4*01 and has VH type chain. Spike protein sequence had shown top V gene match with IGLV1-44*01, IGLV1-47*02 and has VL type chain. Thus we propose possibility for usage of VH and VL type of antibody germline as per sequence obtained for V-D-J chain for whole genome and V chain sequence for Spike protein mentioned in this research work. We propose cytotoxic T lymphocyte epitope peptide selective system as effective tool for the development of SARS-CoV-2 vaccine. Top predicted structural conformation of V-D Chain of neutralizing antibody shown in figure 5 with C-score=−2.03 (Read more about C-score), Estimated TM-score = 0.47±0.15, Estimated RMSD = 5.8±3.6Å. Fab fragment with Polyspecific activity IgG1 like properties CR5840 had shown highest affinity with spike protein PDN ID: 6VSB when compared with CR3302 an available Fab of antibody used in Rapid kit test for SARS-CoV-2.Thus we propose use of novel synthetic Fab active complex as therapeutic and diagnostic tool against SARS-CoV-2.

## Acknowledgement

This work was supported by the Bill and Melinda Gates Foundation Grand Challenge Exploration grant (GCE Round 4: OPP1015489) to SKJ.

