## Supplementary material for "Computational methods to develop potential neutralizing antibody Fab region against SARS-CoV-2 as therapeutic and diagnostic tool": Supp-1

- NCBI Home
- Sign in to NCBI
- Skip to Main Content
- Skip to Navigation
- About NCBI Accesskeys

U.S. National Library of Medicine

NCBI National Center for Biotechnology Information

- 
- My NCBI
- Sign in to NCBI
- Register
- Sign Out


IGBLAST» JOB ID: v6FUBaoHEysgiyS9faYH23yvAspj
Formatting Results

Database: imgt.Homo_sapiens.V.f.orf.p; imgt.Homo_sapiens.D.f.orf;
imgt.Homo_sapiens.J.f.orf
597 sequences; 157,743 total letters

**Query=** MN908947.3

Length=29903

| Sequences producingsignificantalignments: | Score (Bits) | E Value |
|---|---|---|
| IGHV1-38-4*01germlinegene | 29.9 | 4.1 |
| IGHV2-70*19germlinegene | 29.9 | 4.1 |
| IGLV1-50*01germlinegene | 29.9 | 4.1 |
| IGHD6-13*01germlinegene | 14.1 | 636 |
| IGHD2-15*01germlinegene | 14.1 | 636 |
| IGHD2-2*01germlinegene | 14.1 | 636 |

Domain classification requested: imgt

Note that your query represents the minus strand of a V gene and has been converted to the plus strand. The sequence positions refer to the converted sequence.

V-(D)-J rearrangement summary for query sequence (multiple equivalent top matches, **if** present, are separated by a comma):

| Top V gene match | Top D gene match | Top J gene match | Chain type | stop codon | V-J frame | Productive | Strand |
|---|---|---|---|---|---|---|---|
| IGHV1-38-4*01 | IGHD6-13*01 | N/A | VH | Yes | | No | - |

V-(D)-J junction details based on top germline gene matches:

| V region end | V-D junction* | D region | D-Jjunction* | J region start |
|---|---|---|---|---|
| TTTAT | CATTGTAGATGTCAAAAGCCCTGTATACGACATCAGTACTAGTGCCTGTGCCGCACGGTGTAAGACGGGCTGCACTTACACCGCAAACCCGTTTAAAAACGATTGT | GCATCAGCTG | | |

*: Overlapping nucleotides may exist at V-D-J junction (i.e, nucleotides that could be assigned
to either rearranging gene). Such nucleotides are indicated inside a parenthesis (i.e., (TACAT))
butarenotincludedundertheV,DorJgeneitself.

Alignment summary between query and top germline V gene hit:

| | from | to | length | matches | mismatches | gaps | identity(%) |
|---|---|---|---|---|---|---|---|
| FR1-IMGT | 16310 | 16319 | 10 | 8 | 2 | 0 | 80 |
| CDR1-IMGT | 16320 | 16343 | 24 | 17 | 7 | 0 | 70.8 |
| FR2-IMGT | 16344 | 16345 | 2 | 2 | 0 | 0 | 100 |
| Total | | | 36 | 27 | 9 | 0 | 75 |

### Alignments

```
                  lcl|Query_1_reversed  11549  GTAAATTGGTACCAACAGCTTCTCTAGTAGCATGACACCCCTCGACATCGAAGCCAATCCATGCACGTACATGTCTTATAGCTTCTTCGC  11638
V 90.9% (20/22)IGLV1-50*01             103    ...C.........G.......------------------------------------------------------------------  124

                  lcl|Query_1_reversed  11639  GGGTGATAAACATGTTAGGGTAACCATTAACTTGATAATTCATTTTAAAACCCATCATAGAGATGAGTCTTCTATAGGTCATGTCCTTAG  11728

                  lcl|Query_1_reversed  11729  GTATGCCAGGTATGTCAACACATAAACCTTCAGTTTTGAATTTAGTGTCAACACTGAGGTGTGTAGGTGCCTGTGTAGGATGTAACCCAG  11818

                  lcl|Query_1_reversed  11819  TGATTACCTTACTACAATCTTTAAAGAGTCCTGTTACATTTTCAGCTTGTAAAGTTGCCACATTCCTACGTGGAATTTCAAGACTTGTAA  11908

                  lcl|Query_1_reversed  11909  ATTGCAACTTGTCATAAAGGTCTCTATCAGACATTATGCAAAGTATGCCTACTTTTGCTCTGGTAATAGCAACATTAAATCTGTTTAC AT  11998
```

lcl|Query_1_reversed 11999 TACAAGAGTGAGCTGTTTCAGTGGTTTGAGTGAATATGACATAGTCATATTCTGAGCCCTGTGATGAATCAACAGTTTGAGTTGGTAGTC 12088

lcl|Query_1_reversed 12089 CCAAAATCTTTGAGGCTACAGCATTCTGTGAATTATAAGGTGAAATAAAGACAGCTTTTCTCCAAGCAGGGTTACGTGTAAGGAATTCTC 12178

lcl|Query_1_reversed 12179 TTACCACGCCTATTTGTGGCCTGTTAATTGCAGATGAAACATCATGCGTGATAACACCCTTATAAAACATTTTAAAGCATTGAGCTGATT 12268

lcl|Query_1_reversed 12269 TGTCTTTATGTGCTTTAAGCTTATTATCATAAACCAAAGCACTCACAGTGTCAACAATTTCAGCAGGACAACGCCGACAAGTTCCGAGGA 12358

lcl|Query_1_reversed 12359 ACATGTCTGGACCTATAGTTTTCATAAGTCTACACACTGAATTGAAATATTCTGGTTCTAGTGTGCCCTTAGTTAGCAATGTGCGTGGTG 12448

lcl|Query_1_reversed 12449 CAGGTAATTGAGCAGGGTCGCCAATGTACACATAGTGCTTAGCACGTAATCTGGCATTGACAACACTCAAATCATAATTTGTGGCCATTG 12538

lcl|Query_1_reversed 12539 AAATTTCATCAAAGACAACTATATCTGCTGTCGTCTCAGGCAATGCATTTACAGTACAAAAGACATACTGTTCTAATGTTGAATTCACTT 12628

lcl|Query_1_reversed 12629 TGAATTTATCAAAACACTCTACACGAGCACGTGCAGGTATAATTCTACTACATTTATCTATAGGCAAATATTTTAATGCCTTCTCACATA 12718

lcl|Query_1_reversed 12719 GTGCATCAACAGCGGCATGAGAGCAAGCTGTATACACTATGCGAGCAGAAGGGTAGTAGAGAGCTAGGCCAATAGCAAAATGACTCTTAC 12808

lcl|Query_1_reversed 12809 CAGTACCAGGTGGTCCCTGGAGTGTAGAATACTTTTGCATACCAACCTTTTGATAATTTGCAACATTGCTAGAAAACTCATCTGAGATAT 12898

lcl|Query_1_reversed 12899 TGAGTGTTGGGTATAAGCCAGTAATTCTAACATAGTGCTCTTGTGGCACTAGTGTAGGTGCACTTAATGGCATTACTGTATGTGATGTCA 12988

lcl|Query_1_reversed 12989 GCACAAAATAATCACCAACATTTAATTTGTAAGTTGTTGTACCTCGGTAAACAACAGCATCACCATAGTCACCTTTTTCAAAGGTGTACT 13078

lcl|Query_1_reversed 13079 CTCCTATTTGTACTTTACTGTTTTTAGTTACACGATAACCAGTAAAGACATAATTTCGGTTAAGTGGTGGTCTAGGTTTACCAACTTCCC 13168 lcl|Query_1_reversed 13169 ATGAAAGATGTAATTCTCTGTCAGACAGCACTTCACGTACAGTAGCAATACCATAAGACAGTTTAAATGTCTCCTCAGTAGCTTTGAGCG 13258

lcl|Query_1_reversed 13259 TTTCTGCTGCAAAAAGCTTGAGTCTTTCAGTACAGGTGTTAGCTAAAATGTAATCACCAGCATTTGTCCAGTCACATGTTGCAATTGCAT 13348

lcl|Query_1_reversed 13349 TAAAGTCAGTAACATTATCGCTACCAACACATGTATTTTTATATAAACCAAAAACTTGTCCATTAGCACACAATGGAAAACTAATGGGTG 13438

lcl|Query_1_reversed 13439 GTTTATGTGATTTACAATAATAGCTCATACCTCCTAAGTAAAGTTGAGTCACATCTGTGACATCACAACCTGGAGCATTGCAAACATACG 13528

lcl|Query_1_reversed 13529 GATTAACAGACAAGACTAATTTATGTGATGTTGATATGACATGGTCGTAACAGCATTTACAACATAAGAATGGTCTACGTATGCAAGCAC 13618

lcl|Query_1_reversed 13619 CACATCTTAATGAAGTCTGTGAATTGCAAAGAACACAAGCCCCAACAGCCTGTAAGACTGTATGCGGTGTGTACATAGCCTCATAAAACT 13708

lcl|Query_1_reversed 13709 CAGGTTCCCAATACCTTGAAGTGTTATCATTAGTAAGCATAACAGAATACATGTCTAACATGTGTCCTGTTAACTCATCATGTAGCTTTC 13798

lcl|Query_1_reversed 13799 TTATGTATTGTAAGTACAAATGAAAGACATCAGCATACTCCTGATTAGGATGTTTAGTAAGTGGGTAAGCATCTATAGCTAAAGACACGA 13888

lcl|Query_1_reversed 13889 ACCGTTCAATCATAAGTGTACCATCTGTTTTTACGATATCATCTACAAAACAGCCGGCCCCTAGGATTCTTGATGGATCTGGGTAAGGAA 13978

lcl|Query_1_reversed 13979 GGTACACATAATCATCACCCTGTTTAACTAGCATTGTATGTTGAGAGCAAAATTCATGAGGTCCTTTAGTAAGGTCAGTCTCAGTCCAAC 14068

lcl|Query_1_reversed 14069 ATTTTGCTTCAGACATAAAAACATTGTTTTGATAATAAAGAACTGACTTAAAGTTCTTTATGCTAGCCACTAGACCTTGAGATGCATAAG 14158

lcl|Query_1_reversed 14159 TGCTATTGAAACACACAACAGCATCGTCAGAGAGTATCATCATTGAGAAATGTTTACGCAAATATGCGTAAAACTCATTCACAAAGTCTG 14248

lcl|Query_1_reversed 14249 TGTCAACATCTCTATTTCTATAGAGACACTCATAAAGTCTGTGTTGTAAATTGCGGACATACTTATCGGCAATTTTGTTACCATCAGTAG 14338 lcl|Query_1_reversed 14339 ATAAAAGTGCATTAACATTGGCCGTGACAGCTTGACAAATGTTAAAAACACTATTAGCATAAGCAGTTGTGGCATCTCCTGATGAGGTTC 14428

lcl|Query_1_reversed 14429 CACCTGGTTTAACATATAGTGAACCGCCACACATGACCATTTCACTCAATACTTGAGCACACTCATTAGCTAATCTATAGAAACGGTGTG 14518

lcl|Query_1_reversed 14519 ACAAGCTACAACACGTTGTATGTTTGCGAGCAAGAACAAGTGAGGCCATAATTCTAAGCATGTTAGGCATGGCTCTATCACATTTAGGAT 14608

lcl|Query_1_reversed 14609 AATCCCAACCCATAAGGTGAGGGTTTTCTACATCACTATAAACAGTTTTTAACATGTTGTGCCAACCACCATAGAATTTGCTTGTTCCAA 14698

lcl|Query_1_reversed 14699 TTACTACAGTAGCTCCTCTAGTGGCGGCTATTGATTTCAATAATTTTTGATGAAACTGTCTATTGGTCATAGTACTACAGATAGAGACAC 14788

lcl|Query_1_reversed 14789 CAGCTACGGTGCGAGCTCTATTCTTTGCACTAATGGCATACTTAAGATTCATTTGAGTTATAGTAGGGATGACATTACGTTTTGTATATG 14878

lcl|Query_1_reversed 14879 CGAAAAGTGCATCTTGATCCTCATAACTCATTGAATCATAATAAAGTCTAGCCTTACCCCATTTATTAAATGGAAAACCAGCTGATTTGT 14968

lcl|Query_1_reversed 14969 CTAGGTTGTTGACGATGACTTGGTTAGCATTAATACAGCCACCATCGTAACAATCAAAGTACTTATCAACAACTTCAACTACAAATAGTA 15058

lcl|Query_1_reversed 15059 GTTGTCTGATATCACACATTGTTGGTAGATTATAACGATAGTAGTCATAATCGCTGATAGCAGCATTACCATCCTGAGCAAAGAAGAAGT 15148

lcl|Query_1_reversed 15149 GTTTTAATTCAACAGAACTTCCTTCCTTAAAGAAACCCTTAGACACAGCAAAGTCATAGAAGTCTTTGTTAAAATTACCGGGTTTGACAG 15238

lcl|Query_1_reversed 15239 TTTGAAAAGCAACATTGTTAGTAAGTGCAGCTACTGAAAAGCACGTAGTGCGTTTATCTAGTAATAGATTACCAGAAGCAGCGTGCATAG 15328

lcl|Query_1_reversed 15329 CAGGGTCAGCAGCATACACAAGTAATTCCTTAAAACTAAGTCTAGAGCTATGTAAGTTTACATCCTGATTATGTACAACACCTAGCTCTC 15418

lcl|Query_1_reversed 15419 TGAAGTGGTATCCAGTTGAAACTACAAATGGAACACCATCAACAAATATTTTTCTCACTAGTGGTCCAAAACTTGTAGGTGGGAACACTG 15508

```
lcl|Query_1_reversed  15509  TAGAGAATAAAACATTAAAGTTTGCACAATGCAGAATGCATCTGTCATCCAAACAGTTAACACAATTTGGGTGGTATGTCTGATCCCAAT  15598

lcl|Query_1_reversed  15599  ATTTAAAATAACGGTCAAAGAGTTTTAACCTCTCTTCCGTGAAGTCATATTTTAACAAATCCCACTTAATGTAAGGCTTTGTTAAGTCAG  15688

lcl|Query_1_reversed  15689  TGTCAACATGTGACTCTGCAGTTAAAGCCCTGGTCAAGGTTAATATAGGCATTAACAATGAATAATAAGAATCTACAACAGGAACTCCAC  15778

lcl|Query_1_reversed  15779  TACCTGGCGTGGTTTGTATGAAATCACCGAAATCATACCAGTTACCATTGAGATCTTGATTATCTAATGTCAGTACACCAACAATACCAG  15868

lcl|Query_1_reversed  15869  CATTTCGCATGGCATCACAGAATTGTACTGTTTTTAACAAAGCTTGGCGTACACGTTCACCTAAGTTGGCGTATACGCGTAATATATCTG  15958

lcl|Query_1_reversed  15959  GGTTTTCTACAAAATCATACCAGTCCTTTTTATTGAAATAATCATCATCACAACAATTGTATGTGACAAGTATTTCTTTTAATGTGTCAC  16048

lcl|Query_1_reversed  16049  AATTACCTTCATCAAAATGCCTTAAAGCATAGACGAGGTCTGCCATTGTGTATTTAGTAAGACGTTGACGTGATATATGTGGTACCATGT  16138

lcl|Query_1_reversed  16139  CACCGTCTATTCTAAACTTAAAGAAGTCATGTTTAGCAACAGCTGGACAATCCTTAAGTAAATTATAAATTGTTTCTTCATGTTGGTAGT  16228

                                                   <--------------------------------FR1-IMGT---------------------------------
                                                                                                                      S F *
  lcl|Query_1_reversed  16229  TAGAGAAAGTGTGTCTCTTAACTACAAAGTAAGAATCAATTAAATTGTCATCTTCGTCCTTTTCTTGGAAGCGACAACAATTAGTTTTTA  16318
V 75.0% (27/36)IGHV1-38-4*01  66  -------------------------------------------------------------------------------.....................C  74
                                                                                                                      S F S

                               ><-------CDR1-IMGT------><--------------------FR2-IMGT ------------------------------------>
                                                                    E F S K T S Y F I I V D V K S P V Y D I S T S A C A A R C K
  lcl|Query_1_reversed  16319  GGAATTTAGCAAAACCAGCTACTTTATCATTGTAGATGTCAAAAGCCCTGTATACGACATCAGTACTAGTGCCTGTGCCGCACGGTGTAA  16408
V 75.0% (27/36)IGHV1-38-4*01  75  T.GG....C..TC.........GG...---------------------------------------------------------------  101
                                G F T I T S Y G
D 100.0%(7/7)  IGHD2-15*01  6  ----------------------------.......-----------------------------------------------------  12
D 100.0%(7/7)  IGHD2-2*01   6  ----------------------------.......-----------------------------------------------------  12

                                                                    T G C T Y T A N P F K N D C A S A D * S M G S R S * S Q L Q
  lcl|Query_1_reversed  16409  GACGGGCTGCACTTACACCGCAAACCCGTTTAAAAACGATTGTGCATCAGCTGACTGAAGCATGGGTTCGCGGAGTTGATCACAACTACA  16498
D 90.0% (9/10)  IGHD6-13*01  8  ---------------------------------------------...G......-------------------------------------  17

                                                                    P *P F H I P Q T V Q T V F L S V K P T G S L A Q V V G I C
```

lcl|Query_1_reversed 16499 GCCATAACCTTTCCACATACCGCAGACGGTACAGACTGTGTTTTTAAGTGTAAAACCCACAGGGTCATTAGCACAAGTTGTAGGTATTTG 16588

T Y L P F K S Q N P L G F G * S M W Q R Q Y R Q H D A P P K
lcl|Query_1_reversed 16589 TACATACTTACCTTTTAAGTCACAAAATCCTTTAGGATTTGGATGATCTATGTGGCAACGGCAGTACAGACAACACGATGCACCACCAAA 16678

D S * S I L A S G V T V I A * P V P V C V H N I L T Q L V I
lcl|Query_1_reversed 16679 GGATTCTTGATCCATATTGGCTTCCGGTGTAACTGTTATTGCCTGACCAGTACCAGTGTGTGTACACAACATCTTAACACAATTAGTGAT 16768

G C P P L A R * S L * A L A A S T A K A Q K D N T V E L A G
lcl|Query_1_reversed 16769 TGGTTGTCCCCCACTAGCTAGATAATCTTTGTAAGCTTTAGCAGCATCTACAGCAAAAGCACAGAAAGATAATACAGTTGAATTGGCAGG 16858

T S V A L P A C R R T V A A K L P S T I P L F R L F N P L I
lcl|Query_1_reversed 16859 CACTTCTGTTGCATTACCAGCTTGTAGACGTACTGTGGCAGCTAAACTACCAAGTACCATACCTCTATTTAGGTTGTTTAATCCTTTAAT 16948

K Y K Y F T L G P L G V S V T N L Q G G S S S V * I V P V P
lcl|Query_1_reversed 16949 AAAGTATAAATACTTCACTTTAGGACCTTTAGGTGTGTCTGTAACAAACCTACAAGGTGGTTCCAGTTCTGTATAGATAGTACCAGTTCC 17038

S L L G N L A H F K S C K S D N S A S T N L P P F V V L * *
lcl|Query_1_reversed 17039 ATCACTCTTAGGGAATCTAGCCCATTTCAAATCCTGTAAATCGGATAACAGTGCAAGTACAAACCTACCTCCCTTTGTTGTGTTGTAGTA 17128

A N A L S S V Q A V C V V P A A Q D I C R S A T G L S S L F
lcl|Query_1_reversed 17129 AGCTAACGCATTGTCATCAGTGCAAGCAGTTTGTGTAGTACCGGCAGCACAAGACATCTGTCGTAGTGCAACAGGACTAAGCTCATTATT 17218

C N L T A E L A L K A V T I R G H A K L G E L S I L I S L S
lcl|Query_1_reversed 17219 CTGTAATTTGACAGCAGAATTGGCCCTTAAAGCTGTTACAATAAGAGGCCATGCTAAATTAGGTGAATTGTCCATACTAATTTCACTAAG 17308

* T I L L S A S T T C W I S H N A D A * V N V V P S H V F L
lcl|Query_1_reversed 17309 TTGAACAATTTTACTATCTGCATCTACAACCTGTTGGATTTCCCACAATGCTGATGCATAAGTAAATGTTGTACCATCACACGTATTTTT 17398

Y V L * S G M T T I S L A A V V R G I M F K G T Q P S L A L
lcl|Query_1_reversed 17399 ATATGTGTTATAGTCTGGTATGACAACCATTAGTTTGGCTGCTGTTGTAAGAGGTATTATGTTCAAGGGAACACAACCATCTCTTGCATT 17488

L I M L L S A S L S N F L S I V K S I V C I A L V T F A L L
lcl|Query_1_reversed 17489 GTTGATAATGTTGTTGAGTGCATCATTATCCAACTTTCTAAGCATAGTGAAAAGCATTGTCTGCATAGCACTAGTAACTTTTGCCCTCTT 17578

S S D L A C L Y I W V I A * S A I F S N L R C M A A S R S N
lcl|Query_1_reversed 17579 GTCCTCAGATCTAGCCTGTTTATACATTTGGGTCATAGCTTGATCAGCCATCTTTTCCAACTTACGTTGCATGGCTGCATCACGGTCAAA 17668

S D L A T F K D F F N F L R T T S E S P L A T A C S *A S *

```
lcl|Query_1_reversed  17669  TTCAGATTTAGCCACATTCAAAGACTTCTTCAACTTTTTAAGAACAACTTCAGAATCACCATTAGCAACAGCCTGCTCATAAGCTTCTTG  17758

                             A V A K A A Y D G R E L N S E A I A C K V A L L S S I S S Q
lcl|Query_1_reversed  17759  AGCAGTAGCAAAAGCTGCATATGATGGAAGGGAACTAAACTCTGAGGCTATAGCTTGTAAGGTTGCCCTGTTGTCCAGCATTTCTTCACA  17848

                             S L F M S T A P C M E S K T E S S E T I F S K A S V V S L A
lcl|Query_1_reversed  17849  AAGCTTGTTTATGTCTACAGCACCCTGCATGGAAAGCAAAACAGAAAGTAGTGAAACCATTTTTTCAAAGGCTTCAGTAGTATCTTTAGC  17938

                             K R M S L C N W T H * A H N L D D D S T L S C C K T E S K T
lcl|Query_1_reversed  17939  TAAGAGAATGTCATTGTGTAACTGGACACATTGAGCCCACAATTTAGATGATGATTCTACTCTGAGTTGTTGCAAAACTGAGAGTAAGAC  18028

                             T D V H F T S D I L D C T V A T L I Q G L P P T P N N L M L
lcl|Query_1_reversed  18029  TACTGATGTGCACTTTACATCTGACATTTTAGACTGTACAGTGGCTACTTTGATACAAGGTTTGCCACCAACACCCAACAATTTAATGTT  18118

                             S L K A S M L F L G G S S P C E F I Y L N S C V E T K * S *
lcl|Query_1_reversed  18119  GAGTTTGAAGGCATCTATGCTATTCTTGGGTGGGAGTAGTCCCTGTGAATTCATATATCTAAACTCCTGTGTAGAAACTAAGTAATCATA  18208

                             T P R V S L K * R L S K Q K R P K * Q V Q K * P K K Q * T S
lcl|Query_1_reversed  18209  AACACCAAGAGTCAGTCTAAAGTAGCGGTTGAGTAAACAAAAGAGGCCAAAGTAACAAGTACAAAAATAGCCTAAGAAACAATAAACTAG  18298

                             I I H * S V L P V M K K I G Q Y S T H I K T I P L A K N M T
lcl|Query_1_reversed  18299  CATTATACACTGAAGTGTATTACCAGTTATGAAGAAAATAGGGCAATACTCAACACACATAAAAACAATACCTCTGGCCAAAAACATGAC  18388

                             V V T T P E * L E V T E I I R A H M E M A * S K A L P * * T
lcl|Query_1_reversed  18389  AGTTGTAACTACACCTGAGTAGTTAGAAGTAACAGAGATTATAAGAGCCCACATGGAAATGGCTTGATCTAAAGCATTACCATAATAAAC  18478

                             L * T S V K T F I S V H T L L A P S S Y T V L A V I R I S N
lcl|Query_1_reversed  18479  TTTATAAACGAGTGTCAAGACATTCATAAGTGTCCACACTCTCCTAGCACCATCATCATACACAGTTCTTGCTGTCATAAGGATTAGTAA  18568

                             T T A D A Y I T Q S F S L K P D K L V S T I S N H V I I R I
lcl|Query_1_reversed  18569  CACTACAGCTGATGCATACATAACACAGTCTTTTAGCTTAAAACCAGACAAACTAGTATCAACCATATCCAACCATGTCATAATACGCAT  18658

                             T Q L A G I * T I L K * A T V A R E G N K N K Q R N A C L C
lcl|Query_1_reversed  18659  CACCCAACTAGCAGGCATATAGACCATATTAAAATAAGCTACAGTGGCAAGAGAAGGTAACAAAAACAAACAGAGAAATGCATGCTTATG  18748

                             L T N I I A K A D I A I I P I A K G K K A F S Y K K K N K D
lcl|Query_1_reversed  18749  TTTGACAAACATCATTGCAAAAGCAGACATAGCAATAATACCCATAGCAAAAGGTAAAAAGGCATTTTCATACAAAAAAAAGAACAAAGA  18838

                                                                   H * V L W T K T K S E V K I V S N N Q W C V P L I V L F T A
```

```
lcl|Query_1_reversed  18839  CCATTGAGTACTCTGGACTAAAACTAAAAGTGAAGTCAAAATTGTGAGTAACAACCAGTGGTGTGTACCCTTGATTGTTCTTTTCACTGC  18928

                             L W K V T P E H C L T T S K G V N S S S N K A L P N M V R P
lcl|Query_1_reversed  18929  ACTTTGGAAAGTAACACCTGAGCATTGTCTAACAACATCAAAAGGTGTAAATTCATCTTCTAATAAAGCACTACCCAATATGGTACGTCC  19018

                             F I P F C S N S F N E A H I S K T A I P V * A E R G P S M S
lcl|Query_1_reversed  19019  ATTCATACCATTTTGCAGTAATTCTTTTAATGAAGCACACATATCTAAAACGGCAATTCCAGTTTGAGCAGAAAGAGGTCCTAGTATGTC  19108

                             T W S C V R G S * L Y F I A T R L K S L R V V V N R L R N H
lcl|Query_1_reversed  19109  AACATGGTCTTGTGTTAGAGGTTCATAATTGTACTTCATAGCCACAAGGTTAAAGTCATTAAGAGTTGTGGTAAATCGATTGAGAAACCA  19198

                             L S P F I T A A Y N Q A K T L T V I V V S V P A A C A V C L
lcl|Query_1_reversed  19199  CCTGTCTCCATTTATAACAGCAGCGTACAACCAAGCTAAAACATTAACTGTAATAGTTGTGTCCGTACCAGCTGCTTGTGCTGTTTGCCT  19288

                             S T K G P * K L P S K S V P A * T P V G N S I W C M * Q K E
lcl|Query_1_reversed  19289  GTCAACAAAAGGTCCATAAAAGTTACCTTCTAAGTCTGTGCCAGCATGAACTCCAGTTGGTAATTCCATATGGTGCATGTAACAAAAAGA  19378

                             T Q S * S M L K P T L P H E P L R N E P L I V K L G L I A H
lcl|Query_1_reversed  19379  GACACAGTCATAATCTATGTTAAAACCAACACTACCACATGAACCATTAAGGAATGAACCCTTAATAGTGAAATTGGGCCTCATAGCACA  19468

                             W * T P D G E P L * Q A N T E K V C P G * M R T N L Y L G V
lcl|Query_1_reversed  19469  TTGGTAAACACCAGATGGTGAACCATTGTAACAAGCTAACACTGAAAAAGTCTGTCCTGGTTGAATGCGAACAAACTTATACTTAGGTGT  19558

                             L G L A V S T L S L S T Q F C I E C P I T L S * T L P A C T
lcl|Query_1_reversed  19559  CTTAGGATTGGCTGTATCAACCTTAAGCTTAAGTACACAATTTTGCATAGAATGTCCAATAACCCTGAGTTGAACATTACCAGCCTGTAC  19648

                             K K L * L D L R M S K S S * L G L S M S S E V Q I T C L G Q
lcl|Query_1_reversed  19649  CAAGAAATTATGATTAGACTTACGAATGAGTAAATCTTCATAATTAGGGTTAAGCATGTCTTCAGAGGTGCAGATCACATGTCTTGGACA  19738

                             * T T S S S Q R P L S V V V P Q V T C T I Q P S T L P D G N
lcl|Query_1_reversed  19739  GTAAACTACGTCATCAAGCCAAAGACCGTTAAGTGTAGTTGTACCACAAGTTACTTGTACCATACAACCCTCAACTTTACCAGATGGGAA  19828

                             A I F L K P L C K T A E V I E V C G G W * R T S E P E L L K
lcl|Query_1_reversed  19829  TGCCATTTTTCTAAAACCACTCTGCAAAACAGCTGAGGTGATAGAGGTTTGTGGTGGTTGGTAAAGAACATCAGAACCTGAGTTACTGAA  19918

                             S L R A F A R * Q Q A A S L * L V V S I A P L K Y L Y L L *
lcl|Query_1_reversed  19919  GTCATTGAGAGCCTTTGCGAGATGACAACAAGCAGCTTCTCTGTAGCTAGTTGTATCCATTGCTCCACTAAAATACTTGTACTTATTATA  20008

                                                                   R A K Y L L Y C V R G N S T S L R N F R Y I S L F N K K V H
```

```
lcl|Query_1_reversed  20009        AAGAGCTAAGTATCTATTATATTGCGTAAGAGGTAATAGCACATCACTACGCAACTTTAGATACATTTCTTTATTTAACAAAAAGGTGCA  20098

                              S A A S S K V L K E T P L K T T R L F R * L L K N Q * K C F
lcl|Query_1_reversed  20099   CAGCGCAGCTTCTTCAAAAGTACTAAAGGAAACACCATTAAAGACTACACGTCTCTTTAGGTAATTACTAAAGAACCAATAGAAATGCTT  20188

                              V E I Q M I * A I V I Q K G T K G V N I T I H * I C A K K E
lcl|Query_1_reversed  20189   TGTGGAAATACAAATGATATAAGCAATTGTTATCCAGAAAGGTACTAAAGGTGTGAACATAACCATCCACTGAATATGTGCTAAAAAAGA  20278

                              T S L V R * N V K Y K * I T E * T P G K N E * T G V K Q S T
lcl|Query_1_reversed  20279   AACATCATTAGTAAGATAAAATGTCAAGTACAAGTAAATAACAGAATAAACACCAGGTAAGAATGAGTAAACTGGTGTTAAACAGAGTAC  20368

                              V N D I R N S K V L K A T T * L Y S P K A L L N L I K * * A
lcl|Query_1_reversed  20369   AGTGAATGACATAAGGAATAGTAAAGTATTAAAGGCAACTACATGACTGTATTCACCAAAAGCTCTTCTAAACCTCATAAAATAGTAGGC  20458

                              R H V T T I A T I P P A T I D A D M S K A P I G * I S G V N
lcl|Query_1_reversed  20459   AAGGCATGTTACTACGATAGCTACAATACCACCAGCTACTATAGATGCTGATATGTCCAAAGCACCAATAGGTTGAATTAGTGGTGTAAA  20548

                              I L V S K F T A S T P Q K T P G K D L * * S L L S T H L P L
lcl|Query_1_reversed  20549   CATATTAGTAAGTAAATTTACAGCATCTACACCACAGAAAACTCCTGGTAAAGATCTGTAATAATCATTGTTAAGTACCCATCTACCACT  20638

                              V D T Q T P A S D L S Q V P C L Q Y S E S K V V T T L T E P
lcl|Query_1_reversed  20639   AGTAGATACACAAACACCAGCTTCTGATCTTTCACAAGTGCCGTGCCTACAGTACTCAGAATCAAAAGTTGTTACCACTCTAACAGAACC  20728

                              S R * V L G N * I I E P S M S T * R V S G R K L S * A T E P
lcl|Query_1_reversed  20729   TTCAAGGTAGGTGTTAGGAAATTGAATAATAGAGCCATCCATGAGCACATAACGTGTGTCAGGGCGTAAACTTTCATAAGCAACAGAACC  20818

                              S S T L V S * Q Y G T G L P E A S L K I V H S A A K T Q A D
lcl|Query_1_reversed  20819   TTCTAGTACATTGGTATCATAACAATATGGTACTGGCTTACCAGAAGCATCTTTAAAAATTGTACATTCAGCAGCCAAAACACAAGCTGA  20908

                              V A K S V Y S I S F D G V * Q M L P T A L K T L G K K C K K
lcl|Query_1_reversed  20909   TGTTGCAAAGTCAGTGTACTCTATAAGTTTTGATGGTGTGTAACAGATGTTACCAACTGCACTAAAAACTCTAGGTAAGAAATGCAAAAA  20998

                              S P L V V R N I V P G K P G T T K P T S L V M T A A I N G Q
lcl|Query_1_reversed  20999   GTCACCATTAGTTGTGCGTAATATCGTGCCAGGCAAACCAGGCACGACAAAACCCACTTCTCTTGTTATGACTGCAGCAATCAATGGGCA  21088

                              A L S L V * L P P R W L N H V S K S A C L L A K Q V S V D A
lcl|Query_1_reversed  21089   AGCTTTGTCATTAGTATAACTACCACCACGCTGGCTAAACCATGTGTCAAAATCAGCATGTTTGTTAGCAAAACAAGTATCTGTAGATGC  21178

                                                                                M S R V T P P S I A L Y P M I S L E K S V C L D M T * T G V
```

```
lcl|Query_1_reversed  21179     TATGTCACGAGTGACACCACCATCAATAGCCTTGTATCCTATGATTTCACTTGAAAAGTCAGTATGTTTAGACATGACATGAACAGGTGT  21268

                        I K * K I A A T K R N T S V T L I N C F N Q L L T I L P P L
lcl|Query_1_reversed  21269  TATTAAATAGAAAATAGCAGCAACAAAAAGGAACACAAGTGTAACTTTAATTAACTGCTTCAACCAATTATTAACAATTTTACCACCCTT  21358

                        S A I F V V T T L T T C L V V A H V N L K G K L F F L A A L
lcl|Query_1_reversed  21359  AAGTGCTATCTTTGTTGTTACAACATTAACAACTTGTCTAGTAGTTGCACATGTCAACTTAAAAGGTAAGTTATTCTTTTTAGCAGCACT  21448

                        R I C F R S C S D N D M K S L T F H I K A M L * L F A T C A
lcl|Query_1_reversed  21449  ACGTATTTGTTTTCGTAGTTGTTCAGACAATGACATGAAATCTTTAACGTTCCATATCAAAGCAATGTTGTGACTTTTTGCTACCTGCGC  21538

                        L I * R A L Q S I Q A P R S R G V M F S T L L * V S I * L L
lcl|Query_1_reversed  21539  ATTAATATGACGCGCACTACAGTCAATACAAGCACCAAGGTCACGGGGTGTCATGTTTTCAACTTTGTTATAGGTGAGCATATAGTTATT  21628

                        Q L S P V T S M S D * C D N L R H S T T S L V S T S E S T N
lcl|Query_1_reversed  21629  ACAACTATCGCCAGTAACTTCTATGTCAGATTGATGTGACAATTTAAGACATTCAACAACATCTTTAGTTTCTACATCTGAATCAACAAA  21718

                        P C R A A E I K V D K T L S K D T F F A S S A S A V A T S V
lcl|Query_1_reversed  21719  CCCTTGCCGAGCTGCTGAAATAAAAGTAGATAAGACATTGTCTAAGGACACATTCTTTGCAAGTTCAGCTTCTGCAGTTGCAACTAGTGT  21808

                        L S F S I G T L K V D E N V L T * A S N I L T A T S A L S P
lcl|Query_1_reversed  21809  TTTGAGTTTTTCCATTGGTACGTTAAAAGTTGATGAAAACGTATTAACGTAAGCATCAAACATTTTAACTGCAACTTCCGCACTATCACC  21898

                        T S D T N A * S S N S I G * H I S * L * * T D A D F A D D S
lcl|Query_1_reversed  21899  AACATCAGACACTAATGCCTGATCTAGTAACAGTATAGGTTGACACATAAGCTGACTGTAGTAAACAGACGCTGATTTTGCAGATGATTC  21988

                        S H F D L P S K T I T L I G N E P L V L L A L R L S K L T K
lcl|Query_1_reversed  21989  TTCACATTTTGATTTACCATCAAAAACTATAACATTAATAGGCAATGAACCTTTAGTGTTATTAGCTCTCAGGTTGTCTAAGTTAACAAA  22078

                        * E R E C L S * V F * P A L S K * R W M E P F F T V T L S T
lcl|Query_1_reversed  22079  ATGAGAGAGAGAATGTCTTTCATAAGTCTTTTGACCAGCTTTATCAAAGTAAAGATGGATGGAACCATTCTTCACTGTAACACTATCAAC  22168

                        M * E D W S V G F I G L L N C S D K S L A T S S L I N V L P
lcl|Query_1_reversed  22169  GATGTAAGAAGACTGGTCAGTAGGATTTATTGGTCTTTTAAACTGTAGTGACAAGTCTCTCGCAACTTCATCACTAATAAATGTACTACC  22258

                        A Q N V S Q L T Q F Q L C S L Q K P L P P L A * T * K D L L
lcl|Query_1_reversed  22259  AGCACAGAATGTATCACAATTAACACAATTCCAATTGTGTAGTTTGCAAAAGCCTTTACCTCCATTAGCATAGACATAAAAGGACCTTCT  22348

                                                                T P L T I V V H S T L V A L L R L * H I I Q V D E L Q P S T
```

```
lcl|Query_1_reversed  22349     AACACCATTAACAATAGTTGTACATTCGACTCTTGTTGCTCTATTACGTTTGTAACACATCATACAAGTTGATGAATTACAACCGTCTAC  22438

                            T C T * L F H T * * N D A K K M Y I L T I A E I G A I C T R
lcl|Query_1_reversed  22439  AACATGCACATAACTTTTCCATACATAATAAAATGATGCAAAGAAGATGTACATTCTAACCATAGCTGAAATCGGGGCCATTTGTACAAG  22528

                            L I I N H I S Q E L L I K C T A K * L K N N C M I A A N P S
lcl|Query_1_reversed  22529  ATTAATTATTAACCACATAAGCCAAGAATTACTAATAAAATGTACTGCAAAATAGCTGAAAAACAATTGCATGATTGCAGCCAATCCAAG  22618

                            T * K N L V K R I Y A K N H S A T K P K A V K S H L K D E M
lcl|Query_1_reversed  22619  TACATAGAAAAACCTAGTGAAAAGAATATATGCCAAAAACCACTCTGCAACTAAGCCAAAAGCAGTTAAATCCCATTTAAAAGATGAAAT  22708

                            V I C I V S K E G * V S K E S K P L R Q T L Q G I E P V Q *
lcl|Query_1_reversed  22709  GGTAATTTGTATAGTTTCTAAAGAAGGATAGGTGTCTAAAGAATCTAAACCACTAAGACAAACACTACAAGGTATAGAACCAGTACAGTA  22798

                            V A I V T L V E F K * P S L * P V Q * E G M P K L D I K T P
lcl|Query_1_reversed  22799  GGTTGCAATAGTGACATTAGTAGAGTTCAAATAGCCTTCTCTGTAACCAGTACAGTAAGAAGGCATGCCTAAATTAGACATTAAAACACC  22888

                            K A A V E * I K E P R Q T L N S K N Q I I I F I S L E K L G
lcl|Query_1_reversed  22889  TAAAGCAGCGGTTGAGTAGATTAAAGAACCTAGGCAAACACTTAATAGTAAAAACCAAATTATAATATTTATCAGTTTAGAAAAATTAGG  22978

                            D F K * L N E A S R Q N L P T L L T V F F A I V V G I D A L
lcl|Query_1_reversed  22979  TGACTTCAAATAATTAAATGAAGCCTCTAGACAAAATTTACCGACACTCTTAACAGTATTCTTTGCTATAGTAGTCGGCATAGATGCTTT  23068

                            I L E F V L L V K V H N C S N K V K K * G I * L V Q T R F K
lcl|Query_1_reversed  23069  AATTCTAGAATTTGTACTTCTAGTAAAAGTACACAATTGTAGCAATAAAGTAAAGAAATAAGGCATATAATTAGTACAAACACGGTTTAA  23158

                            H R V T M L V V V L T T L L R K G L A * L A I V S Q G T L L
lcl|Query_1_reversed  23159  ACACCGTGTAACTATGTTAGTAGTTGTACTAACAACTTTGTTAAGAAAAGGCTTAGCATAATTAGCTATAGTATCCCAAGGGACACTATT  23248

                            T A A K P * V A R V F K P N T L D N S L G F L I V R L E L S
lcl|Query_1_reversed  23249  AACAGCAGCTAAACCATGAGTAGCAAGGGTTTTCAAACCTAATACTCTAGATAATTCATTAGGTTTCTTAATAGTAAGACTAGAATTGTC  23338

                            T * A A I R S V W P T S S V I F K L L F A G L S I M S P T T
lcl|Query_1_reversed  23339  TACATAAGCAGCCATTAGATCTGTGTGGCCAACCTCTTCTGTAATTTTTAAACTATTATTTGCTGGTTTAAGTATAATGTCTCCTACAAC  23428

                            S V V F T L H S R T S F C M V G F S T T S S E T G F R S S Q
lcl|Query_1_reversed  23429  TTCGGTAGTTTTCACATTACACTCAAGAACGTCTTTCTGTATGGTAGGATTTTCCACTACTTCTTCAGAGACTGGTTTTAGATCTTCGCA  23518

                                                       A R L S I P C A S S D F S T S N E F D V S T G F V L Q R Q R
```

```
                 lcl|Query_1_reversed  23519  GGCAAGATTATCCATTCCCTGCGCGTCCTCTGACTTCAGTACATCAAACGAATTTGATGTTTCAACTGGTTTTGTGCTCCAAAGACAACG  23608

                                                I H Q V F G L Y V A L L V A L L T C Q T I G L C N N L A P F
                 lcl|Query_1_reversed  23609  TATACACCAGGTATTTGGTTTATACGTGGCTTTATTAGTTGCATTGTTAACATGCCAAACAATAGGTTTATGTAACAATTTAGCTCCTTT  23698
V 78.1% (25/32)IGHV2-70*19             199    -----------------------------------------------------------------------------------------.  199

                                                L K E G V * C L *S
                 lcl|Query_1_reversed  23699  CTTAAAAGAGGGTGTGTAGTGTTTATAATCA 23729
V 78.1% (25/32)IGHV2-70*19             198    ...C.G...T.TGC...........C. ........... 168
```

| Lambda | K | H |
|---|---|---|
| 1.10 | 0.333 | 0.549 |

| Gapped Lambda | K | H |
|---|---|---|
| 1.08 | 0.280 | 0.540 |

Effective search space used: 4073052391

---

Total queries = 1
Total identifiable CDR3 = 0
Total unique clonotypes = 0

Database: imgt.Homo_sapiens.V.f.orf.p
Posted date: Feb 10, 2020 12:47 PM
Number of letters in database: 155,489
Number of sequences in database: 531

Database: imgt.Homo_sapiens.D.f.orf
Posted date: May 17, 2012 12:56 PM
Number of letters in database: 828
Number of sequences in database: 34

Database: imgt.Homo_sapiens.J.f.orf
Posted date: May 17, 2012 12:56 PM
Number of letters indatabase: 1,426
Number of sequences in database:32

Matrix: blastn matrix 1 -1
Gap Penalties: Existence: 4, Extension: 1

BLAST is a registered trademark of the National Library of Medicine

Support center Mailing list YouTube

- National Library Of Medicine
- National Institutes Of Health
- U.S. Department of Health & Human Services
- USA.gov

### NCBI

*National Centerfor Biotechnology Information, U.S. National Library of Medicine 8600 Rockville Pike, Bethesda MD, 20894 USA*
Policies and Guidelines| Contact
