## Supplementary material for "Computational methods to develop potential neutralizing antibody Fab region against SARS-CoV-2 as therapeutic and diagnostic tool": Supp-2

- NCBI Home
- Sign in to NCBI
- Skip to Main Content
- Skip to Navigation
- About NCBI Accesskeys

U.S. National Library of Medicine

NCBI National Center for Biotechnology Information

- 
- My NCBI
- Sign in to
- NCBI Register
- Sign Out

IGBLAST» JOB ID: 1Mo-yQ3LtOeHR4Nx2mqgF9tjpQbE
Formatting Results

Database: imgt.Homo_sapiens.V.f.orf.p; imgt.Homo_sapiens.D.f.orf;
imgt.Homo_sapiens.J.f.orf
597 sequences; 157,743 total letters

**Query=** reverse translation of 6VSB:A|PDBID|CHAIN|SEQUENCE

Length=3864

| Sequences producing significant alignments: | Score (Bits) | E Value |
|---|---|---|
| IGLV1-44*01 germline gene | 28.8 | 4.6 |
| IGLV1-47*02 germline gene | 28.8 | 4.6 |
| IGLV3-22*01 germline gene | 28.8 | 4.6 |

Domain classification requested: imgt

Note that your query represents the minus strand of a V gene and has been converted to the plus strand. The sequence positions refer to the converted sequence. V-

(D)-J rearrangement summary for query sequence (multiple equivalent top matches, if present, are separated by a comma):

| Top V gene match | Top J gene match | Chain type | stop codon | V-J frame | Productive | Strand |
|---|---|---|---|---|---|---|

| | | | | | | |
|---|---|---|---|---|---|---|
| IGLV1-44*01,IGLV1-47*02 | N/A | VL | No | | | – |

V-(D)-J junction details based on top germline gene matches:

| V region end | V-Jjunction* | J regionstart |
|---|---|---|
| TGCGG | | |

*: Overlapping nucleotides may exist at V-D-J junction (i.e, nucleotides that could be assigned to either rearranging gene). Suchnucleotides are indicated inside a parenthesis (i.e., (TACAT)) butarenotincludedundertheV,DorJgeneitself.

Alignment summary between query and top germline V gene hit:

| | from | to | length | matches | mismatches | gaps | identity(%) |
|---|---|---|---|---|---|---|---|
| FR2-IMGT | 478 | 516 | 39 | 22 | 17 | 0 | 56.4 |
| CDR2-IMGT | 517 | 525 | 9 | 8 | 1 | 0 | 88.9 |
| FR3-IMGT | 526 | 531 | 6 | 5 | 1 | 0 | 83.3 |
| Total | | | 54 | 35 | 19 | 0 | 64.8 |

### Alignments

```
                                         <--------------FR2-IMGT---------------><CDR2-IM><----------------------------------------
                                          H H I A V A A H K G V I G G N N L R F I K V A L G H K P M G
             lcl|Query_1_reversed    478  CACCACATCGCAGTTGCCGCTCACAAAGGTGTTATCGGTGGTAATAATCTGCGGTTCATAAAAGTTGCGCTGGGTCACAAACCAATGGGT    567
V 64.8% (35/54) IGLV1-44*01          112  ..G..GC..C...GAA.G..C.C....CTCC.C...TA.A.........A....-------------------------------------    165
                                          Q Q L P G T A P K L L I Y S N N Q R
V 64.8% (35/54) IGLV1-47*02          112  ..G..GC..C...GAA.G..C.C....CTCC.C...TA.A.........A....-------------------------------------    165

                                         --------FR3-IMGT ----------------------------------------------------------------------->
                                          A V A H K H A F A R K M R F A IM A N R R R G G K V F F L R
             lcl|Query_1_reversed    568  GCCGTTGCTCACAAACACGCCTTCGCGCGGAAAATGCGCTTTGCCATCATGGCAAATCGCCGGCGCGGTGGTAAAGTTTTTTTCCTGCGC    657

                                          R H I G H M Q K H H A M R R A L R K A H Q M I A F A A K I H
             lcl|Query_1_reversed    658  CGGCACATAGGTCACATGCAGAAACACCACGCCATGCGGCGCGCTCTGCGGAAAGCTCATCAGATGATAGCCTTTGCCGCAAAAATCCAC    747

                                          A F A L A Q H A F A H F G R R Q V R A R A N F R R A N Q L L
             lcl|Query_1_reversed    748  GCGTTTGCTCTGGCCCAGCACGCATTCGCTCATTTTGGTCGCCGCCAGGTTCGCGCTCGCGCGAATTTCCGCCGCGCGAATCAGCTGCTG    837

                                          G H IG L Q A L Q A A G N Q A IN L H F R F R R IQ A A Q N
             lcl|Query_1_reversed    838  GGTCACATAGGTCTGCAGGCTCTGCAGGCGGCCGGTAATCAGGCGATCAATCTGCACTTCCGCTTCCGGCGGATCCAGGCGGCTCAGAAT    927

                                           IV Q H A A N R A K V A A Q L F H Q G V Q R L R V L V H H I
```

```
lcl|Query_1_reversed  928   ATCGTTCAGCACGCTGCTAATCGCGCCAAAGTTGCTGCTCAGCTGTTTCACCAGGGTGTTCAGCGCCTGCGCGTTCTGGTTCACCACATC  1017

                             L Q F A Q R A R G A A Q A I L N F A N R A V K L V R N Q F L
lcl|Query_1_reversed  1018  CTGCAGTTTGCCCAGCGCGCTCGCGGTGCTGCTCAGGCTATCCTGAATTTTGCCAATCGCGCTGTTAAACTGGTTCGCAATCAGTTTCTG  1107

                             V F I Q H V L G H A N A V K A I R H L H R K R N L Q R R A R
lcl|Query_1_reversed  1108  GTTTTCATACAGCACGTTCTGGGTCACGCCAATGCCGTTAAAGCGATACGCCATCTGCATCGCAAACGGAATCTGCAGCGCCGCGCCCGC  1197

                             A K G P A A G N G A R Q Q R A G I L R N H F I G Q Q R R Q H
lcl|Query_1_reversed  1198  GCCAAAGGTCCAGCCGCTGGTAATGGTGCCCGCCAGCAGCGCGCTGGTATACTGCGCAATCATTTCATCGGTCAGCAGCGGCGGCAGCAC  1287

                             G Q A V K F L R A N Q I A R R N I A Q A I A I L F N K A R I
lcl|Query_1_reversed  1288  GGTCAGGCCGTTAAATTTCTGCGCGCAAATCAGATCGCGCGCCGCAATATCGCCCAGGCAATCGCCATACTGTTTAATAAAGCCCGCATC  1377

                             R Q G H F V K Q Q I F N K A A F A R F A R I R Q N L A K V K
lcl|Query_1_reversed  1378  CGCCAGGGTCACTTTGTTAAACAGCAGATCTTCAATAAAGCTGCGTTTGCTCGGTTTGCTCGGATCCGGCAGAATCTGGCTAAAGTTAAA  1467

                             A A K I F N R R G F I N L F H L R K H F L G V F I L F H R N
lcl|Query_1_reversed  1468  GCCGCCAAAATCTTTAATCGGCGGGGTTTTATAAATCTGTTTCACCTGCGCAAACACTTCCTGGGTGTTTTTATCCTGTTCCACCGCAAT  1557

                             A G Q R A V Q L G A K A A I L Q Q Q V A A F G A I A A N I H
lcl|Query_1_reversed  1558  GCCGGTCAGCGCGCGGTTCAGCTGGGTGCAAAAGCTGCCATACTGCAGCAGCAGGTTGCTGCATTCGGTGCTATCGCCGCAAATATACAT  1647

                             G A I H A G F G H A H R Q N F G G H A N G K V G R N R N A V
lcl|Query_1_reversed  1648  GGTGCAATCCACGCTGGTTTTGGTCATGCTCACCGGCAGAATTTCGGTGGTCACGCTAATGGTAAAGTTGGTCGGAATCGCAATGCTGTT  1737

                             V A I R H A V F R A Q A H G I R N N A L A R H A A R A A R A
lcl|Query_1_reversed  1738  GTTGCTATACGCCACGCTGTTTTCCGCGCCCAGGCTCATGGTATACGCAATAATGCTCTGGCTCGCCACGCTGCTCGCGCTGCCCGGGCT  1827

                             V G L G L I A R A N A R A N R N I A F I A V V H M F R A N Q
lcl|Query_1_reversed  1828  GTTGGTCTGGGTCTGATAGCTCGCGCAAATGCCCGCGCCAATCGGAATATCGCATTCATAGCTGTTGTTCACATGTTCCGCGCCAATCAG  1917

                             A A R A G L K H V A A G A I H A P G R G Q L I R M N R H R H
lcl|Query_1_reversed  1918  GCAGCCCGCGCGGGTCTGAAACACGTTGCTGCCGGTGCTATACACGCGCCAGGTCGGGGTCAGCTGATCCGCATGAATCGCCACCGGCAC  2007

                             F G A V H I L I Q H R H L V A G V G A R G N H A H A A K A A
lcl|Query_1_reversed  2008  TTCGGTGCAGTTCACATCCTGATACAGCACCGCCACCTGGTTGCTGGTGTTGGTGCCCGGGGTAATCACGCTCACGCCGCCAAAGCTGCA  2097

                                                     R G N I Q N F Q G L R I A H R I G G I R N I A A K L L K R Q
```

lcl|Query_1_reversed 2098 CGGGGTAATATCCAGAATTTCCAGGGTCTGCGGATCGCGCACCGCATCGGTGGTATCCGCAATATCGCGGCCAAACTGCTGAAACGGCAG 2187

```
                                        K F F V A F G Q H A G A G Q
                   lcl|Query_1_reversed 2188 AAATTTTTTGTTGCTTTCGGTCAGCACGCCGGTGCCGGTCAGG 2230
V 78.6% (22/28) IGLV3-22*01             240  ----------------.........G..C.T.C..AT.......  213
```

| Lambda | K | H |
|---|---|---|
| 1.10 | 0.333 | 0.549 |

| Gapped Lambda | K | H |
|---|---|---|
| 1.08 | 0.280 | 0.540 |

Effective search space used: 530720504

---

Total queries = 1
Total identifiable CDR3 = 0
Total unique clonotypes = 0

Database: imgt.Homo_sapiens.V.f.orf.p
Posted date: Feb 10, 2020 12:47PM
Number of letters in database: 155,489
Number of sequences in database:531

Database: imgt.Homo_sapiens.D.f.orf
Posted date: May 17, 2012 12:56 PM
Number of letters in database: 828
Number of sequences in database: 34

Database: imgt.Homo_sapiens.J.f.orf
Posted date: May 17, 2012 12:56 PM
Number of letters in database:1,426
Number of sequences in database: 32

Matrix: blastn matrix 1 -1
Gap Penalties: Existence: 4, Extension: 1

BLAST is a registered trademark of the National Library ofMedicine
Support center Mailing list YouTube

- NATIONAL LIBRARY OF MEDICINE National Library Of Medicine
- NIH National Institutes Of Health
- U.S. Department of Health & Human Services
- USA.gov Government Made Easy USA.gov

### NCBI

*National Centerfor Biotechnology Information, U.S. National Library of Medicine 8600 Rockville Pike, Bethesda MD, 20894 USA*
Policies and Guidelines | Contact
