## Supplementary material for "Computational methods to develop potential neutralizing antibody Fab region against SARS-CoV-2 as therapeutic and diagnostic tool": Supp-3

### NetMHC 4.0 Server - prediction results

#### Technical University of Denmark - DTU

---

### NetMHC version 4.0

### Input is in FSA format

### Peptide length14
### Rank Threshold for Strong binding peptides 0.500
### Rank Threshold for Weakbindingpeptides 2.000

| pos | HLA | peptide | Core | Offset | I_pos | I_len | D_pos | D_len | iCore | Identity | 1-log50k(aff) | Affinity(nM) | %Rank | BindLevel |
|---|---|---|---|---|---|---|---|---|---|---|---|---|---|---|
| 0 | HLA-A0101 | HCRCQKPCIRHQYC | CRCCIRHQY | 1 | 0 | 0 | 3 | 3 | CRCQKPCIRHQY | vdchain | 0.047 | 30056.59 | 36.00 | |
| 1 | HLA-A0101 | CRCQKPCIRHQYCL | PCIRHQYCL | 5 | 0 | 0 | 0 | 0 | PCIRHQYCL | vdchain | 0.051 | 28883.99 | 32.00 | |
| 2 | HLA-A0101 | RCQKPCIRHQYCLC | PCIRHQYCL | 4 | 0 | 0 | 0 | 0 | PCIRHQYCL | vdchain | 0.042 | 31706.83 | 44.00 | |
| 3 | HLA-A0101 | CQKPCIRHQYCLCR | PCIRHQYCL | 3 | 0 | 0 | 0 | 0 | PCIRHQYCL | vdchain | 0.039 | 32883.22 | 50.00 | |
| 4 | HLA-A0101 | QKPCIRHQYCLCRT | PCIRHQYCL | 2 | 0 | 0 | 0 | 0 | PCIRHQYCL | vdchain | 0.035 | 34195.24 | 60.00 | |
| 5 | HLA-A0101 | KPCIRHQYCLCRTV | CIYCLCRTV | 2 | 0 | 0 | 2 | 3 | CIRHQYCLCRTV | vdchain | 0.054 | 27776.04 | 27.00 | |
| 6 | HLA-A0101 | PCIRHQYCLCRTVD | PCIRHQYCL | 0 | 0 | 0 | 0 | 0 | PCIRHQYCL | vdchain | 0.048 | 29669.84 | 35.00 | |
| 7 | HLA-A0101 | CIRHQYCLCRTVDG | YCLCRTVDG | 5 | 0 | 0 | 0 | 0 | YCLCRTVDG | vdchain | 0.048 | 29842.72 | 36.00 | |
| 8 | HLA-A0101 | IRHQYCLCRTVDGL | YLCRTVDGL | 4 | 0 | 0 | 1 | 1 | YCLCRTVDGL | vdchain | 0.076 | 21886.22 | 13.00 | |
| 9 | HLA-A0101 | RHQYCLCRTVDGLH | HQYCLVDGL | 1 | 0 | 0 | 5 | 3 | HQYCLCRTVDGL | vdchain | 0.063 | 25204.78 | 20.00 | |
| 10 | HLA-A0101 | HQYCLCRTVDGLHL | LCRTVDGLL | 4 | 0 | 0 | 8 | 1 | LCRTVDGLHL | vdchain | 0.068 | 24008.29 | 17.00 | |
| 11 | HLA-A0101 | QYCLCRTVDGLHLH | YLCRTVDGL | 1 | 0 | 0 | 1 | 1 | YCLCRTVDGL | vdchain | 0.058 | 26822.96 | 24.00 | |
| 12 | HLA-A0101 | YCLCRTVDGLHLHR | YLCRTVDGL | 0 | 0 | 0 | 1 | 1 | YCLCRTVDGL | vdchain | 0.062 | 25460.78 | 20.00 | |
| 13 | HLA-A0101 | CLCRTVDGLHLHRK | TVDGLHLRK | 4 | 0 | 0 | 7 | 1 | TVDGLHLHRK | vdchain | 0.049 | 29497.63 | 34.00 | |
| 14 | HLA-A0101 | LCRTVDGLHLHRKP | LCRTVDGLL | 0 | 0 | 0 | 8 | 1 | LCRTVDGLHL | vdchain | 0.040 | 32512.45 | 48.00 | |
| 15 | HLA-A0101 | CRTVDGLHLHRKPV | TVDGLHLPV | 2 | 0 | 0 | 7 | 3 | TVDGLHLHRKPV | vdchain | 0.101 | 16721.65 | 6.50 | |
| 16 | HLA-A0101 | RTVDGLHLHRKPVK | TVDGLHLPV | 1 | 0 | 0 | 7 | 3 | TVDGLHLHRKPV | vdchain | 0.060 | 26227.75 | 22.00 | |
| 17 | HLA-A0101 | TVDGLHLHRKPVKR | TVDGLHLPV | 0 | 0 | 0 | 7 | 3 | TVDGLHLHRKPV | vdchain | 0.048 | 29724.46 | 35.00 | |
| 18 | HLA-A0101 | VDGLHLHRKPVKRL | LHRKPVKRL | 5 | 0 | 0 | 0 | 0 | LHRKPVKRL | vdchain | 0.029 | 36598.32 | 75.00 | |
| 19 | HLA-A0101 | DGLHLHRKPVKRLC | LHRKPVKRL | 4 | 0 | 0 | 0 | 0 | LHRKPVKRL | vdchain | 0.025 | 38185.22 | 80.00 | |

Protein vdchain. Allele HLA-A0101. Number of high binders 0. Number of weak binders 0. Number of peptides 20

Link to Allele Frequencies in Worldwide PopulationsHLA-A0101

### Rank Threshold for Strong binding peptides 0.500
### Rank Threshold for Weakbindingpeptides 2.000

| pos | HLA | peptide | CoreOffset | | I_pos | I_len | D_pos | D_len | iCore | Identity | 1-log50k(aff) | Affinity(nM) | %Rank | BindLevel |
|---|---|---|---|---|---|---|---|---|---|---|---|---|---|---|
| 0 | HLA-A0201 | HCRCQKPCIRHQYC | CQKPCIQYC | 3 | 0 | 0 | 6 | 2 | CQKPCIRHQYC | vdchain | 0.041 | 32017.43 | 55.00 | |
| 1 | HLA-A0201 | CRCQKPCIRHQYCL | CQKPCIYCL | 2 | 0 | 0 | 6 | 3 | CQKPCIRHQYCL | vdchain | 0.063 | 25274.15 | 36.00 | |
| 2 | HLA-A0201 | RCQKPCIRHQYCLC | CQKPCIYCL | 1 | 0 | 0 | 6 | 3 | CQKPCIRHQYCL | vdchain | 0.055 | 27680.03 | 41.00 | |
| 3 | HLA-A0201 | CQKPCIRHQYCLCR | CQKPCIYCL | 0 | 0 | 0 | 6 | 3 | CQKPCIRHQYCL | vdchain | 0.052 | 28640.00 | 43.00 | |
| 4 | HLA-A0201 | QKPCIRHQYCLCRT | CIQYCLCRT | 3 | 0 | 0 | 2 | 2 | CIRHQYCLCRT | vdchain | 0.050 | 29042.87 | 44.00 | |
| 5 | HLA-A0201 | KPCIRHQYCLCRTV | KQYCLCRTV | 0 | 0 | 0 | 1 | 5 | KPCIRHQYCLCRTV | vdchain | 0.196 | 6003.43 | 11.00 | |
| 6 | HLA-A0201 | PCIRHQYCLCRTVD | CIYCLCRTV | 1 | 0 | 0 | 2 | 3 | CIRHQYCLCRTV | vdchain | 0.120 | 13596.01 | 19.00 | |
| 7 | HLA-A0201 | CIRHQYCLCRTVDG | CIYCLCRTV | 0 | 0 | 0 | 2 | 3 | CIRHQYCLCRTV | vdchain | 0.121 | 13515.05 | 19.00 | |
| 8 | HLA-A0201 | IRHQYCLCRTVDGL | YLCRTVDGL | 4 | 0 | 0 | 1 | 1 | YCLCRTVDGL | vdchain | 0.200 | 5768.28 | 11.00 | |
| 9 | HLA-A0201 | RHQYCLCRTVDGLH | YLCRTVDGL | 3 | 0 | 0 | 1 | 1 | YCLCRTVDGL | vdchain | 0.130 | 12267.80 | 18.00 | |
| 10 | HLA-A0201 | HQYCLCRTVDGLHL | HQYCLCLHL | 0 | 0 | 0 | 6 | 5 | HQYCLCRTVDGLHL | vdchain | 0.201 | 5701.33 | 11.00 | |
| 11 | HLA-A0201 | QYCLCRTVDGLHLH | CLCRTVLHL | 2 | 0 | 0 | 6 | 2 | CLCRTVDGLHL | vdchain | 0.132 | 11989.47 | 17.00 | |
| 12 | HLA-A0201 | YCLCRTVDGLHLHR | CLCRTVLHL | 1 | 0 | 0 | 6 | 2 | CLCRTVDGLHL | vdchain | 0.124 | 13092.32 | 19.00 | |
| 13 | HLA-A0201 | CLCRTVDGLHLHRK | CLCRTVLHL | 0 | 0 | 0 | 6 | 2 | CLCRTVDGLHL | vdchain | 0.121 | 13545.79 | 19.00 | |
| 14 | HLA-A0201 | LCRTVDGLHLHRKP | LCRTVGLHL | 0 | 0 | 0 | 5 | 1 | LCRTVDGLHL | vdchain | 0.040 | 32439.02 | 55.00 | |
| 15 | HLA-A0201 | CRTVDGLHLHRKPV | RTVDGLHPV | 1 | 0 | 0 | 7 | 4 | RTVDGLHLHRKPV | vdchain | 0.185 | 6747.66 | 12.00 | |
| 16 | HLA-A0201 | RTVDGLHLHRKPVK | RTVDGLHPV | 0 | 0 | 0 | 7 | 4 | RTVDGLHLHRKPV | vdchain | 0.113 | 14712.15 | 21.00 | |
| 17 | HLA-A0201 | TVDGLHLHRKPVKR | TVDGLHLPV | 0 | 0 | 0 | 7 | 3 | TVDGLHLHRKPV | vdchain | 0.070 | 23462.32 | 33.00 | |
| 18 | HLA-A0201 | VDGLHLHRKPVKRL | GLHLHVKRL | 2 | 0 | 0 | 5 | 3 | GLHLHRKPVKRL | vdchain | 0.070 | 23522.06 | 33.00 | |
| 19 | HLA-A0201 | DGLHLHRKPVKRLC | GLHLVKRLC | 1 | 0 | 0 | 4 | 4 | GLHLHRKPVKRLC | vdchain | 0.061 | 25950.27 | 38.00 | |

Protein vdchain. Allele HLA-A0201. Number of high binders 0. Number of weak binders 0. Number of peptides 20

Link to Allele Frequencies in Worldwide PopulationsHLA-A0201

### Rank Threshold for Strong binding peptides 0.500
### Rank Threshold for Weakbindingpeptides 2.000

| pos | HLA | peptide | CoreOffset | | I_pos | I_len | D_pos | D_len | iCore | Identity | 1-log50k(aff) | Affinity(nM) | %Rank | BindLevel |
|---|---|---|---|---|---|---|---|---|---|---|---|---|---|---|
| 0 | HLA-A0202 | HCRCQKPCIRHQYC | CQKPCIQYC | 3 | 0 | 0 | 6 | 2 | CQKPCIRHQYC | vdchain | 0.024 | 38690.93 | 70.00 | |
| 1 | HLA-A0202 | CRCQKPCIRHQYCL | CQKPCIYCL | 2 | 0 | 0 | 6 | 3 | CQKPCIRHQYCL | vdchain | 0.098 | 17343.94 | 30.00 | |
| 2 | HLA-A0202 | RCQKPCIRHQYCLC | CQKPCIYCL | 1 | 0 | 0 | 6 | 3 | CQKPCIRHQYCL | vdchain | 0.069 | 23691.92 | 39.00 | |
| 3 | HLA-A0202 | CQKPCIRHQYCLCR | CQKPCIYCL | 0 | 0 | 0 | 6 | 3 | CQKPCIRHQYCL | vdchain | 0.068 | 24082.16 | 40.00 | |
| 4 | HLA-A0202 | QKPCIRHQYCLCRT | CIQYCLCRT | 3 | 0 | 0 | 2 | 2 | CIRHQYCLCRT | vdchain | 0.036 | 33974.70 | 60.00 | |
| 5 | HLA-A0202 | KPCIRHQYCLCRTV | CIYCLCRTV | 2 | 0 | 0 | 2 | 3 | CIRHQYCLCRTV | vdchain | 0.167 | 8243.69 | 19.00 | |
| 6 | HLA-A0202 | PCIRHQYCLCRTVD | CIYCLCRTV | 1 | 0 | 0 | 2 | 3 | CIRHQYCLCRTV | vdchain | 0.128 | 12549.05 | 24.00 | |
| 7 | HLA-A0202 | CIRHQYCLCRTVDG | CIYCLCRTV | 0 | 0 | 0 | 2 | 3 | CIRHQYCLCRTV | vdchain | 0.125 | 12896.05 | 24.00 | |
| 8 | HLA-A0202 | IRHQYCLCRTVDGL | YLCRTVDGL | 4 | 0 | 0 | 1 | 1 | YCLCRTVDGL | vdchain | 0.222 | 4515.09 | 13.00 | |
| 9 | HLA-A0202 | RHQYCLCRTVDGLH | YLCRTVDGL | 3 | 0 | 0 | 1 | 1 | YCLCRTVDGL | vdchain | 0.142 | 10779.18 | 22.00 | |
| 10 | HLA-A0202 | HQYCLCRTVDGLHL | HQYCLCLHL | 0 | 0 | 0 | 6 | 5 | HQYCLCRTVDGLHL | vdchain | 0.207 | 5348.34 | 15.00 | |
| 11 | HLA-A0202 | QYCLCRTVDGLHLH | YLCRTVDGL | 1 | 0 | 0 | 1 | 1 | YCLCRTVDGL | vdchain | 0.120 | 13692.70 | 25.00 | |
| 12 | HLA-A0202 | YCLCRTVDGLHLHR | YLCRTVDGL | 0 | 0 | 0 | 1 | 1 | YCLCRTVDGL | vdchain | 0.125 | 12885.17 | 24.00 | |
| 13 | HLA-A0202 | CLCRTVDGLHLHRK | CLCRTVLHL | 0 | 0 | 0 | 6 | 2 | CLCRTVDGLHL | vdchain | 0.096 | 17613.61 | 30.00 | |
| 14 | HLA-A0202 | LCRTVDGLHLHRKP | LCRTVDLHL | 0 | 0 | 0 | 6 | 1 | LCRTVDGLHL | vdchain | 0.032 | 35548.86 | 65.00 | |
| 15 | HLA-A0202 | CRTVDGLHLHRKPV | RTVDGLHLV | 1 | 0 | 0 | 8 | 4 | RTVDGLHLHRKPV | vdchain | 0.135 | 11601.78 | 23.00 | |
| 16 | HLA-A0202 | RTVDGLHLHRKPVK | RTVDGLHLV | 0 | 0 | 0 | 8 | 4 | RTVDGLHLHRKPV | vdchain | 0.089 | 19042.47 | 32.00 | |
| 17 | HLA-A0202 | TVDGLHLHRKPVKR | TVDGLHLPV | 0 | 0 | 0 | 7 | 3 | TVDGLHLHRKPV | vdchain | 0.037 | 33461.05 | 60.00 | |

| | | | | | | | | | | | | |
|---|---|---|---|---|---|---|---|---|---|---|---|---|
| 18 | HLA-A0202VDGLHLHRKPVKRL | GLHLHVKRL | 2 | 0 | 0 | 5 | 3 | GLHLHRKPVKRL | vdchain | 0.095 | 17971.87 | 31.00 |
| 19 | HLA-A0202DGLHLHRKPVKRLC | GLHLHVKRL | 1 | 0 | 0 | 5 | 3 | GLHLHRKPVKRL | vdchain | 0.067 | 24267.09 | 40.00 |

---

Protein vdchain. Allele HLA-A0202. Number of high binders 0. Number of weak binders 0. Number of peptides 20

Link to Allele Frequencies in Worldwide Populations HLA-A0202

---

### Rank Threshold for Strong binding peptides 0.500
### Rank Threshold for Weakbindingpeptides 2.000

---

| pos | HLA | peptide | CoreOffset | | I_pos | I_len | D_pos | D_len | iCore | Identity | 1-log50k(aff) | Affinity(nM) | %Rank | BindLevel |
|---|---|---|---|---|---|---|---|---|---|---|---|---|---|---|
| 0 | HLA-A0203 | HCRCQKPCIRHQYC | CQKPCIRYC | 3 | 0 | 0 | 7 | 2 | CQKPCIRHQYC | vdchain | 0.028 | 36905.30 | 75.00 | |
| 1 | HLA-A0203 | CRCQKPCIRHQYCL | CQKPCIRHL | 2 | 0 | 0 | 8 | 3 | CQKPCIRHQYCL | vdchain | 0.071 | 23223.14 | 41.00 | |
| 2 | HLA-A0203 | RCQKPCIRHQYCLC | CQKPCIYCL | 1 | 0 | 0 | 6 | 3 | CQKPCIRHQYCL | vdchain | 0.048 | 29758.58 | 55.00 | |
| 3 | HLA-A0203 | CQKPCIRHQYCLCR | CQKPCIYCL | 0 | 0 | 0 | 6 | 3 | CQKPCIRHQYCL | vdchain | 0.046 | 30320.85 | 60.00 | |
| 4 | HLA-A0203 | QKPCIRHQYCLCRT | CIRYCLCRT | 3 | 0 | 0 | 3 | 2 | CIRHQYCLCRT | vdchain | 0.068 | 23939.02 | 43.00 | |
| 5 | HLA-A0203 | KPCIRHQYCLCRTV | KQYCLCRTV | 0 | 0 | 0 | 1 | 5 | KPCIRHQYCLCRTV | vdchain | 0.281 | 2392.70 | 9.00 | |
| 6 | HLA-A0203 | PCIRHQYCLCRTVD | CIYCLCRTV | 1 | 0 | 0 | 2 | 3 | CIRHQYCLCRTV | vdchain | 0.168 | 8090.73 | 18.00 | |
| 7 | HLA-A0203 | CIRHQYCLCRTVDG | CIYCLCRTV | 0 | 0 | 0 | 2 | 3 | CIRHQYCLCRTV | vdchain | 0.162 | 8633.12 | 19.00 | |
| 8 | HLA-A0203 | IRHQYCLCRTVDGL | YLCRTVDGL | 4 | 0 | 0 | 1 | 1 | YCLCRTVDGL | vdchain | 0.140 | 11037.29 | 22.00 | |
| 9 | HLA-A0203 | RHQYCLCRTVDGLH | RQYCLCRTV | 0 | 0 | 0 | 1 | 1 | RHQYCLCRTV | vdchain | 0.110 | 15247.28 | 28.00 | |
| 10 | HLA-A0203 | HQYCLCRTVDGLHL | HQYCLCLHL | 0 | 0 | 0 | 6 | 5 | HQYCLCRTVDGLHL | vdchain | 0.170 | 7945.59 | 18.00 | |
| 11 | HLA-A0203 | QYCLCRTVDGLHLH | CLCRTVLHL | 2 | 0 | 0 | 6 | 2 | CLCRTVDGLHL | vdchain | 0.096 | 17684.84 | 32.00 | |
| 12 | HLA-A0203 | YCLCRTVDGLHLHR | CLCRTVLHL | 1 | 0 | 0 | 6 | 2 | CLCRTVDGLHL | vdchain | 0.088 | 19290.07 | 34.00 | |
| 13 | HLA-A0203 | CLCRTVDGLHLHRK | CLCRTVLHL | 0 | 0 | 0 | 6 | 2 | CLCRTVDGLHL | vdchain | 0.092 | 18453.24 | 33.00 | |
| 14 | HLA-A0203 | LCRTVDGLHLHRKP | LCRTVGLHL | 0 | 0 | 0 | 5 | 1 | LCRTVDGLHL | vdchain | 0.037 | 33681.91 | 65.00 | |
| 15 | HLA-A0203 | CRTVDGLHLHRKPV | RTVDGLHPV | 1 | 0 | 0 | 7 | 4 | RTVDGLHLHRKPV | vdchain | 0.290 | 2161.81 | 8.50 | |
| 16 | HLA-A0203 | RTVDGLHLHRKPVK | RTVDGLHPV | 0 | 0 | 0 | 7 | 4 | RTVDGLHLHRKPV | vdchain | 0.176 | 7461.17 | 17.00 | |
| 17 | HLA-A0203 | TVDGLHLHRKPVKR | TLHLHRKPV | 0 | 0 | 0 | 1 | 3 | TVDGLHLHRKPV | vdchain | 0.100 | 17024.33 | 31.00 | |
| 18 | HLA-A0203 | VDGLHLHRKPVKRL | GLHLHVKRL | 2 | 0 | 0 | 5 | 3 | GLHLHRKPVKRL | vdchain | 0.136 | 11441.96 | 23.00 | |
| 19 | HLA-A0203 | DGLHLHRKPVKRLC | GLHLHVKRL | 1 | 0 | 0 | 5 | 3 | GLHLHRKPVKRL | vdchain | 0.086 | 19762.01 | 35.00 | |

---

Protein vdchain. Allele HLA-A0203. Number of high binders 0. Number of weak binders 0. Number of peptides 20

Link to Allele Frequencies in Worldwide Populations HLA-A0203

---

### Rank Threshold for Strong binding peptides 0.500
### Rank Threshold for Weakbindingpeptides 2.000

---

| pos | HLA | peptide | CoreOffset | | I_pos | I_len | D_pos | D_len | iCore | Identity | 1-log50k(aff) | Affinity(nM) | %Rank | BindLevel |
|---|---|---|---|---|---|---|---|---|---|---|---|---|---|---|
| 0 | HLA-A0205 | HCRCQKPCIRHQYC | RCQKPCIYC | 2 | 0 | 0 | 7 | 3 | RCQKPCIRHQYC | vdchain | 0.011 | 44559.96 | 43.00 | |
| 1 | HLA-A0205 | CRCQKPCIRHQYCL | RCQKPCIYC | 1 | 0 | 0 | 7 | 3 | RCQKPCIRHQYC | vdchain | 0.011 | 44629.91 | 43.00 | |
| 2 | HLA-A0205 | RCQKPCIRHQYCLC | RCQKPCIYC | 0 | 0 | 0 | 7 | 3 | RCQKPCIRHQYC | vdchain | 0.010 | 44757.60 | 44.00 | |
| 3 | HLA-A0205 | CQKPCIRHQYCLCR | KCIRHQYCL | 2 | 0 | 0 | 1 | 1 | KPCIRHQYCL | vdchain | 0.008 | 45964.89 | 55.00 | |
| 4 | HLA-A0205 | QKPCIRHQYCLCRT | QKPRHQYCL | 0 | 0 | 0 | 3 | 2 | QKPCIRHQYCL | vdchain | 0.008 | 45875.46 | 55.00 | |
| 5 | HLA-A0205 | KPCIRHQYCLCRTV | IRHQYCLTV | 3 | 0 | 0 | 7 | 2 | IRHQYCLCRTV | vdchain | 0.053 | 28251.89 | 9.50 | |
| 6 | HLA-A0205 | PCIRHQYCLCRTVD | IRHQYCLTV | 2 | 0 | 0 | 7 | 2 | IRHQYCLCRTV | vdchain | 0.047 | 29949.80 | 11.00 | |

| | | | | | | | | | | | | | |
|---|---|---|---|---|---|---|---|---|---|---|---|---|---|
| 7 | HLA-A0205 | CIRHQYCLCRTVDG | IRHQYCLTV | 1 | 0 | 0 | 7 | 2 | IRHQYCLCRTV | vdchain | 0.039 | 32724.55 | 13.00 |
| 8 | HLA-A0205 | IRHQYCLCRTVDGL | IRHQYCLTV | 0 | 0 | 0 | 7 | 2 | IRHQYCLCRTV | vdchain | 0.047 | 30047.18 | 11.00 |
| 9 | HLA-A0205 | RHQYCLCRTVDGLH | YLCRTVDGL | 3 | 0 | 0 | 1 | 1 | YCLCRTVDGL | vdchain | 0.025 | 38353.73 | 21.00 |
| 10 | HLA-A0205 | HQYCLCRTVDGLHL | YLCRTVDGL | 2 | 0 | 0 | 1 | 1 | YCLCRTVDGL | vdchain | 0.028 | 36967.64 | 18.00 |
| 11 | HLA-A0205 | QYCLCRTVDGLHLH | YLCRTVDGL | 1 | 0 | 0 | 1 | 1 | YCLCRTVDGL | vdchain | 0.025 | 38200.09 | 20.00 |
| 12 | HLA-A0205 | YCLCRTVDGLHLHR | YLCRTVDGL | 0 | 0 | 0 | 1 | 1 | YCLCRTVDGL | vdchain | 0.025 | 38195.12 | 20.00 |
| 13 | HLA-A0205 | CLCRTVDGLHLHRK | CLVDGLHLH | 0 | 0 | 0 | 2 | 3 | CLCRTVDGLHLH | vdchain | 0.014 | 42792.79 | 33.00 |
| 14 | HLA-A0205 | LCRTVDGLHLHRKP | LCRTVGLHL | 0 | 0 | 0 | 5 | 1 | LCRTVDGLHL | vdchain | 0.006 | 47043.10 | 70.00 |
| 15 | HLA-A0205 | CRTVDGLHLHRKPV | CLHLHRKPV | 0 | 0 | 0 | 1 | 5 | CRTVDGLHLHRKPV | vdchain | 0.035 | 34381.10 | 15.00 |
| 16 | HLA-A0205 | RTVDGLHLHRKPVK | RTVDGLHPV | 0 | 0 | 0 | 7 | 4 | RTVDGLHLHRKPV | vdchain | 0.030 | 36091.83 | 17.00 |
| 17 | HLA-A0205 | TVDGLHLHRKPVKR | VLHLHRKPV | 1 | 0 | 0 | 1 | 2 | VDGLHLHRKPV | vdchain | 0.025 | 38187.29 | 20.00 |
| 18 | HLA-A0205 | VDGLHLHRKPVKRL | VLHLHRKPV | 0 | 0 | 0 | 1 | 2 | VDGLHLHRKPV | vdchain | 0.026 | 37810.68 | 20.00 |
| 19 | HLA-A0205 | DGLHLHRKPVKRLC | GLHLHRKPV | 1 | 0 | 0 | 0 | 0 | GLHLHRKPV | vdchain | 0.022 | 39617.80 | 23.00 |

Protein vdchain. Allele HLA-A0205. Number of high binders 0. Number of weak binders 0. Number of peptides 20

Link to Allele Frequencies in Worldwide Populations HLA-A0205

\# Rank Threshold for Strong binding peptides 0.500
\# Rank Threshold for Weakbindingpeptides 2.000

| pos | HLA | peptide | CoreOffset | I_pos | I_len | D_pos | D_len | iCore | Identity | 1-log50k(aff) | Affinity(nM) | %Rank | BindLevel |
|---|---|---|---|---|---|---|---|---|---|---|---|---|---|
| 0 | HLA-A0206 | HCRCQKPCIRHQYC | CQKPCIQYC | 3 | 0 | 0 | 6 | 2 | CQKPCIRHQYC | vdchain | 0.020 | 40313.12 | 80.00 |
| 1 | HLA-A0206 | CRCQKPCIRHQYCL | CQKPCIYCL | 2 | 0 | 0 | 6 | 3 | CQKPCIRHQYCL | vdchain | 0.063 | 25390.63 | 43.00 |
| 2 | HLA-A0206 | RCQKPCIRHQYCLC | CQKPCIYCL | 1 | 0 | 0 | 6 | 3 | CQKPCIRHQYCL | vdchain | 0.044 | 31062.34 | 55.00 |
| 3 | HLA-A0206 | CQKPCIRHQYCLCR | CQKPCIYCL | 0 | 0 | 0 | 6 | 3 | CQKPCIRHQYCL | vdchain | 0.050 | 29116.79 | 49.00 |
| 4 | HLA-A0206 | QKPCIRHQYCLCRT | CIRHCLCRT | 3 | 0 | 0 | 4 | 2 | CIRHQYCLCRT | vdchain | 0.028 | 37012.06 | 70.00 |
| 5 | HLA-A0206 | KPCIRHQYCLCRTV | KQYCLCRTV | 0 | 0 | 0 | 1 | 5 | KPCIRHQYCLCRTV | vdchain | 0.300 | 1939.86 | 7.50 |
| 6 | HLA-A0206 | PCIRHQYCLCRTVD | IQYCLCRTV | 2 | 0 | 0 | 1 | 2 | IRHQYCLCRTV | vdchain | 0.158 | 9026.86 | 19.00 |
| 7 | HLA-A0206 | CIRHQYCLCRTVDG | CQYCLCRTV | 0 | 0 | 0 | 1 | 3 | CIRHQYCLCRTV | vdchain | 0.164 | 8484.58 | 18.00 |
| 8 | HLA-A0206 | IRHQYCLCRTVDGL | IQYCLCRTV | 0 | 0 | 0 | 1 | 2 | IRHQYCLCRTV | vdchain | 0.183 | 6927.12 | 16.00 |
| 9 | HLA-A0206 | RHQYCLCRTVDGLH | RQYCLCRTV | 0 | 0 | 0 | 1 | 1 | RHQYCLCRTV | vdchain | 0.144 | 10565.56 | 21.00 |
| 10 | HLA-A0206 | HQYCLCRTVDGLHL | HQYCLCLHL | 0 | 0 | 0 | 6 | 5 | HQYCLCRTVDGLHL | vdchain | 0.267 | 2768.71 | 9.00 |
| 11 | HLA-A0206 | QYCLCRTVDGLHLH | YTVDGLHLH | 1 | 0 | 0 | 1 | 4 | YCLCRTVDGLHLH | vdchain | 0.050 | 28954.07 | 49.00 |
| 12 | HLA-A0206 | YCLCRTVDGLHLHR | YVDGLHLHR | 0 | 0 | 0 | 1 | 5 | YCLCRTVDGLHLHR | vdchain | 0.057 | 27078.98 | 46.00 |
| 13 | HLA-A0206 | CLCRTVDGLHLHRK | CLCRTVLHL | 0 | 0 | 0 | 6 | 2 | CLCRTVDGLHL | vdchain | 0.048 | 29623.65 | 50.00 |
| 14 | HLA-A0206 | LCRTVDGLHLHRKP | RTVDGLHLP | 2 | 0 | 0 | 8 | 3 | RTVDGLHLHRKP | vdchain | 0.027 | 37347.17 | 70.00 |
| 15 | HLA-A0206 | CRTVDGLHLHRKPV | RTVDGLHPV | 1 | 0 | 0 | 7 | 4 | RTVDGLHLHRKPV | vdchain | 0.400 | 658.29 | 4.00 |
| 16 | HLA-A0206 | RTVDGLHLHRKPVK | RTVDGLHPV | 0 | 0 | 0 | 7 | 4 | RTVDGLHLHRKPV | vdchain | 0.318 | 1602.37 | 6.50 |
| 17 | HLA-A0206 | TVDGLHLHRKPVKR | TVDGLHLPV | 0 | 0 | 0 | 7 | 3 | TVDGLHLHRKPV | vdchain | 0.139 | 11139.62 | 22.00 |
| 18 | HLA-A0206 | VDGLHLHRKPVKRL | VLHLHRKPV | 0 | 0 | 0 | 1 | 2 | VDGLHLHRKPV | vdchain | 0.045 | 30772.97 | 55.00 |
| 19 | HLA-A0206 | DGLHLHRKPVKRLC | GLHLHRKPV | 1 | 0 | 0 | 0 | 0 | GLHLHRKPV | vdchain | 0.027 | 37336.64 | 70.00 |

Protein vdchain. Allele HLA-A0206. Number of high binders 0. Number of weak binders 0. Number of peptides 20

Link to Allele Frequencies in Worldwide Populations HLA-A0206

\# Rank Threshold for Strongbindingpeptides 0.500

\# Rank Threshold for Weakbindingpeptides 2.000

| pos | HLA | peptide | CoreOffset | I_pos | I_len | D_pos | D_len | iCore | Identity | 1-log50k(aff) | Affinity(nM) | %Rank | BindLevel |
|---|---|---|---|---|---|---|---|---|---|---|---|---|---|
| 0 | HLA-A0207 | HCRCQKPCIRHQYC | HCRCQKPCI | 0 | 0 | 0 | 0 | 0 | HCRCQKPCI | vdchain | 0.014 | 42796.49 | 40.00 |
| 1 | HLA-A0207 | CRCQKPCIRHQYCL | QKPCIQYCL | 3 | 0 | 0 | 5 | 2 | QKPCIRHQYCL | vdchain | 0.015 | 42447.41 | 37.00 |
| 2 | HLA-A0207 | RCQKPCIRHQYCLC | QKPCIYCLC | 2 | 0 | 0 | 5 | 3 | QKPCIRHQYCLC | vdchain | 0.016 | 42029.68 | 34.00 |
| 3 | HLA-A0207 | CQKPCIRHQYCLCR | QKPCIYCLC | 1 | 0 | 0 | 5 | 3 | QKPCIRHQYCLC | vdchain | 0.015 | 42310.29 | 36.00 |
| 4 | HLA-A0207 | QKPCIRHQYCLCRT | QKPCIYCLC | 0 | 0 | 0 | 5 | 3 | QKPCIRHQYCLC | vdchain | 0.015 | 42297.03 | 36.00 |
| 5 | HLA-A0207 | KPCIRHQYCLCRTV | RHQYCLCTV | 4 | 0 | 0 | 7 | 1 | RHQYCLCRTV | vdchain | 0.025 | 37952.50 | 18.00 |
| 6 | HLA-A0207 | PCIRHQYCLCRTVD | RHQYCLCTV | 3 | 0 | 0 | 7 | 1 | RHQYCLCRTV | vdchain | 0.025 | 38278.69 | 19.00 |
| 7 | HLA-A0207 | CIRHQYCLCRTVDG | RHQYCLCTV | 2 | 0 | 0 | 7 | 1 | RHQYCLCRTV | vdchain | 0.024 | 38407.30 | 19.00 |
| 8 | HLA-A0207 | IRHQYCLCRTVDGL | RHQYCLCTV | 1 | 0 | 0 | 7 | 1 | RHQYCLCRTV | vdchain | 0.021 | 39826.25 | 24.00 |
| 9 | HLA-A0207 | RHQYCLCRTVDGLH | RHQYCLCTV | 0 | 0 | 0 | 7 | 1 | RHQYCLCRTV | vdchain | 0.021 | 39867.65 | 24.00 |
| 10 | HLA-A0207 | HQYCLCRTVDGLHL | YLCRTVDGL | 2 | 0 | 0 | 1 | 1 | YCLCRTVDGL | vdchain | 0.017 | 41411.29 | 31.00 |
| 11 | HLA-A0207 | QYCLCRTVDGLHLH | YLCRTVDGL | 1 | 0 | 0 | 1 | 1 | YCLCRTVDGL | vdchain | 0.016 | 42020.14 | 34.00 |
| 12 | HLA-A0207 | YCLCRTVDGLHLHR | YVDGLHLHR | 0 | 0 | 0 | 1 | 5 | YCLCRTVDGLHLHR | vdchain | 0.018 | 41013.52 | 29.00 |
| 13 | HLA-A0207 | CLCRTVDGLHLHRK | CLDGLHLHR | 0 | 0 | 0 | 2 | 4 | CLCRTVDGLHLHR | vdchain | 0.015 | 42331.36 | 36.00 |
| 14 | HLA-A0207 | LCRTVDGLHLHRKP | RVDGLHLHR | 2 | 0 | 0 | 1 | 1 | RTVDGLHLHR | vdchain | 0.011 | 44551.29 | 60.00 |
| 15 | HLA-A0207 | CRTVDGLHLHRKPV | TVDGLHLPV | 2 | 0 | 0 | 7 | 3 | TVDGLHLHRKPV | vdchain | 0.082 | 20630.50 | 2.00 | <= WB |
| 16 | HLA-A0207 | RTVDGLHLHRKPVK | RLHLHRKPV | 0 | 0 | 0 | 1 | 4 | RTVDGLHLHRKPV | vdchain | 0.082 | 20557.64 | 1.90 | <= WB |
| 17 | HLA-A0207 | TVDGLHLHRKPVKR | GLHLHRKPV | 3 | 0 | 0 | 0 | 0 | GLHLHRKPV | vdchain | 0.077 | 21756.83 | 2.50 |
| 18 | HLA-A0207 | VDGLHLHRKPVKRL | GLHLHRKPV | 2 | 0 | 0 | 0 | 0 | GLHLHRKPV | vdchain | 0.057 | 27101.85 | 4.50 |
| 19 | HLA-A0207 | DGLHLHRKPVKRLC | GLHLHRKPV | 1 | 0 | 0 | 0 | 0 | GLHLHRKPV | vdchain | 0.057 | 26987.73 | 4.50 |

Protein vdchain. Allele HLA-A0207. Number of high binders 0. Number of weak binders 2. Number of peptides 20

Link to Allele Frequencies in Worldwide PopulationsHLA-A0207

\# Rank Threshold for Strong binding peptides 0.500
\# Rank Threshold for Weakbindingpeptides 2.000

| pos | HLA | peptide | CoreOffset | I_pos | I_len | D_pos | D_len | iCore | Identity | 1-log50k(aff) | Affinity(nM) | %Rank | BindLevel |
|---|---|---|---|---|---|---|---|---|---|---|---|---|---|
| 0 | HLA-A0211 | HCRCQKPCIRHQYC | CQCIRHQYC | 3 | 0 | 0 | 2 | 2 | CQKPCIRHQYC | vdchain | 0.011 | 44154.41 | 60.00 |
| 1 | HLA-A0211 | CRCQKPCIRHQYCL | CQKPCIYCL | 2 | 0 | 0 | 6 | 3 | CQKPCIRHQYCL | vdchain | 0.019 | 40896.54 | 43.00 |
| 2 | HLA-A0211 | RCQKPCIRHQYCLC | CQKPCIYCL | 1 | 0 | 0 | 6 | 3 | CQKPCIRHQYCL | vdchain | 0.017 | 41407.25 | 45.00 |
| 3 | HLA-A0211 | CQKPCIRHQYCLCR | CQKPCIYCL | 0 | 0 | 0 | 6 | 3 | CQKPCIRHQYCL | vdchain | 0.017 | 41520.31 | 45.00 |
| 4 | HLA-A0211 | QKPCIRHQYCLCRT | CIQYCLCRT | 3 | 0 | 0 | 2 | 2 | CIRHQYCLCRT | vdchain | 0.012 | 43809.42 | 60.00 |
| 5 | HLA-A0211 | KPCIRHQYCLCRTV | CIYCLCRTV | 2 | 0 | 0 | 2 | 3 | CIRHQYCLCRTV | vdchain | 0.056 | 27232.36 | 19.00 |
| 6 | HLA-A0211 | PCIRHQYCLCRTVD | CIYCLCRTV | 1 | 0 | 0 | 2 | 3 | CIRHQYCLCRTV | vdchain | 0.046 | 30246.79 | 22.00 |
| 7 | HLA-A0211 | CIRHQYCLCRTVDG | CIYCLCRTV | 0 | 0 | 0 | 2 | 3 | CIRHQYCLCRTV | vdchain | 0.050 | 29222.21 | 21.00 |
| 8 | HLA-A0211 | IRHQYCLCRTVDGL | YLCRTVDGL | 4 | 0 | 0 | 1 | 1 | YCLCRTVDGL | vdchain | 0.082 | 20588.13 | 15.00 |
| 9 | HLA-A0211 | RHQYCLCRTVDGLH | YLCRTVDGL | 3 | 0 | 0 | 1 | 1 | YCLCRTVDGL | vdchain | 0.066 | 24456.35 | 17.00 |
| 10 | HLA-A0211 | HQYCLCRTVDGLHL | YLCRTVDGL | 2 | 0 | 0 | 1 | 1 | YCLCRTVDGL | vdchain | 0.082 | 20654.62 | 15.00 |
| 11 | HLA-A0211 | QYCLCRTVDGLHLH | YLCRTVDGL | 1 | 0 | 0 | 1 | 1 | YCLCRTVDGL | vdchain | 0.068 | 23830.49 | 17.00 |
| 12 | HLA-A0211 | YCLCRTVDGLHLHR | YLCRTVDGL | 0 | 0 | 0 | 1 | 1 | YCLCRTVDGL | vdchain | 0.074 | 22533.56 | 16.00 |
| 13 | HLA-A0211 | CLCRTVDGLHLHRK | CLCRTVDGL | 0 | 0 | 0 | 0 | 0 | CLCRTVDGL | vdchain | 0.071 | 23264.40 | 16.00 |
| 14 | HLA-A0211 | LCRTVDGLHLHRKP | RTVDGLHLP | 2 | 0 | 0 | 8 | 3 | RTVDGLHLHRKP | vdchain | 0.008 | 45623.49 | 70.00 |
| 15 | HLA-A0211 | CRTVDGLHLHRKPV | RTVDGLHPV | 1 | 0 | 0 | 7 | 4 | RTVDGLHLHRKPV | vdchain | 0.061 | 25902.02 | 18.00 |

| | | | | | | | | | | | | | |
|---|---|---|---|---|---|---|---|---|---|---|---|---|---|
| 16 | HLA-A0211 | RTVDGLHLHRKPVK | RTVDGLHPV | 0 | 0 | 0 | 7 | 4 | RTVDGLHLHRKPV | vdchain | 0.059 | 26431.44 | 19.00 |
| 17 | HLA-A0211 | TVDGLHLHRKPVKR | TVDGLHLHV | 0 | 0 | 0 | 8 | 3 | TVDGLHLHRKPV | vdchain | 0.028 | 37129.18 | 33.00 |
| 18 | HLA-A0211 | VDGLHLHRKPVKRL | VLHLHRKPV | 0 | 0 | 0 | 1 | 2 | VDGLHLHRKPV | vdchain | 0.021 | 39933.26 | 40.00 |
| 19 | HLA-A0211 | DGLHLHRKPVKRLC | GLHLHRKPV | 1 | 0 | 0 | 0 | 0 | GLHLHRKPV | vdchain | 0.017 | 41378.13 | 45.00 |

Protein vdchain. Allele HLA-A0211. Number of high binders 0. Number of weak binders 0. Number of peptides 20

Link to Allele Frequencies in Worldwide PopulationsHLA-A0211

### Rank Threshold for Strong binding peptides 0.500
### Rank Threshold for Weakbindingpeptides 2.000

| pos | HLA | peptide | CoreOffset | I_pos | I_len | D_pos | D_len | iCore | Identity | 1-log50k(aff) | Affinity(nM) | %Rank | BindLevel |
|---|---|---|---|---|---|---|---|---|---|---|---|---|---|
| 0 | HLA-A0212 | HCRCQKPCIRHQYC | CQKPCIQYC | 3 | 0 | 0 | 6 | 2 | CQKPCIRHQYC | vdchain | 0.014 | 42987.24 | 60.00 |
| 1 | HLA-A0212 | CRCQKPCIRHQYCL | CQIRHQYCL | 2 | 0 | 0 | 2 | 3 | CQKPCIRHQYCL | vdchain | 0.046 | 30311.99 | 20.00 |
| 2 | HLA-A0212 | RCQKPCIRHQYCLC | CQIRHQYCL | 1 | 0 | 0 | 2 | 3 | CQKPCIRHQYCL | vdchain | 0.038 | 33068.75 | 24.00 |
| 3 | HLA-A0212 | CQKPCIRHQYCLCR | CQIRHQYCL | 0 | 0 | 0 | 2 | 3 | CQKPCIRHQYCL | vdchain | 0.039 | 32895.67 | 23.00 |
| 4 | HLA-A0212 | QKPCIRHQYCLCRT | KCIRHQYCL | 1 | 0 | 0 | 1 | 1 | KPCIRHQYCL | vdchain | 0.014 | 42957.48 | 60.00 |
| 5 | HLA-A0212 | KPCIRHQYCLCRTV | HQYCLCRTV | 5 | 0 | 0 | 0 | 0 | HQYCLCRTV | vdchain | 0.032 | 35403.76 | 28.00 |
| 6 | HLA-A0212 | PCIRHQYCLCRTVD | HQYCLCRTV | 4 | 0 | 0 | 0 | 0 | HQYCLCRTV | vdchain | 0.027 | 37371.00 | 33.00 |
| 7 | HLA-A0212 | CIRHQYCLCRTVDG | CIYCLCRTV | 0 | 0 | 0 | 2 | 3 | CIRHQYCLCRTV | vdchain | 0.027 | 37287.80 | 32.00 |
| 8 | HLA-A0212 | IRHQYCLCRTVDGL | CLCRTVDGL | 5 | 0 | 0 | 0 | 0 | CLCRTVDGL | vdchain | 0.068 | 24076.43 | 14.00 |
| 9 | HLA-A0212 | RHQYCLCRTVDGLH | CLCRTVDGL | 4 | 0 | 0 | 0 | 0 | CLCRTVDGL | vdchain | 0.055 | 27451.57 | 17.00 |
| 10 | HLA-A0212 | HQYCLCRTVDGLHL | YLCRTVDGL | 2 | 0 | 0 | 1 | 1 | YCLCRTVDGL | vdchain | 0.054 | 27739.39 | 17.00 |
| 11 | HLA-A0212 | QYCLCRTVDGLHLH | YLCRTVDGL | 1 | 0 | 0 | 1 | 1 | YCLCRTVDGL | vdchain | 0.053 | 28301.77 | 18.00 |
| 12 | HLA-A0212 | YCLCRTVDGLHLHR | YLCRTVDGL | 0 | 0 | 0 | 1 | 1 | YCLCRTVDGL | vdchain | 0.056 | 27379.79 | 17.00 |
| 13 | HLA-A0212 | CLCRTVDGLHLHRK | CLCRTVDGL | 0 | 0 | 0 | 0 | 0 | CLCRTVDGL | vdchain | 0.046 | 30311.99 | 20.00 |
| 14 | HLA-A0212 | LCRTVDGLHLHRKP | RTVDGLHLP | 2 | 0 | 0 | 8 | 3 | RTVDGLHLHRKP | vdchain | 0.009 | 45472.18 | 80.00 |
| 15 | HLA-A0212 | CRTVDGLHLHRKPV | RTVDGLHPV | 1 | 0 | 0 | 7 | 4 | RTVDGLHLHRKPV | vdchain | 0.053 | 28151.21 | 18.00 |
| 16 | HLA-A0212 | RTVDGLHLHRKPVK | RTVDGLHPV | 0 | 0 | 0 | 7 | 4 | RTVDGLHLHRKPV | vdchain | 0.046 | 30362.54 | 20.00 |
| 17 | HLA-A0212 | TVDGLHLHRKPVKR | GLHLHRKPV | 3 | 0 | 0 | 0 | 0 | GLHLHRKPV | vdchain | 0.023 | 38871.78 | 37.00 |
| 18 | HLA-A0212 | VDGLHLHRKPVKRL | GLHLHRKPV | 2 | 0 | 0 | 0 | 0 | GLHLHRKPV | vdchain | 0.023 | 38940.38 | 37.00 |
| 19 | HLA-A0212 | DGLHLHRKPVKRLC | GLHLHRKPV | 1 | 0 | 0 | 0 | 0 | GLHLHRKPV | vdchain | 0.023 | 38927.32 | 37.00 |

Protein vdchain. Allele HLA-A0212. Number of high binders 0. Number of weak binders 0. Number of peptides 20

Link to Allele Frequencies in Worldwide PopulationsHLA-A0212

### Rank Threshold for Strong binding peptides 0.500
### Rank Threshold for Weakbindingpeptides 2.000

| pos | HLA | peptide | CoreOffset | I_pos | I_len | D_pos | D_len | iCore | Identity | 1-log50k(aff) | Affinity(nM) | %Rank | BindLevel |
|---|---|---|---|---|---|---|---|---|---|---|---|---|---|
| 0 | HLA-A0216 | HCRCQKPCIRHQYC | HCRCQKPCI | 0 | 0 | 0 | 0 | 0 | HCRCQKPCI | vdchain | 0.012 | 43974.68 | 55.00 |
| 1 | HLA-A0216 | CRCQKPCIRHQYCL | CQKPCIYCL | 2 | 0 | 0 | 6 | 3 | CQKPCIRHQYCL | vdchain | 0.033 | 35101.66 | 25.00 |
| 2 | HLA-A0216 | RCQKPCIRHQYCLC | CQKPCIYCL | 1 | 0 | 0 | 6 | 3 | CQKPCIRHQYCL | vdchain | 0.029 | 36363.06 | 27.00 |
| 3 | HLA-A0216 | CQKPCIRHQYCLCR | CQKPCIYCL | 0 | 0 | 0 | 6 | 3 | CQKPCIRHQYCL | vdchain | 0.033 | 35121.43 | 25.00 |
| 4 | HLA-A0216 | QKPCIRHQYCLCRT | KCIRHQYCL | 1 | 0 | 0 | 1 | 1 | KPCIRHQYCL | vdchain | 0.014 | 43138.18 | 49.00 |

| | | | | | | | | | | | | | |
|---|---|---|---|---|---|---|---|---|---|---|---|---|---|
| 5 | HLA-A0216 | KPCIRHQYCLCRTV | KQYCLCRTV | 0 | 0 | 0 | 1 | 5 | KPCIRHQYCLCRTV | vdchain | 0.025 | 38000.16 | 31.00 |
| 6 | HLA-A0216 | PCIRHQYCLCRTVD | HQYCLCRTV | 4 | 0 | 0 | 0 | 0 | HQYCLCRTV | vdchain | 0.021 | 39768.55 | 35.00 |
| 7 | HLA-A0216 | CIRHQYCLCRTVDG | CQYCLCRTV | 0 | 0 | 0 | 1 | 3 | CIRHQYCLCRTV | vdchain | 0.022 | 39381.05 | 34.00 |
| 8 | HLA-A0216 | IRHQYCLCRTVDGL | YLCRTVDGL | 4 | 0 | 0 | 1 | 1 | YCLCRTVDGL | vdchain | 0.035 | 34387.44 | 24.00 |
| 9 | HLA-A0216 | RHQYCLCRTVDGLH | YLCRTVDGL | 3 | 0 | 0 | 1 | 1 | YCLCRTVDGL | vdchain | 0.031 | 35861.00 | 26.00 |
| 10 | HLA-A0216 | HQYCLCRTVDGLHL | YLCRTVDGL | 2 | 0 | 0 | 1 | 1 | YCLCRTVDGL | vdchain | 0.031 | 35942.57 | 26.00 |
| 11 | HLA-A0216 | QYCLCRTVDGLHLH | YLCRTVDGL | 1 | 0 | 0 | 1 | 1 | YCLCRTVDGL | vdchain | 0.030 | 36329.63 | 27.00 |
| 12 | HLA-A0216 | YCLCRTVDGLHLHR | YLCRTVDGL | 0 | 0 | 0 | 1 | 1 | YCLCRTVDGL | vdchain | 0.034 | 34533.58 | 24.00 |
| 13 | HLA-A0216 | CLCRTVDGLHLHRK | CLCRTVDGL | 0 | 0 | 0 | 0 | 0 | CLCRTVDGL | vdchain | 0.027 | 37146.07 | 29.00 |
| 14 | HLA-A0216 | LCRTVDGLHLHRKP | LVDGLHLHR | 0 | 0 | 0 | 1 | 3 | LCRTVDGLHLHR | vdchain | 0.006 | 46852.12 | 85.00 |
| 15 | HLA-A0216 | CRTVDGLHLHRKPV | RTVDGLHPV | 1 | 0 | 0 | 7 | 4 | RTVDGLHLHRKPV | vdchain | 0.023 | 38917.23 | 33.00 |
| 16 | HLA-A0216 | RTVDGLHLHRKPVK | RTVDGLHPV | 0 | 0 | 0 | 7 | 4 | RTVDGLHLHRKPV | vdchain | 0.023 | 38913.84 | 33.00 |
| 17 | HLA-A0216 | TVDGLHLHRKPVKR | TVDGLHLPV | 0 | 0 | 0 | 7 | 3 | TVDGLHLHRKPV | vdchain | 0.019 | 40782.11 | 39.00 |
| 18 | HLA-A0216 | VDGLHLHRKPVKRL | GLHLHRKPV | 2 | 0 | 0 | 0 | 0 | GLHLHRKPV | vdchain | 0.012 | 43820.78 | 55.00 |
| 19 | HLA-A0216 | DGLHLHRKPVKRLC | GLHLHRKPV | 1 | 0 | 0 | 0 | 0 | GLHLHRKPV | vdchain | 0.011 | 44306.11 | 60.00 |

---

Protein vdchain. Allele HLA-A0216. Number of high binders 0. Number of weak binders 0. Number of peptides 20

Link to Allele Frequencies in Worldwide PopulationsHLA-A0216

---

### Rank Threshold for Strong binding peptides 0.500
### Rank Threshold for Weakbindingpeptides 2.000

---

| pos | HLA | peptide | CoreOffset | I_pos | I_len | D_pos | D_len | iCore | Identity | 1-log50k(aff) | Affinity(nM) | %Rank | BindLevel |
|---|---|---|---|---|---|---|---|---|---|---|---|---|---|
| 0 | HLA-A0217 | HCRCQKPCIRHQYC | RCQKPCIYC | 2 | 0 | 0 | 7 | 3 | RCQKPCIRHQYC | vdchain | 0.006 | 46770.57 | 80.00 |
| 1 | HLA-A0217 | CRCQKPCIRHQYCL | RCQKPCIRL | 1 | 0 | 0 | 8 | 4 | RCQKPCIRHQYCL | vdchain | 0.017 | 41414.87 | 43.00 |
| 2 | HLA-A0217 | RCQKPCIRHQYCLC | RCQKPCIRL | 0 | 0 | 0 | 8 | 4 | RCQKPCIRHQYCL | vdchain | 0.014 | 42747.91 | 48.00 |
| 3 | HLA-A0217 | CQKPCIRHQYCLCR | CIHQYCLCR | 4 | 0 | 0 | 2 | 1 | CIRHQYCLCR | vdchain | 0.013 | 43347.34 | 55.00 |
| 4 | HLA-A0217 | QKPCIRHQYCLCRT | CIHQYCLCR | 3 | 0 | 0 | 2 | 1 | CIRHQYCLCR | vdchain | 0.011 | 44362.23 | 60.00 |
| 5 | HLA-A0217 | KPCIRHQYCLCRTV | CIYCLCRTV | 2 | 0 | 0 | 2 | 3 | CIRHQYCLCRTV | vdchain | 0.032 | 35387.67 | 28.00 |
| 6 | HLA-A0217 | PCIRHQYCLCRTVD | CIYCLCRTV | 1 | 0 | 0 | 2 | 3 | CIRHQYCLCRTV | vdchain | 0.027 | 37416.31 | 32.00 |
| 7 | HLA-A0217 | CIRHQYCLCRTVDG | CIYCLCRTV | 0 | 0 | 0 | 2 | 3 | CIRHQYCLCRTV | vdchain | 0.025 | 38231.94 | 34.00 |
| 8 | HLA-A0217 | IRHQYCLCRTVDGL | YLCRTVDGL | 4 | 0 | 0 | 1 | 1 | YCLCRTVDGL | vdchain | 0.091 | 18765.53 | 11.00 |
| 9 | HLA-A0217 | RHQYCLCRTVDGLH | YLCRTVDGL | 3 | 0 | 0 | 1 | 1 | YCLCRTVDGL | vdchain | 0.075 | 22271.77 | 13.00 |
| 10 | HLA-A0217 | HQYCLCRTVDGLHL | CLCRTVLHL | 3 | 0 | 0 | 6 | 2 | CLCRTVDGLHL | vdchain | 0.085 | 20024.60 | 12.00 |
| 11 | HLA-A0217 | QYCLCRTVDGLHLH | CLCRTVLHL | 2 | 0 | 0 | 6 | 2 | CLCRTVDGLHL | vdchain | 0.075 | 22268.88 | 13.00 |
| 12 | HLA-A0217 | YCLCRTVDGLHLHR | YLCRTVDGL | 0 | 0 | 0 | 1 | 1 | YCLCRTVDGL | vdchain | 0.071 | 23140.63 | 14.00 |
| 13 | HLA-A0217 | CLCRTVDGLHLHRK | CLCRTVLHL | 0 | 0 | 0 | 6 | 2 | CLCRTVDGLHL | vdchain | 0.060 | 26012.40 | 16.00 |
| 14 | HLA-A0217 | LCRTVDGLHLHRKP | LCRTVDLHL | 0 | 0 | 0 | 6 | 1 | LCRTVDGLHL | vdchain | 0.009 | 45233.72 | 65.00 |
| 15 | HLA-A0217 | CRTVDGLHLHRKPV | TVDGLHLPV | 2 | 0 | 0 | 7 | 3 | TVDGLHLHRKPV | vdchain | 0.028 | 36935.26 | 31.00 |
| 16 | HLA-A0217 | RTVDGLHLHRKPVK | TVDGLHLPV | 1 | 0 | 0 | 7 | 3 | TVDGLHLHRKPV | vdchain | 0.023 | 38791.09 | 35.00 |
| 17 | HLA-A0217 | TVDGLHLHRKPVKR | TVDGLHLPV | 0 | 0 | 0 | 7 | 3 | TVDGLHLHRKPV | vdchain | 0.022 | 39447.57 | 37.00 |
| 18 | HLA-A0217 | VDGLHLHRKPVKRL | HLHKPVKRL | 4 | 0 | 0 | 3 | 1 | HLHRKPVKRL | vdchain | 0.065 | 24874.53 | 15.00 |
| 19 | HLA-A0217 | DGLHLHRKPVKRLC | HLHKPVKRL | 3 | 0 | 0 | 3 | 1 | HLHRKPVKRL | vdchain | 0.053 | 28086.71 | 18.00 |

---

Protein vdchain. Allele HLA-A0217. Number of high binders 0. Number of weak binders 0. Number of peptides 20

Link to Allele Frequencies in Worldwide PopulationsHLA-A0217

---------------------------------------------------------------------------------------------------
### Rank Threshold for Strong binding peptides 0.500
### Rank Threshold for Weakbindingpeptides 2.000
---------------------------------------------------------------------------------------------------

| pos | HLA | peptide | CoreOffset | I_pos | I_len | D_pos | D_len | | iCore | Identity | 1-log50k(aff) | Affinity(nM) | %Rank | BindLevel |
|---|---|---|---|---|---|---|---|---|---|---|---|---|---|---|
| 0 | HLA-A0219 | HCRCQKPCIRHQYC | CQKPCIQYC | 3 | 0 | 0 | 6 | 2 | CQKPCIRHQYC | vdchain | 0.014 | 43057.05 | 65.00 | |
| 1 | HLA-A0219 | CRCQKPCIRHQYCL | CQIRHQYCL | 2 | 0 | 0 | 2 | 3 | CQKPCIRHQYCL | vdchain | 0.025 | 38003.44 | 33.00 | |
| 2 | HLA-A0219 | RCQKPCIRHQYCLC | CQIRHQYCL | 1 | 0 | 0 | 2 | 3 | CQKPCIRHQYCL | vdchain | 0.025 | 38122.05 | 33.00 | |
| 3 | HLA-A0219 | CQKPCIRHQYCLCR | CQIRHQYCL | 0 | 0 | 0 | 2 | 3 | CQKPCIRHQYCL | vdchain | 0.028 | 37124.37 | 30.00 | |
| 4 | HLA-A0219 | QKPCIRHQYCLCRT | PCIRHQYCL | 2 | 0 | 0 | 0 | 0 | PCIRHQYCL | vdchain | 0.013 | 43289.68 | 65.00 | |
| 5 | HLA-A0219 | KPCIRHQYCLCRTV | CIYCLCRTV | 2 | 0 | 0 | 2 | 3 | CIRHQYCLCRTV | vdchain | 0.017 | 41684.14 | 50.00 | |
| 6 | HLA-A0219 | PCIRHQYCLCRTVD | CIYCLCRTV | 1 | 0 | 0 | 2 | 3 | CIRHQYCLCRTV | vdchain | 0.016 | 42094.32 | 55.00 | |
| 7 | HLA-A0219 | CIRHQYCLCRTVDG | CIYCLCRTV | 0 | 0 | 0 | 2 | 3 | CIRHQYCLCRTV | vdchain | 0.018 | 41361.14 | 48.00 | |
| 8 | HLA-A0219 | IRHQYCLCRTVDGL | YLCRTVDGL | 4 | 0 | 0 | 1 | 1 | YCLCRTVDGL | vdchain | 0.048 | 29747.63 | 17.00 | |
| 9 | HLA-A0219 | RHQYCLCRTVDGLH | YLCRTVDGL | 3 | 0 | 0 | 1 | 1 | YCLCRTVDGL | vdchain | 0.047 | 29924.21 | 17.00 | |
| 10 | HLA-A0219 | HQYCLCRTVDGLHL | YLCRTVDGL | 2 | 0 | 0 | 1 | 1 | YCLCRTVDGL | vdchain | 0.047 | 29990.34 | 17.00 | |
| 11 | HLA-A0219 | QYCLCRTVDGLHLH | YLCRTVDGL | 1 | 0 | 0 | 1 | 1 | YCLCRTVDGL | vdchain | 0.049 | 29302.32 | 16.00 | |
| 12 | HLA-A0219 | YCLCRTVDGLHLHR | YLCRTVDGL | 0 | 0 | 0 | 1 | 1 | YCLCRTVDGL | vdchain | 0.056 | 27407.95 | 14.00 | |
| 13 | HLA-A0219 | CLCRTVDGLHLHRK | CLCRTVDGL | 0 | 0 | 0 | 0 | 0 | CLCRTVDGL | vdchain | 0.052 | 28503.37 | 15.00 | |
| 14 | HLA-A0219 | LCRTVDGLHLHRKP | CTVDGLHLH | 1 | 0 | 0 | 1 | 1 | CRTVDGLHLH | vdchain | 0.009 | 45164.75 | 85.00 | |
| 15 | HLA-A0219 | CRTVDGLHLHRKPV | RTVDGLHPV | 1 | 0 | 0 | 7 | 4 | RTVDGLHLHRKPV | vdchain | 0.028 | 37002.47 | 29.00 | |
| 16 | HLA-A0219 | RTVDGLHLHRKPVK | RTVDGLHPV | 0 | 0 | 0 | 7 | 4 | RTVDGLHLHRKPV | vdchain | 0.030 | 36175.49 | 27.00 | |
| 17 | HLA-A0219 | TVDGLHLHRKPVKR | TVDGLHLPV | 0 | 0 | 0 | 7 | 3 | TVDGLHLHRKPV | vdchain | 0.021 | 40046.64 | 41.00 | |
| 18 | HLA-A0219 | VDGLHLHRKPVKRL | VDGLHLHPV | 0 | 0 | 0 | 7 | 2 | VDGLHLHRKPV | vdchain | 0.021 | 39984.29 | 40.00 | |
| 19 | HLA-A0219 | DGLHLHRKPVKRLC | GLHLHRKPV | 1 | 0 | 0 | 0 | 0 | GLHLHRKPV | vdchain | 0.017 | 41422.48 | 49.00 | |

Protein vdchain. Allele HLA-A0219. Number of high binders 0. Number of weak binders 0. Number of peptides 20

Link to Allele Frequencies in Worldwide PopulationsHLA-A0219

---------------------------------------------------------------------------------------------------
### Rank Threshold for Strong binding peptides 0.500
### Rank Threshold for Weakbindingpeptides 2.000
---------------------------------------------------------------------------------------------------

| pos | HLA | peptide | CoreOffset | I_pos | I_len | D_pos | D_len | | iCore | Identity | 1-log50k(aff) | Affinity(nM) | %Rank | BindLevel |
|---|---|---|---|---|---|---|---|---|---|---|---|---|---|---|
| 0 | HLA-A0250 | HCRCQKPCIRHQYC | CQKPCIQYC | 3 | 0 | 0 | 6 | 2 | CQKPCIRHQYC | vdchain | 0.008 | 45773.79 | 55.00 | |
| 1 | HLA-A0250 | CRCQKPCIRHQYCL | QKPCIRHQL | 3 | 0 | 0 | 8 | 2 | QKPCIRHQYCL | vdchain | 0.008 | 45948.47 | 60.00 | |
| 2 | HLA-A0250 | RCQKPCIRHQYCLC | CIRHQYCLC | 5 | 0 | 0 | 0 | 0 | CIRHQYCLC | vdchain | 0.037 | 33576.37 | 24.00 | |
| 3 | HLA-A0250 | CQKPCIRHQYCLCR | CIRHQYCLC | 4 | 0 | 0 | 0 | 0 | CIRHQYCLC | vdchain | 0.026 | 37535.95 | 29.00 | |
| 4 | HLA-A0250 | QKPCIRHQYCLCRT | CIRHQYCLC | 3 | 0 | 0 | 0 | 0 | CIRHQYCLC | vdchain | 0.029 | 36544.50 | 28.00 | |
| 5 | HLA-A0250 | KPCIRHQYCLCRTV | CIRHQYCTV | 2 | 0 | 0 | 7 | 3 | CIRHQYCLCRTV | vdchain | 0.039 | 32717.48 | 23.00 | |
| 6 | HLA-A0250 | PCIRHQYCLCRTVD | CIRHQYCTV | 1 | 0 | 0 | 7 | 3 | CIRHQYCLCRTV | vdchain | 0.031 | 35601.58 | 26.00 | |
| 7 | HLA-A0250 | CIRHQYCLCRTVDG | CIRHQYCTV | 0 | 0 | 0 | 7 | 3 | CIRHQYCLCRTV | vdchain | 0.033 | 34952.73 | 26.00 | |
| 8 | HLA-A0250 | IRHQYCLCRTVDGL | YLCRTVDGL | 4 | 0 | 0 | 1 | 1 | YCLCRTVDGL | vdchain | 0.037 | 33374.26 | 24.00 | |
| 9 | HLA-A0250 | RHQYCLCRTVDGLH | YLCRTVDGL | 3 | 0 | 0 | 1 | 1 | YCLCRTVDGL | vdchain | 0.029 | 36471.82 | 28.00 | |
| 10 | HLA-A0250 | HQYCLCRTVDGLHL | YLCRTVDGL | 2 | 0 | 0 | 1 | 1 | YCLCRTVDGL | vdchain | 0.028 | 37112.30 | 29.00 | |
| 11 | HLA-A0250 | QYCLCRTVDGLHLH | YLCRTVDGL | 1 | 0 | 0 | 1 | 1 | YCLCRTVDGL | vdchain | 0.037 | 33585.82 | 24.00 | |
| 12 | HLA-A0250 | YCLCRTVDGLHLHR | YLCRTVDGL | 0 | 0 | 0 | 1 | 1 | YCLCRTVDGL | vdchain | 0.034 | 34676.61 | 25.00 | |
| 13 | HLA-A0250 | CLCRTVDGLHLHRK | CLCDGLHLH | 0 | 0 | 0 | 3 | 3 | CLCRTVDGLHLH | vdchain | 0.026 | 37721.18 | 30.00 | |

| | | | | | | | | | | | | | |
|---|---|---|---|---|---|---|---|---|---|---|---|---|---|
| 14 | HLA-A0250 | LCRTVDGLHLHRKP | RTVDGLHLP | 2 | 0 | 0 | 8 | 3 | RTVDGLHLHRKP | vdchain | 0.006 | 47060.91 | 65.00 |
| 15 | HLA-A0250 | CRTVDGLHLHRKPV | GLHLHRKPV | 5 | 0 | 0 | 0 | 0 | GLHLHRKPV | vdchain | 0.035 | 34349.49 | 25.00 |
| 16 | HLA-A0250 | RTVDGLHLHRKPVK | RTVDGLHLV | 0 | 0 | 0 | 8 | 4 | RTVDGLHLHRKPV | vdchain | 0.028 | 37094.65 | 29.00 |
| 17 | HLA-A0250 | TVDGLHLHRKPVKR | GLHLHRKPV | 3 | 0 | 0 | 0 | 0 | GLHLHRKPV | vdchain | 0.013 | 43660.81 | 44.00 |
| 18 | HLA-A0250 | VDGLHLHRKPVKRL | HLHKPVKRL | 4 | 0 | 0 | 3 | 1 | HLHRKPVKRL | vdchain | 0.015 | 42409.75 | 40.00 |
| 19 | HLA-A0250 | DGLHLHRKPVKRLC | GLHLHRKPV | 1 | 0 | 0 | 0 | 0 | GLHLHRKPV | vdchain | 0.017 | 41493.36 | 38.00 |

Protein vdchain. Allele HLA-A0250. Number of high binders 0. Number of weak binders 0. Number of peptides 20

Link to Allele Frequencies in Worldwide Populations HLA-A0250

\# Rank Threshold for Strong binding peptides 0.500
\# Rank Threshold for Weakbindingpeptides 2.000

| pos | HLA | peptide | Core | Offset | I_pos | I_len | D_pos | D_len | iCore | Identity | 1-log50k(aff) | Affinity(nM) | %Rank | BindLevel |
|---|---|---|---|---|---|---|---|---|---|---|---|---|---|---|
| 0 | HLA-A0301 | HCRCQKPCIRHQYC | RCQKPCIRY | 2 | 0 | 0 | 8 | 2 | RCQKPCIRHQY | vdchain | 0.039 | 32848.37 | 47.00 | |
| 1 | HLA-A0301 | CRCQKPCIRHQYCL | RCQKPCIRY | 1 | 0 | 0 | 8 | 2 | RCQKPCIRHQY | vdchain | 0.038 | 32995.46 | 48.00 | |
| 2 | HLA-A0301 | RCQKPCIRHQYCLC | RIRHQYCLC | 0 | 0 | 0 | 1 | 5 | RCQKPCIRHQYCLC | vdchain | 0.071 | 23153.65 | 23.00 | |
| 3 | HLA-A0301 | CQKPCIRHQYCLCR | CIRHQYCLR | 4 | 0 | 0 | 8 | 1 | CIRHQYCLCR | vdchain | 0.112 | 14928.14 | 13.00 | |
| 4 | HLA-A0301 | QKPCIRHQYCLCRT | CIRHQYCLR | 3 | 0 | 0 | 8 | 1 | CIRHQYCLCR | vdchain | 0.081 | 20793.86 | 20.00 | |
| 5 | HLA-A0301 | KPCIRHQYCLCRTV | CIRHQYCLR | 2 | 0 | 0 | 8 | 1 | CIRHQYCLCR | vdchain | 0.076 | 22018.28 | 21.00 | |
| 6 | HLA-A0301 | PCIRHQYCLCRTVD | CIRHQYCLR | 1 | 0 | 0 | 8 | 1 | CIRHQYCLCR | vdchain | 0.074 | 22559.91 | 22.00 | |
| 7 | HLA-A0301 | CIRHQYCLCRTVDG | CIRHQYCLR | 0 | 0 | 0 | 8 | 1 | CIRHQYCLCR | vdchain | 0.081 | 20900.33 | 20.00 | |
| 8 | HLA-A0301 | IRHQYCLCRTVDGL | RLCRTVDGL | 1 | 0 | 0 | 1 | 4 | RHQYCLCRTVDGL | vdchain | 0.046 | 30399.69 | 39.00 | |
| 9 | HLA-A0301 | RHQYCLCRTVDGLH | HQYCLCGLH | 1 | 0 | 0 | 6 | 4 | HQYCLCRTVDGLH | vdchain | 0.076 | 22020.66 | 21.00 | |
| 10 | HLA-A0301 | HQYCLCRTVDGLHL | HQYCLCGLH | 0 | 0 | 0 | 6 | 4 | HQYCLCRTVDGLH | vdchain | 0.071 | 23266.15 | 23.00 | |
| 11 | HLA-A0301 | QYCLCRTVDGLHLH | CLCRTLHLH | 2 | 0 | 0 | 5 | 3 | CLCRTVDGLHLH | vdchain | 0.081 | 20782.84 | 20.00 | |
| 12 | HLA-A0301 | YCLCRTVDGLHLHR | CLCRTHLHR | 1 | 0 | 0 | 5 | 4 | CLCRTVDGLHLHR | vdchain | 0.172 | 7768.27 | 7.00 | |
| 13 | HLA-A0301 | CLCRTVDGLHLHRK | CLCRHLHRK | 0 | 0 | 0 | 4 | 5 | CLCRTVDGLHLHRK | vdchain | 0.383 | 790.12 | 1.70 | <= WB |
| 14 | HLA-A0301 | LCRTVDGLHLHRKP | RTVLHLHRK | 2 | 0 | 0 | 3 | 2 | RTVDGLHLHRK | vdchain | 0.227 | 4306.71 | 5.00 | |
| 15 | HLA-A0301 | CRTVDGLHLHRKPV | RTVLHLHRK | 1 | 0 | 0 | 3 | 2 | RTVDGLHLHRK | vdchain | 0.215 | 4884.75 | 5.00 | |
| 16 | HLA-A0301 | RTVDGLHLHRKPVK | RTLHRKPVK | 0 | 0 | 0 | 2 | 5 | RTVDGLHLHRKPVK | vdchain | 0.458 | 350.91 | 1.10 | <= WB |
| 17 | HLA-A0301 | TVDGLHLHRKPVKR | TVLHRKPVK | 0 | 0 | 0 | 2 | 4 | TVDGLHLHRKPVK | vdchain | 0.213 | 4986.15 | 5.00 | |
| 18 | HLA-A0301 | VDGLHLHRKPVKRL | GLLHRKPVK | 2 | 0 | 0 | 2 | 1 | GLHLHRKPVK | vdchain | 0.175 | 7526.60 | 7.00 | |
| 19 | HLA-A0301 | DGLHLHRKPVKRLC | GLLHRKPVK | 1 | 0 | 0 | 2 | 1 | GLHLHRKPVK | vdchain | 0.174 | 7571.93 | 7.00 | |

Protein vdchain. Allele HLA-A0301. Number of high binders 0. Number of weak binders 2. Number of peptides 20

Link to Allele Frequencies in Worldwide Populations HLA-A0301

\# Rank Threshold for Strong binding peptides 0.500
\# Rank Threshold for Weakbindingpeptides 2.000

| pos | HLA | peptide | Core | Offset | I_pos | I_len | D_pos | D_len | iCore | Identity | 1-log50k(aff) | Affinity(nM) | %Rank | BindLevel |
|---|---|---|---|---|---|---|---|---|---|---|---|---|---|---|
| 0 | HLA-A1101 | HCRCQKPCIRHQYC | CQKPIRHQY | 3 | 0 | 0 | 4 | 1 | CQKPCIRHQY | vdchain | 0.029 | 36394.57 | 55.00 | |
| 1 | HLA-A1101 | CRCQKPCIRHQYCL | CQIRHQYCL | 2 | 0 | 0 | 2 | 3 | CQKPCIRHQYCL | vdchain | 0.028 | 36735.97 | 55.00 | |
| 2 | HLA-A1101 | RCQKPCIRHQYCLC | CQKPCIHQY | 1 | 0 | 0 | 6 | 1 | CQKPCIRHQY | vdchain | 0.026 | 37561.95 | 60.00 | |

| | | | | | | | | | | | | | | |
|---|---|---|---|---|---|---|---|---|---|---|---|---|---|---|
| 3 | HLA-A1101 | CQKPCIRHQYCLCR | CQKPCILCR | 0 | 0 | 0 | 6 | 5 | CQKPCIRHQYCLCR | vdchain | 0.069 | 23807.80 | 21.00 | |
| 4 | HLA-A1101 | QKPCIRHQYCLCRT | CIHQYCLCR | 3 | 0 | 0 | 2 | 1 | CIRHQYCLCR | vdchain | 0.029 | 36578.93 | 55.00 | |
| 5 | HLA-A1101 | KPCIRHQYCLCRTV | CIHQYCLCR | 2 | 0 | 0 | 2 | 1 | CIRHQYCLCR | vdchain | 0.027 | 37379.10 | 60.00 | |
| 6 | HLA-A1101 | PCIRHQYCLCRTVD | CIHQYCLCR | 1 | 0 | 0 | 2 | 1 | CIRHQYCLCR | vdchain | 0.023 | 38822.18 | 70.00 | |
| 7 | HLA-A1101 | CIRHQYCLCRTVDG | CIHQYCLCR | 0 | 0 | 0 | 2 | 1 | CIRHQYCLCR | vdchain | 0.024 | 38490.09 | 65.00 | |
| 8 | HLA-A1101 | IRHQYCLCRTVDGL | RLCRTVDGL | 1 | 0 | 0 | 1 | 4 | RHQYCLCRTVDGL | vdchain | 0.025 | 38299.82 | 65.00 | |
| 9 | HLA-A1101 | RHQYCLCRTVDGLH | HQYTVDGLH | 1 | 0 | 0 | 3 | 4 | HQYCLCRTVDGLH | vdchain | 0.027 | 37136.41 | 60.00 | |
| 10 | HLA-A1101 | HQYCLCRTVDGLHL | HQYCLGLHL | 0 | 0 | 0 | 5 | 5 | HQYCLCRTVDGLHL | vdchain | 0.032 | 35452.82 | 50.00 | |
| 11 | HLA-A1101 | QYCLCRTVDGLHLH | QTVDGLHLH | 0 | 0 | 0 | 1 | 5 | QYCLCRTVDGLHLH | vdchain | 0.072 | 23051.67 | 20.00 | |
| 12 | HLA-A1101 | YCLCRTVDGLHLHR | RTVDGLHLR | 4 | 0 | 0 | 8 | 1 | RTVDGLHLHR | vdchain | 0.256 | 3117.33 | 4.50 | |
| 13 | HLA-A1101 | CLCRTVDGLHLHRK | RTVDGLHLK | 3 | 0 | 0 | 8 | 2 | RTVDGLHLHRK | vdchain | 0.499 | 226.59 | 1.20 | <= WB |
| 14 | HLA-A1101 | LCRTVDGLHLHRKP | RTVDGLHLK | 2 | 0 | 0 | 8 | 2 | RTVDGLHLHRK | vdchain | 0.335 | 1330.23 | 3.00 | |
| 15 | HLA-A1101 | CRTVDGLHLHRKPV | RTVDGLHLK | 1 | 0 | 0 | 8 | 2 | RTVDGLHLHRK | vdchain | 0.292 | 2131.17 | 4.00 | |
| 16 | HLA-A1101 | RTVDGLHLHRKPVK | RTVDGLHLK | 0 | 0 | 0 | 8 | 5 | RTVDGLHLHRKPVK | vdchain | 0.558 | 119.50 | 0.80 | <= WB |
| 17 | HLA-A1101 | TVDGLHLHRKPVKR | TVDGLHLVK | 0 | 0 | 0 | 7 | 4 | TVDGLHLHRKPVK | vdchain | 0.223 | 4455.01 | 5.50 | |
| 18 | HLA-A1101 | VDGLHLHRKPVKRL | GLHLHRPVK | 2 | 0 | 0 | 6 | 1 | GLHLHRKPVK | vdchain | 0.045 | 30760.33 | 34.00 | |
| 19 | HLA-A1101 | DGLHLHRKPVKRLC | GLHLHRPVK | 1 | 0 | 0 | 6 | 1 | GLHLHRKPVK | vdchain | 0.042 | 31793.05 | 37.00 | |

Protein vdchain. Allele HLA-A1101. Number of high binders 0. Number of weak binders 2. Number of peptides 20

Link to Allele Frequencies in Worldwide PopulationsHLA-A1101

\# Rank Threshold for Strong binding peptides 0.500
\# Rank Threshold for Weakbindingpeptides 2.000

| pos | HLA | peptide | Core | Offset | I_pos | I_len | D_pos | D_len | iCore | Identity | 1-log50k(aff) | Affinity(nM) | %Rank | BindLevel |
|---|---|---|---|---|---|---|---|---|---|---|---|---|---|---|
| 0 | HLA-A2301 | HCRCQKPCIRHQYC | CQKPCIQYC | 3 | 0 | 0 | 6 | 2 | CQKPCIRHQYC | vdchain | 0.013 | 43376.42 | 75.00 | |
| 1 | HLA-A2301 | CRCQKPCIRHQYCL | CQKPCIYCL | 2 | 0 | 0 | 6 | 3 | CQKPCIRHQYCL | vdchain | 0.047 | 30122.36 | 26.00 | |
| 2 | HLA-A2301 | RCQKPCIRHQYCLC | CQKPCIYCL | 1 | 0 | 0 | 6 | 3 | CQKPCIRHQYCL | vdchain | 0.036 | 33754.49 | 33.00 | |
| 3 | HLA-A2301 | CQKPCIRHQYCLCR | CQKPCIYCL | 0 | 0 | 0 | 6 | 3 | CQKPCIRHQYCL | vdchain | 0.032 | 35209.31 | 36.00 | |
| 4 | HLA-A2301 | QKPCIRHQYCLCRT | QCIRHQYCL | 0 | 0 | 0 | 1 | 2 | QKPCIRHQYCL | vdchain | 0.020 | 40444.18 | 55.00 | |
| 5 | HLA-A2301 | KPCIRHQYCLCRTV | RHQYCLCTV | 4 | 0 | 0 | 7 | 1 | RHQYCLCRTV | vdchain | 0.045 | 30639.76 | 27.00 | |
| 6 | HLA-A2301 | PCIRHQYCLCRTVD | RYCLCRTVD | 3 | 0 | 0 | 1 | 2 | RHQYCLCRTVD | vdchain | 0.045 | 30776.98 | 27.00 | |
| 7 | HLA-A2301 | CIRHQYCLCRTVDG | QYLCRTVDG | 4 | 0 | 0 | 2 | 1 | QYCLCRTVDG | vdchain | 0.053 | 28113.16 | 23.00 | |
| 8 | HLA-A2301 | IRHQYCLCRTVDGL | QYCLCRTVL | 3 | 0 | 0 | 8 | 2 | QYCLCRTVDGL | vdchain | 0.085 | 19960.15 | 14.00 | |
| 9 | HLA-A2301 | RHQYCLCRTVDGLH | QYCLCRTVL | 2 | 0 | 0 | 8 | 2 | QYCLCRTVDGL | vdchain | 0.060 | 26019.72 | 20.00 | |
| 10 | HLA-A2301 | HQYCLCRTVDGLHL | QYCLCRLHL | 1 | 0 | 0 | 6 | 4 | QYCLCRTVDGLHL | vdchain | 0.153 | 9569.98 | 7.00 | |
| 11 | HLA-A2301 | QYCLCRTVDGLHLH | QYCLCRLHL | 0 | 0 | 0 | 6 | 4 | QYCLCRTVDGLHL | vdchain | 0.114 | 14642.12 | 10.00 | |
| 12 | HLA-A2301 | YCLCRTVDGLHLHR | YCLCRTLHL | 0 | 0 | 0 | 6 | 3 | YCLCRTVDGLHL | vdchain | 0.045 | 30749.68 | 27.00 | |
| 13 | HLA-A2301 | CLCRTVDGLHLHRK | CLCRTVLHL | 0 | 0 | 0 | 6 | 2 | CLCRTVDGLHL | vdchain | 0.025 | 38287.40 | 46.00 | |
| 14 | HLA-A2301 | LCRTVDGLHLHRKP | RTVDGLHLP | 2 | 0 | 0 | 8 | 3 | RTVDGLHLHRKP | vdchain | 0.017 | 41600.80 | 65.00 | |
| 15 | HLA-A2301 | CRTVDGLHLHRKPV | RTVDGLHLV | 1 | 0 | 0 | 8 | 4 | RTVDGLHLHRKPV | vdchain | 0.030 | 36002.89 | 39.00 | |
| 16 | HLA-A2301 | RTVDGLHLHRKPVK | RTVDGLHLV | 0 | 0 | 0 | 8 | 4 | RTVDGLHLHRKPV | vdchain | 0.027 | 37405.00 | 43.00 | |
| 17 | HLA-A2301 | TVDGLHLHRKPVKR | VHLHRKPVK | 1 | 0 | 0 | 1 | 3 | VDGLHLHRKPVK | vdchain | 0.012 | 43705.71 | 75.00 | |
| 18 | HLA-A2301 | VDGLHLHRKPVKRL | LHLHRKPVL | 3 | 0 | 0 | 8 | 2 | LHLHRKPVKRL | vdchain | 0.028 | 36741.95 | 41.00 | |
| 19 | HLA-A2301 | DGLHLHRKPVKRLC | LHLHRKPVL | 2 | 0 | 0 | 8 | 2 | LHLHRKPVKRL | vdchain | 0.021 | 39789.64 | 55.00 | |

Protein vdchain. Allele HLA-A2301. Number of high binders 0. Number of weak binders 0. Number of peptides 20

Link to Allele Frequencies in Worldwide PopulationsHLA-A2301

---

### Rank Threshold for Strong binding peptides 0.500
### Rank Threshold for Weakbindingpeptides 2.000

---

| pos | HLA | peptide | CoreOffset | I_pos | I_len | D_pos | D_len | iCore | Identity | 1-log50k(aff) | Affinity(nM) | %Rank | BindLevel |
|---|---|---|---|---|---|---|---|---|---|---|---|---|---|
| 0 | HLA-A2402 | HCRCQKPCIRHQYC | RCQKRHQYC | 2 | 0 | 0 | 4 | 3 | RCQKPCIRHQYC | vdchain | 0.016 | 41953.37 | 60.00 |
| 1 | HLA-A2402 | CRCQKPCIRHQYCL | RCIRHQYCL | 1 | 0 | 0 | 1 | 4 | RCQKPCIRHQYCL | vdchain | 0.059 | 26468.64 | 19.00 |
| 2 | HLA-A2402 | RCQKPCIRHQYCLC | RCIRHQYCL | 0 | 0 | 0 | 1 | 4 | RCQKPCIRHQYCL | vdchain | 0.053 | 28167.04 | 21.00 |
| 3 | HLA-A2402 | CQKPCIRHQYCLCR | CQIRHQYCL | 0 | 0 | 0 | 2 | 3 | CQKPCIRHQYCL | vdchain | 0.042 | 31623.57 | 26.00 |
| 4 | HLA-A2402 | QKPCIRHQYCLCRT | KCIRHQYCL | 1 | 0 | 0 | 1 | 1 | KPCIRHQYCL | vdchain | 0.024 | 38443.90 | 42.00 |
| 5 | HLA-A2402 | KPCIRHQYCLCRTV | KCIRHQYCL | 0 | 0 | 0 | 1 | 1 | KPCIRHQYCL | vdchain | 0.036 | 33951.20 | 30.00 |
| 6 | HLA-A2402 | PCIRHQYCLCRTVD | RYCLCRTVD | 3 | 0 | 0 | 1 | 2 | RHQYCLCRTVD | vdchain | 0.042 | 31629.04 | 26.00 |
| 7 | HLA-A2402 | CIRHQYCLCRTVDG | RYCLCRTVD | 2 | 0 | 0 | 1 | 2 | RHQYCLCRTVD | vdchain | 0.037 | 33659.67 | 29.00 |
| 8 | HLA-A2402 | IRHQYCLCRTVDGL | QYCLCRTVL | 3 | 0 | 0 | 8 | 2 | QYCLCRTVDGL | vdchain | 0.056 | 27335.38 | 20.00 |
| 9 | HLA-A2402 | RHQYCLCRTVDGLH | RHQYCLCRL | 0 | 0 | 0 | 8 | 4 | RHQYCLCRTVDGL | vdchain | 0.048 | 29720.61 | 23.00 |
| 10 | HLA-A2402 | HQYCLCRTVDGLHL | QYCLCGLHL | 1 | 0 | 0 | 5 | 4 | QYCLCRTVDGLHL | vdchain | 0.078 | 21430.44 | 14.00 |
| 11 | HLA-A2402 | QYCLCRTVDGLHLH | QYCLCGLHL | 0 | 0 | 0 | 5 | 4 | QYCLCRTVDGLHL | vdchain | 0.075 | 22245.53 | 15.00 |
| 12 | HLA-A2402 | YCLCRTVDGLHLHR | YCLCRTLHL | 0 | 0 | 0 | 6 | 3 | YCLCRTVDGLHL | vdchain | 0.036 | 33812.99 | 30.00 |
| 13 | HLA-A2402 | CLCRTVDGLHLHRK | CLCRTVLHL | 0 | 0 | 0 | 6 | 2 | CLCRTVDGLHL | vdchain | 0.025 | 38105.95 | 40.00 |
| 14 | HLA-A2402 | LCRTVDGLHLHRKP | LCRTVGLHL | 0 | 0 | 0 | 5 | 1 | LCRTVDGLHL | vdchain | 0.013 | 43335.15 | 65.00 |
| 15 | HLA-A2402 | CRTVDGLHLHRKPV | RTVDGLHLV | 1 | 0 | 0 | 8 | 4 | RTVDGLHLHRKPV | vdchain | 0.016 | 41839.13 | 60.00 |
| 16 | HLA-A2402 | RTVDGLHLHRKPVK | RTVDGLHLV | 0 | 0 | 0 | 8 | 4 | RTVDGLHLHRKPV | vdchain | 0.016 | 41945.20 | 60.00 |
| 17 | HLA-A2402 | TVDGLHLHRKPVKR | TVDGLHLHV | 0 | 0 | 0 | 8 | 3 | TVDGLHLHRKPV | vdchain | 0.011 | 44480.95 | 75.00 |
| 18 | HLA-A2402 | VDGLHLHRKPVKRL | VDGLHLHRL | 0 | 0 | 0 | 7 | 5 | VDGLHLHRKPVKRL | vdchain | 0.026 | 37784.09 | 39.00 |
| 19 | HLA-A2402 | DGLHLHRKPVKRLC | LHLHRKPVL | 2 | 0 | 0 | 8 | 2 | LHLHRKPVKRL | vdchain | 0.017 | 41438.16 | 55.00 |

Protein vdchain. Allele HLA-A2402. Number of high binders 0. Number of weak binders 0. Number of peptides 20

Link to Allele Frequencies in Worldwide PopulationsHLA-A2402

---

**Explain**the output. Go**back**.
