## Supplementary material for "Computational methods to develop potential neutralizing antibody Fab region against SARS-CoV-2 as therapeutic and diagnostic tool": Supp-4

### Supplementary file 4

#### List of all matched Gene Ontology (GO) terms for cn40

| Type | GO no. | Description | Score | BLAST | IPR | SUP | PDB | SSM | ENZ | DNA | LIG | REV |
|---|---|---|---|---|---|---|---|---|---|---|---|---|
| Cellular component | | | | | | | | | | | | |
| | 0019814 | immunoglobulin complex | 0.90 | - | - | - | - | - | - | - | - | 0.90 (1) |
| | 0009897 | external side of plasma membrane | 0.90 | - | - | - | - | - | - | - | - | 0.90 (1) |
| | 0042571 | immunoglobulin complex\, circulating | 0.90 | - | - | - | - | - | - | - | - | 0.90 (1) |
| | 0032991 | macromolecular complex | 0.90 | - | - | - | - | - | - | - | - | 0.90 (1) |
| | 0043234 | protein complex | 0.90 | - | - | - | - | - | - | - | - | 0.90 (1) |
| | 0005623 | cell | 0.90 | - | - | - | - | - | - | - | - | 0.90 (1) |
| | 0044464 | cell part | 0.90 | - | - | - | - | - | - | - | - | 0.90 (1) |
| | 0009986 | cell surface | 0.90 | - | - | - | - | - | - | - | - | 0.90 (1) |
| | 0016020 | membrane | 0.90 | - | - | - | - | - | - | - | - | 0.90 (1) |
| | 0044425 | membrane part | 0.90 | - | - | - | - | - | - | - | - | 0.90 (1) |
| | 0044459 | plasma membrane part | 0.90 | - | - | - | - | - | - | - | - | 0.90 (1) |
| | 0005886 | plasma membrane | 0.90 | - | - | - | - | - | - | - | - | 0.90 (1) |
| | 0005576 | extracellular region | 0.90 | - | - | - | - | - | - | - | - | 0.90 (1) |
| | 0044421 | extracellular region part | 0.90 | - | - | - | - | - | - | - | - | 0.90 (1) |
| | 0005615 | extracellular space | 0.90 | - | - | - | - | - | - | - | - | 0.90 (1) |
| Biological process | | | | | | | | | | | | |
| | 0002250 | adaptive immune response | 0.90 | - | - | - | - | - | - | - | - | 0.90 (1) |
| | 0002376 | immune system process | 0.90 | - | - | - | - | - | - | - | - | 0.90 (1) |
| | 0006910 | phagocytosis\, recognition | 0.90 | - | - | - | - | - | - | - | - | 0.90 (1) |
| | 0006911 | phagocytosis\, engulfment | 0.90 | - | - | - | - | - | - | - | - | 0.90 (1) |
| | 0006958 | complement activation\, classical pathway | 0.90 | - | - | - | - | - | - | - | - | 0.90 (1) |
| | 0042742 | defense response to bacterium | 0.90 | - | - | - | - | - | - | - | - | 0.90 (1) |
| | 0045087 | innate immune response | 0.90 | - | - | - | - | - | - | - | - | 0.90 (1) |
| | 0050853 | B cell receptor signaling pathway | 0.90 | - | - | - | - | - | - | - | - | 0.90 (1) |
| | 0050871 | positive regulation of B cell activation | 0.90 | - | - | - | - | - | - | - | - | 0.90 (1) |
| | 0006955 | immune response | 0.90 | - | - | - | - | - | - | - | - | 0.90 (1) |
| | 0050896 | response to stimulus | 0.90 | - | - | - | - | - | - | - | - | 0.90 (1) |
| | 0016043 | cellular component organization | 0.90 | - | - | - | - | - | - | - | - | 0.90 (1) |
| | 0016044 | membrane organization | 0.90 | - | - | - | - | - | - | - | - | 0.90 (1) |
| | 0010324 | membrane invagination | 0.90 | - | - | - | - | - | - | - | - | 0.90 (1) |
| | 0006897 | endocytosis | 0.90 | - | - | - | - | - | - | - | - | 0.90 (1) |
| | 0006909 | phagocytosis | 0.90 | - | - | - | - | - | - | - | - | 0.90 (1) |
| | 0009987 | cellular process | 0.90 | - | - | - | - | - | - | - | - | 0.90 (1) |
| | 0008037 | cell recognition | 0.90 | - | - | - | - | - | - | - | - | 0.90 (1) |
| | 0016192 | vesicle\-mediated transport | 0.90 | - | - | - | - | - | - | - | - | 0.90 (1) |
| | 0051234 | establishment of localization | 0.90 | - | - | - | - | - | - | - | - | 0.90 (1) |
| | 0006810 | transport | 0.90 | - | - | - | - | - | - | - | - | 0.90 (1) |
| | 0051179 | localization | 0.90 | - | - | - | - | - | - | - | - | 0.90 (1) |
| | 0065007 | biological regulation | 0.90 | - | - | - | - | - | - | - | - | 0.90 (1) |
| | 0050789 | regulation of biological process | 0.90 | - | - | - | - | - | - | - | - | 0.90 (1) |
| | 0048518 | positive regulation of biological process | 0.90 | - | - | - | - | - | - | - | - | 0.90 (1) |
| | 0002684 | positive regulation of immune system process | 0.90 | - | - | - | - | - | - | - | - | 0.90 (1) |
| | 0050778 | positive regulation of immune response | 0.90 | - | - | - | - | - | - | - | - | 0.90 (1) |
| | 0002253 | activation of immune response | 0.90 | - | - | - | - | - | - | - | - | 0.90 (1) |
| | 0006956 | complement activation | 0.90 | - | - | - | - | - | - | - | - | 0.90 (1) |
| | 0048584 | positive regulation of response to stimulus | 0.90 | - | - | - | - | - | - | - | - | 0.90 (1) |
| | 0002682 | regulation of immune system process | 0.90 | - | - | - | - | - | - | - | - | 0.90 (1) |
| | 0050776 | regulation of immune response | 0.90 | - | - | - | - | - | - | - | - | 0.90 (1) |
| | 0048583 | regulation of response to stimulus | 0.90 | - | - | - | - | - | - | - | - | 0.90 (1) |

| | | | | | | | | | | | |
|---|---|---|---|---|---|---|---|---|---|---|---|
| 0002252 | immune effector process | 0.90 | - | - | - | - | - | - | - | - | 0.90 (1) |
| 0002443 | leukocyte mediated immunity | 0.90 | - | - | - | - | - | - | - | - | 0.90 (1) |
| 0002449 | lymphocyte mediated immunity | 0.90 | - | - | - | - | - | - | - | - | 0.90 (1) |
| 0019724 | B cell mediated immunity | 0.90 | - | - | - | - | - | - | - | - | 0.90 (1) |
| 0016064 | immunoglobulin mediated immune response | 0.90 | - | - | - | - | - | - | - | - | 0.90 (1) |
| 0002455 | humoral immune response mediated by circulating immunoglobulin | 0.90 | - | - | - | - | - | - | - | - | 0.90 (1) |
| 0002460 | adaptive immune response based on somatic recombination of immune receptors built from immunoglobulin superfamily domains | 0.90 | - | - | - | - | - | - | - | - | 0.90 (1) |
| 0006959 | humoral immune response | 0.90 | - | - | - | - | - | - | - | - | 0.90 (1) |
| 0008152 | metabolic process | 0.90 | - | - | - | - | - | - | - | - | 0.90 (1) |
| 0043170 | macromolecule metabolic process | 0.90 | - | - | - | - | - | - | - | - | 0.90 (1) |
| 0043283 | biopolymer metabolic process | 0.90 | - | - | - | - | - | - | - | - | 0.90 (1) |
| 0019538 | protein metabolic process | 0.90 | - | - | - | - | - | - | - | - | 0.90 (1) |
| 0051604 | protein maturation | 0.90 | - | - | - | - | - | - | - | - | 0.90 (1) |
| 0016485 | protein processing | 0.90 | - | - | - | - | - | - | - | - | 0.90 (1) |
| 0051605 | protein maturation by peptide bond cleavage | 0.90 | - | - | - | - | - | - | - | - | 0.90 (1) |
| 0002541 | activation of plasma proteins involved in acute inflammatory response | 0.90 | - | - | - | - | - | - | - | - | 0.90 (1) |
| 0010467 | gene expression | 0.90 | - | - | - | - | - | - | - | - | 0.90 (1) |
| 0044238 | primary metabolic process | 0.90 | - | - | - | - | - | - | - | - | 0.90 (1) |
| 0009605 | response to external stimulus | 0.90 | - | - | - | - | - | - | - | - | 0.90 (1) |
| 0009611 | response to wounding | 0.90 | - | - | - | - | - | - | - | - | 0.90 (1) |
| 0006954 | inflammatory response | 0.90 | - | - | - | - | - | - | - | - | 0.90 (1) |
| 0002526 | acute inflammatory response | 0.90 | - | - | - | - | - | - | - | - | 0.90 (1) |
| 0006950 | response to stress | 0.90 | - | - | - | - | - | - | - | - | 0.90 (1) |
| 0006952 | defense response | 0.90 | - | - | - | - | - | - | - | - | 0.90 (1) |
| 0051704 | multi\-organism process | 0.90 | - | - | - | - | - | - | - | - | 0.90 (1) |
| 0051707 | response to other organism | 0.90 | - | - | - | - | - | - | - | - | 0.90 (1) |
| 0009617 | response to bacterium | 0.90 | - | - | - | - | - | - | - | - | 0.90 (1) |
| 0009607 | response to biotic stimulus | 0.90 | - | - | - | - | - | - | - | - | 0.90 (1) |
| 0002757 | immune response\-activating signal transduction | 0.90 | - | - | - | - | - | - | - | - | 0.90 (1) |
| 0002429 | immune response\-activating cell surface receptor signaling pathway | 0.90 | - | - | - | - | - | - | - | - | 0.90 (1) |
| 0050851 | antigen receptor\-mediated signaling pathway | 0.90 | - | - | - | - | - | - | - | - | 0.90 (1) |
| 0050794 | regulation of cellular process | 0.90 | - | - | - | - | - | - | - | - | 0.90 (1) |
| 0007165 | signal transduction | 0.90 | - | - | - | - | - | - | - | - | 0.90 (1) |
| 0007166 | cell surface receptor linked signal transduction | 0.90 | - | - | - | - | - | - | - | - | 0.90 (1) |
| 0002768 | immune response\-regulating cell surface receptor signaling pathway | 0.90 | - | - | - | - | - | - | - | - | 0.90 (1) |
| 0002764 | immune response\-regulating signal transduction | 0.90 | - | - | - | - | - | - | - | - | 0.90 (1) |
| 0007154 | cell communication | 0.90 | - | - | - | - | - | - | - | - | 0.90 (1) |
| 0048522 | positive regulation of cellular process | 0.90 | - | - | - | - | - | - | - | - | 0.90 (1) |
| 0050867 | positive regulation of cell activation | 0.90 | - | - | - | - | - | - | - | - | 0.90 (1) |
| 0002696 | positive regulation of leukocyte activation | 0.90 | - | - | - | - | - | - | - | - | 0.90 (1) |
| 0051251 | positive regulation of lymphocyte activation | 0.90 | - | - | - | - | - | - | - | - | 0.90 (1) |
| 0050865 | regulation of cell activation | 0.90 | - | - | - | - | - | - | - | - | 0.90 (1) |
| 0002694 | regulation of leukocyte | 0.90 | - | - | - | - | - | - | - | - | 0.90 (1) |

| | | | | | | | | | | | |
|---|---|---|---|---|---|---|---|---|---|---|---|
| | activation | | | | | | | | | | |
| 0051249 | regulation of lymphocyte activation | 0.90 | - | - | - | - | - | - | - | - | 0.90 (1) |
| 0050864 | regulation of B cell activation | 0.90 | - | - | - | - | - | - | - | - | 0.90 (1) |
| 0001775 | cell activation | 0.90 | - | - | - | - | - | - | - | - | 0.90 (1) |
| 0045321 | leukocyte activation | 0.90 | - | - | - | - | - | - | - | - | 0.90 (1) |
| 0046649 | lymphocyte activation | 0.90 | - | - | - | - | - | - | - | - | 0.90 (1) |
| 0042113 | B cell activation | 0.90 | - | - | - | - | - | - | - | - | 0.90 (1) |

**Biochemical function**

| | | | | | | | | | | | |
|---|---|---|---|---|---|---|---|---|---|---|---|
| 0003823 | antigen binding | 0.90 | - | - | - | - | - | - | - | - | 0.90 (1) |
| 0034987 | immunoglobulin receptor binding | 0.90 | - | - | - | - | - | - | - | - | 0.90 (1) |
| 0005488 | binding | 0.90 | - | - | - | - | - | - | - | - | 0.90 (1) |
| 0005515 | protein binding | 0.90 | - | - | - | - | - | - | - | - | 0.90 (1) |
| 0005102 | receptor binding | 0.90 | - | - | - | - | - | - | - | - | 0.90 (1) |

The score in red is a measure of how strongly the term is predicted from the hits obtained by the different methods. The scores in blue show each method's contribution to the total score (with the number of relevant sequences/structures shown in brackets in grey). Click on any score to see the list of hits from the given program and the source of the given term.

#### List of all matched protein name terms for cn40

| Terms extracted from protein names | Score | BLAST | IPR | SUP | PDB | SSM | ENZ | DNA | LIG | REV |
|---|---|---|---|---|---|---|---|---|---|---|
| fab | 6.31 | - | - | - | - | - | - | - | - | 6.31 (11) |
| antibody | 4.99 | - | - | - | - | - | - | - | - | 4.99 (12) |
| fragment | 4.80 | - | - | - | - | - | - | - | - | 4.80 (9) |
| fab fragment | 3.77 | - | - | - | - | - | - | - | - | 3.77 (6) |
| human | 1.91 | - | - | - | - | - | - | - | - | 1.91 (3) |
| ige | 1.80 | - | - | - | - | - | - | - | - | 1.80 (2) |
| neutralizing antibody | 1.57 | - | - | - | - | - | - | - | - | 1.57 (2) |
| peptide | 1.57 | - | - | - | - | - | - | - | - | 1.57 (2) |
| specific | 1.54 | - | - | - | - | - | - | - | - | 1.54 (2) |
| rabbit | 1.47 | - | - | - | - | - | - | - | - | 1.47 (3) |
| murine | 1.19 | - | - | - | - | - | - | - | - | 1.19 (4) |
| conformation | 1.18 | - | - | - | - | - | - | - | - | 1.18 (2) |
| germline | 1.14 | - | - | - | - | - | - | - | - | 1.14 (4) |
| germline antibody | 1.00 | - | - | - | - | - | - | - | - | 1.00 (2) |
| anti- 4-hydroxy-3- nitrophenyl -acetyl | 0.97 | - | - | - | - | - | - | - | - | 0.97 (3) |
| 4-hydroxy-3- nitrophenyl -acetyl murine | 0.97 | - | - | - | - | - | - | - | - | 0.97 (3) |
| nitrophenyl -acetyl murine germline | 0.97 | - | - | - | - | - | - | - | - | 0.97 (3) |
| antibody bbe6 12h3 | 0.97 | - | - | - | - | - | - | - | - | 0.97 (3) |
| -acetyl murine germline monoclonal | 0.95 | - | - | - | - | - | - | - | - | 0.95 (2) |
| murine germline monoclonal antibody | 0.95 | - | - | - | - | - | - | - | - | 0.95 (2) |
| germline monoclonal antibody bbe6 | 0.95 | - | - | - | - | - | - | - | - | 0.95 (2) |
| monoclonal antibody bbe6 12h3 | 0.95 | - | - | - | - | - | - | - | - | 0.95 (2) |
| antibody bbe6 12h3 fab | 0.95 | - | - | - | - | - | - | - | - | 0.95 (2) |
| bbe6 12h3 fab fragment | 0.95 | - | - | - | - | - | - | - | - | 0.95 (2) |
| phage display derived dodecapeptide | 0.95 | - | - | - | - | - | - | - | - | 0.95 (2) |
| implications | 0.90 | - | - | - | - | - | - | - | - | 0.90 (1) |
| hiv-1 rev dimer | 0.90 | - | - | - | - | - | - | - | - | 0.90 (1) |
| multimeric binding | 0.90 | - | - | - | - | - | - | - | - | 0.90 (1) |
| response element | 0.90 | - | - | - | - | - | - | - | - | 0.90 (1) |
| calcium-activated chloride channel bestrophin-1 | 0.90 | - | - | - | - | - | - | - | - | 0.90 (1) |
| chicken | 0.90 | - | - | - | - | - | - | - | - | 0.90 (1) |
| fab antibody fragments | 0.90 | - | - | - | - | - | - | - | - | 0.90 (1) |
| free conformation ab1 | 0.90 | - | - | - | - | - | - | - | - | 0.90 (1) |
| ige spe-7 | 0.90 | - | - | - | - | - | - | - | - | 0.90 (1) |
| trivalent antibody fragment | 0.90 | - | - | - | - | - | - | - | - | 0.90 (1) |

| | | | | | | | | | | |
|---|---|---|---|---|---|---|---|---|---|---|
| display derived dodecapeptide yqlrpnaetlrf | 0.90 | - | - | - | - | - | - | - | - | 0.90 (1) |
| display derived dodecapeptide ppypawhapgni | 0.90 | - | - | - | - | - | - | - | - | 0.90 (1) |
| fab fragment anti- 4-hydroxy-3- | 0.90 | - | - | - | - | - | - | - | - | 0.90 (1) |
| fragment anti- 4-hydroxy-3-nitrophenyl | 0.90 | - | - | - | - | - | - | - | - | 0.90 (1) |
| -acetyl murine germline antibody | 0.90 | - | - | - | - | - | - | - | - | 0.90 (1) |
| murine germline antibody bbe6 | 0.90 | - | - | - | - | - | - | - | - | 0.90 (1) |
| germline antibody bbe6 12h3 | 0.90 | - | - | - | - | - | - | - | - | 0.90 (1) |
| b1-8 fragment | 0.90 | - | - | - | - | - | - | - | - | 0.90 (1) |
| 4-hydroxy-3-nitrophenyl acetate compound | 0.90 | - | - | - | - | - | - | - | - | 0.90 (1) |
| fab-fragment enantioselective antibody | 0.90 | - | - | - | - | - | - | - | - | 0.90 (1) |
| finrozole | 0.90 | - | - | - | - | - | - | - | - | 0.90 (1) |
| peptide gdprpsyishll | 0.90 | - | - | - | - | - | - | - | - | 0.90 (1) |
| rabbit mab r20 fab | 0.90 | - | - | - | - | - | - | - | - | 0.90 (1) |
| antigen-binding fragment | 0.90 | - | - | - | - | - | - | - | - | 0.90 (1) |
| murine anti-phenylarsonate antibody | 0.90 | - | - | - | - | - | - | - | - | 0.90 (1) |
| "fab | 0.90 | - | - | - | - | - | - | - | - | 0.90 (1) |
| human prostate specific antigen | 0.90 | - | - | - | - | - | - | - | - | 0.90 (1) |
| prostate specific antigen psa | 0.90 | - | - | - | - | - | - | - | - | 0.90 (1) |
| specific antigen psa fab | 0.90 | - | - | - | - | - | - | - | - | 0.90 (1) |
| antigen psa fab sandwich | 0.90 | - | - | - | - | - | - | - | - | 0.90 (1) |
| high affinity | 0.90 | - | - | - | - | - | - | - | - | 0.90 (1) |
| pca selective antibody | 0.90 | - | - | - | - | - | - | - | - | 0.90 (1) |
| conformation-dependent rabbit igg fab | 0.90 | - | - | - | - | - | - | - | - | 0.90 (1) |
| rabbit igg fab specific | 0.90 | - | - | - | - | - | - | - | - | 0.90 (1) |
| amyloid prefibrillar oligomers | 0.90 | - | - | - | - | - | - | - | - | 0.90 (1) |
| human tweak | 0.90 | - | - | - | - | - | - | - | - | 0.90 (1) |
| fab fragment neutralizing antibody | 0.90 | - | - | - | - | - | - | - | - | 0.90 (1) |
| anti-hinge rabbit antibody | 0.90 | - | - | - | - | - | - | - | - | 0.90 (1) |
| ides hinge peptide | 0.90 | - | - | - | - | - | - | - | - | 0.90 (1) |
| leut | 0.90 | - | - | - | - | - | - | - | - | 0.90 (1) |
| inward-open conformation | 0.90 | - | - | - | - | - | - | - | - | 0.90 (1) |
| recombinant ige fab fragment | 0.90 | - | - | - | - | - | - | - | - | 0.90 (1) |
| bovine beta-lactoglobulin allergen | 0.90 | - | - | - | - | - | - | - | - | 0.90 (1) |
| influenza virus hemagglutinin | 0.90 | - | - | - | - | - | - | - | - | 0.90 (1) |
| tr1 fab fragment human | 0.90 | - | - | - | - | - | - | - | - | 0.90 (1) |
| fab fragment human igg1 | 0.90 | - | - | - | - | - | - | - | - | 0.90 (1) |
| fragment human igg1 kappa | 0.90 | - | - | - | - | - | - | - | - | 0.90 (1) |

| | | | | | | | | | | |
|---|---|---|---|---|---|---|---|---|---|---|
| **human igg1 kappa autoantibody** | 0.90 | - | - | - | - | - | - | - | - | 0.90 (1) |

The score in red is a measure of how strongly the term is predicted from the hits obtained by the different methods. The scores in blue show each method's contribution to the total score (with the number of relevant sequences/structures shown in brackets in grey). Click on any score to see the list of hits from the given program and the source of the given term.

Binding sites

### Binding site analysis

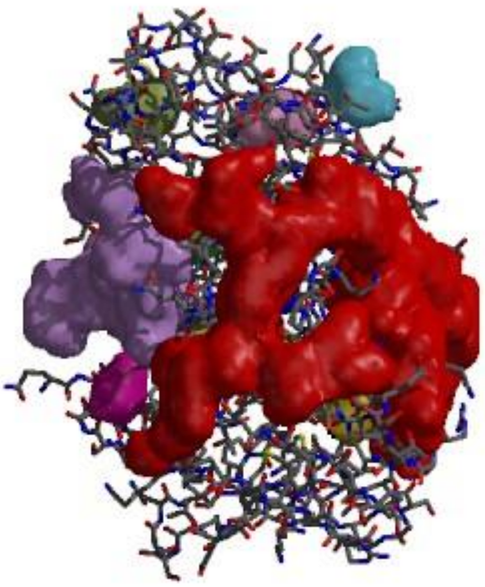

*Clefts and cavities in protein structure,*
*the colours corresponding to the entries in the table below.*

The following table shows the "gap" regions (clefts and cavities) in the protein's surface. The gaps are ordered by decreasing volume (in Å$^3$), and the table includes various parameters that are indicative of binding sites. (Ignore data shown in grey).

| Gap region | Volume | R1 ratio | Accessible vertices | | Buried vertices | | Ave. depth | | Residue type* | Residue conservation ** | Ligands |
|---|---|---|---|---|---|---|---|---|---|---|---|
| ☑ 1 | 3825.56 | 2.80 | 65.37 % | 4 | 10.66 % | 2 | 13.70 | 1 | 21 1<br>874471<br>. | . . . . . . . . .<br>. | |

| | | | | | | | | | | |
|---|---|---|---|---|---|---|---|---|---|---|
| ☑ 2 | 1364.34 | - | 63.66% | 5 | 9.44% | 3 | 9.75 | 2 | 336575. | .......... |
| ☑ 3 | 410.48 | - | 61.17% | 6 | 3.21% | 8 | 8.06 | 3 | .11341. | .......... |
| ☑ 4 | 210.94 | - | 97.70% | 1 | 13.03% | 1 | 0.00 | 9 | .23263. | .......... |
| ☐ 5 | 239.62 | - | 58.44% | 9 | 7.48% | 4 | 5.71 | 4 | .1224.. | .......... |
| ☐ 6 | 211.36 | - | 54.39% | 10 | 2.67% | 9 | 5.47 | 5 | 112211. | .......... |
| ☐ 7 | 137.53 | - | 59.18% | 8 | 4.08% | 6 | 5.31 | 6 | .14112. | .......... |
| ☐ 8 | 125.30 | - | 66.67% | 3 | 4.30% | 5 | 0.00 | 10 | ..3311. | .......... |
| ☐ 9 | 119.39 | - | 59.36% | 7 | 1.17% | 10 | 5.06 | 8 | ..3211. | .......... |
| ☐ 10 | 128.67 | - | 69.52% | 2 | 3.21% | 7 | 5.27 | 7 | 113...1 | .......... |

**Total number of gap regions analysed: 10**

##### Key to residue colour codes

| * Residue-type colouring: | | | | | | |
|---|---|---|---|---|---|---|
| Positive | Negative | Neutral | Aliphatic | Aromatic | Pro & Gly | Cysteine |
| H,K,R | D,E | S,T,N,Q | A,V,L,I,M | F,Y,W | P,G | C |

| ** Colouring by residue-conservation score: | | | | | | | | | |
|---|---|---|---|---|---|---|---|---|---|
| blue | purple | skyblue | cyan | grey | green | yellow | orange | pink | Red |
| 0.0-0.1 | 0.1-0.2 | 0.2-0.3 | 0.3-0.4 | 0.4-0.5 | 0.5-0.6 | 0.6-0.7 | 0.7-0.9 | 0.8-0.9 | 0.9-1.0 |

#### Reverse template search results for cn40

Below are the results reported by the template search program after scanning 400 auto-generated templates from the query structure against representative structures in the PDB. Click on the coloured ball or hit number at the left of each hit to see further details of the template match, together with a sequence alignment based on it.

The top 20 hits are listed below.

### a. Certain matches (E-value < 1.00E-06)

| Hit no. | E-value | Similarity score | Neighbours ident/simil [equiv] | Template id | Matched PDB entry | Longest fitted segment | Seq lengths query/target | Overlap | %-tage seq id | Structural similarity |
|---|---|---|---|---|---|---|---|---|---|---|
| 1 | 0.00E+00 | 940.00 | 48/0 [48] | TMP00336 | 2x7l×53: Implications of the HIV-1 rev dimer structure at 3.2a resolution for multimeric binding to the rev response element | 109/109 | 110/217 | 110 | 100.00% | 99.1% |
| 2 | 0.00E+00 | 920.00 | 47/0 [47] | TMP00351 | 4rdq×64: Calcium-activated chloride channel bestrophin-1, from chicken, in complex with fab antibody fragments, chloride and calcium | 115/115 | 116/211 | 116 | 100.00% | 99.1% |
| 3 | 0.00E+00 | 835.56 | 42/2 [44] | TMP00351 | 1oaq×53: Free conformation ab1 of the ige spe-7 | 115/115 | 116/120 | 120 | 81.03% | 99.1% |
| 4 | 0.00E+00 | 835.56 | 42/2 [44] | TMP00351 | 1nqb×35: Trivalent antibody fragment | 114/114 | 116/232 | 119 | 80.17% | 99.1% |
| 5 | 0.00E+00 | 834.31 | 41/4 [45] | TMP00351 | 2xzq×56: Crystal structure analysis of the anti-(4-hydroxy-3-nitrophenyl)-acetyl murine germline monoclonal antibody bbe6.12h3 fab fragment in complex with a phage display derived dodecapeptide yqlrpnaetlrf | 115/115 | 116/215 | 120 | 81.03% | 99.1% |
| 6 | 0.00E+00 | 824.94 | 41/3 [44] | TMP00351 | 2y07×58: Crystal structure analysis of the anti-(4-hydroxy-3-nitrophenyl) -acetyl murine germline monoclonal antibody bbe6.12h3 fab fragment in complex with a phage display derived dodecapeptide ppypawhapgni | 115/115 | 116/215 | 120 | 79.31% | 99.1% |
| 7 | 0.00E+00 | 824.94 | 41/3 [44] | TMP00351 | 4a6y×30: Crystal structure of fab fragment of anti-(4-hydroxy-3-nitrophenyl)-acetyl murine germline antibody bbe6.12h3 | 115/115 | 116/215 | 120 | 79.31% | 99.1% |
| 8 | 0.00E+00 | 824.94 | 41/3 [44] | TMP00351 | 1a6v×29: B1-8 fv fragment complexed with a (4-hydroxy-3-nitrophenyl) acetate | 113/113 | 116/118 | 118 | 78.45% | 99.1% |

| | | | | | | | | | | |
|---|---|---|---|---|---|---|---|---|---|---|
| | | | | | compound | | | | | |
| 9 | 0.00E+00 | 821.88 | 39/6 [45] | TMP00251 | 2c1p×63: Fab-fragment of enantioselective antibody complexed with finrozole | 114/114 | 116/217 | 115 | 75.00% | 99.1% |
| 10 | 0.00E+00 | 798.00 | 39/4 [43] | TMP00071 | 4bh8×58: Crystal structure of germline antibody 36-65 in complex with peptide gdprpsyishll | 115/115 | 116/218 | 121 | 78.45% | 99.1% |
| 11 | 0.00E+00 | 793.91 | 39/3 [42] | TMP00372 | 4jo4×62: Crystal structure of rabbit mab r20 fab | 109/109 | 110/214 | 110 | 71.82% | 99.1% |
| 12 | 0.00E+00 | 788.62 | 39/3 [42] | TMP00071 | 1jfq×71: Antigen-binding fragment of the murine anti-phenylarsonate antibody 36-71, "fab 36-71" | 114/114 | 116/220 | 120 | 76.72% | 99.1% |
| 13 | 0.00E+00 | 786.38 | 40/1 [41] | TMP00251 | 3qum×115: Crystal structure of human prostate specific antigen (psa) in fab sandwich with a high affinity and a pca selective antibody | 115/115 | 116/219 | 117 | 76.72% | 99.1% |
| 14 | 0.00E+00 | 782.16 | 38/4 [42] | TMP00372 | 4hbc×58: Crystal structure of a conformation-dependent rabbit igg fab specific for amyloid prefibrillar oligomers | 108/108 | 110/213 | 110 | 70.91% | 99.1% |
| 15 | 0.00E+00 | 781.22 | 38/4 [42] | TMP00372 | 4ht1×63: Human tweak in complex with the fab fragment of a neutralizing antibody | 107/107 | 110/217 | 113 | 70.91% | 99.1% |
| 16 | 0.00E+00 | 780.19 | 36/8 [44] | TMP00336 | 4o4y×53: Crystal structure of the anti-hinge rabbit antibody 2095-2 in complex with ides hinge peptide | 108/108 | 110/219 | 112 | 68.18% | 99.1% |
| 17 | 0.00E+00 | 776.94 | 37/6 [43] | TMP00251 | 3tt3×56: Crystal structure of leut in the inward-open conformation in complex with fab | 115/115 | 116/218 | 118 | 73.28% | 99.1% |
| 18 | 0.00E+00 | 772.97 | 36/7 [43] | TMP00336 | 2r56×53: Crystal structure of a recombinant ige fab fragment in complex with bovine beta-lactoglobulin allergen | 106/106 | 110/211 | 110 | 66.36% | 99.1% |
| 19 | 0.00E+00 | 768.50 | 37/5 [42] | TMP00071 | 1eo8×63: Influenza virus hemagglutinin complexed with a neutralizing antibody | 115/115 | 116/217 | 118 | 64.66% | 99.1% |
| 20 | 0.00E+00 | 758.75 | 36/6 [42] | TMP00336 | 1vge×55: Tr1.9 fab fragment of a human igg1 kappa autoantibody | 106/106 | 110/214 | 110 | 66.36% | 99.1% |
