## Supplementary material for "Computational methods to develop potential neutralizing antibody Fab region against SARS-CoV-2 as therapeutic and diagnostic tool": Supp-5

### Supplementary file 5

#### 1. Ramachandran Plot statistics

| | | No. of residues | %-tage |
|---|---|---|---|
| Most favoured regions | [A,B,L] | 172 | 90.5% |
| Additional allowed regions | [a,b,l,p] | 15 | 7.9% |
| Generously allowed regions | [~a,~b,~l,~p] | 2 | 1.1% |
| Disallowed regions | [XX] | 1 | 0.5%* |
| Non-glycine and non-proline residues | | 190 | 100.0% |
| End-residues (excl. Gly and Pro) | | 4 | |
| Glycine residues | | 21 | |
| Proline residues | | 11 | |
| Total number of residues | | 226 | |

Based on an analysis of **118** structures of resolution of at least **2.0** Angstroms and *R*-factor no greater than **20.0** a good quality model would be expected to have over **90%** in the most favoured regions [A,B,L].

#### 2. G-Factors

| Parameter | Score | Average Score |
|---|---|---|
| Dihedral angles:- | | |
| Phi-psi distribution | -0.48 | |
| Chi1-chi2 distribution | -0.15 | |
| Chi1 only | 0.03 | |
| Chi3 & chi4 | 0.55 | |
| Omega | 0.16 | |
| | | -0.07 |
| Main-chain covalent forces:- | | |
| Main-chain bond lengths | 0.55 | |
| Main-chain bond angles | 0.39 | |
| | | 0.46 |
| OVERALL AVERAGE | | 0.14 |

**G-factors** provide a measure of how **unusual**, or out-of-the-ordinary, a property is.
Values below -0.5* - unusual
Values below **-1.0**** - highly unusual
**Important note:** The main-chain bond-lengths and bond angles are compared with the Engh & Huber (1991) ideal values derived from small-molecule data. Therefore, structures refined using different restraints may show apparently large deviations from normality.

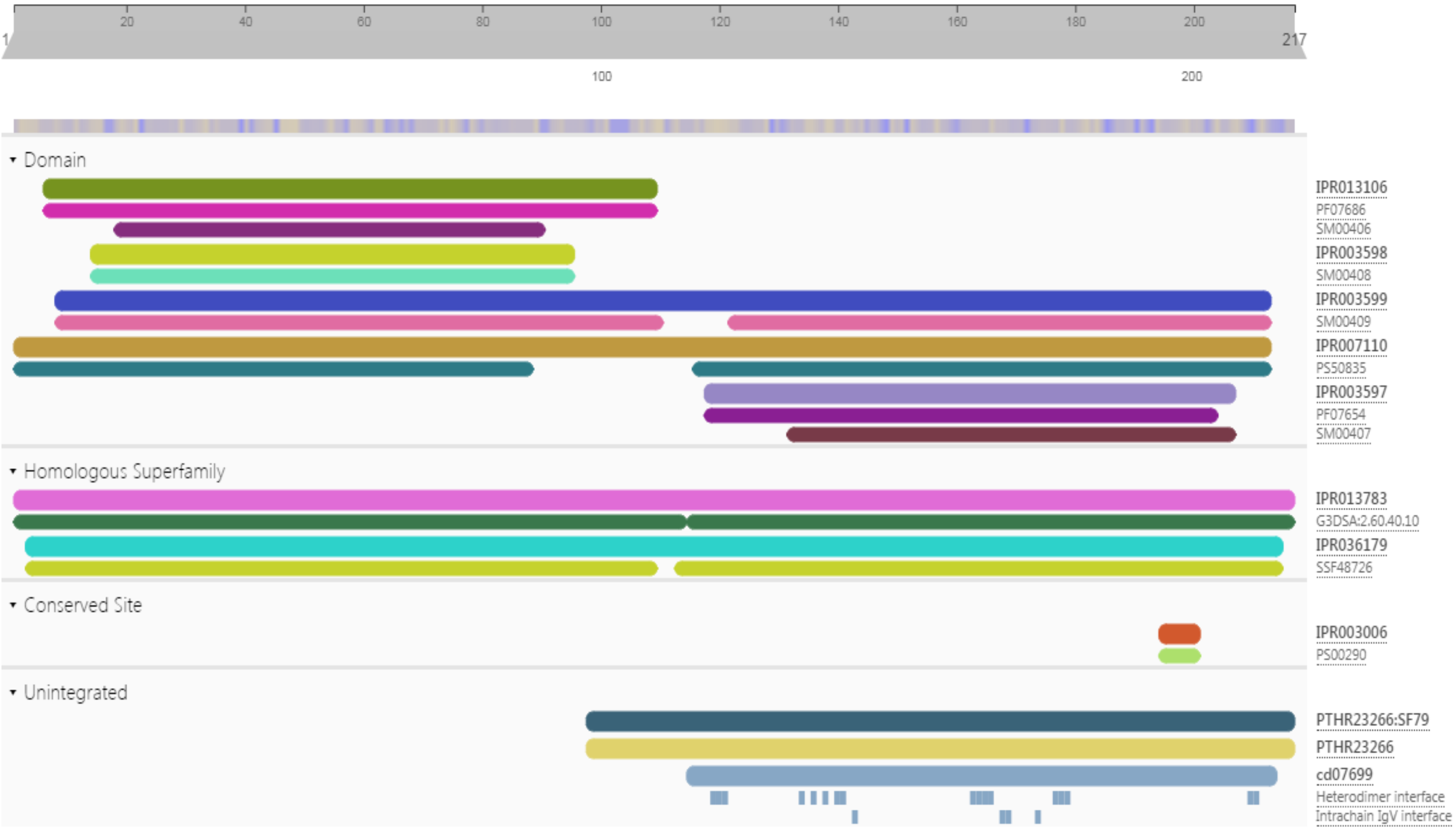


Heavy chain superfamily analysis

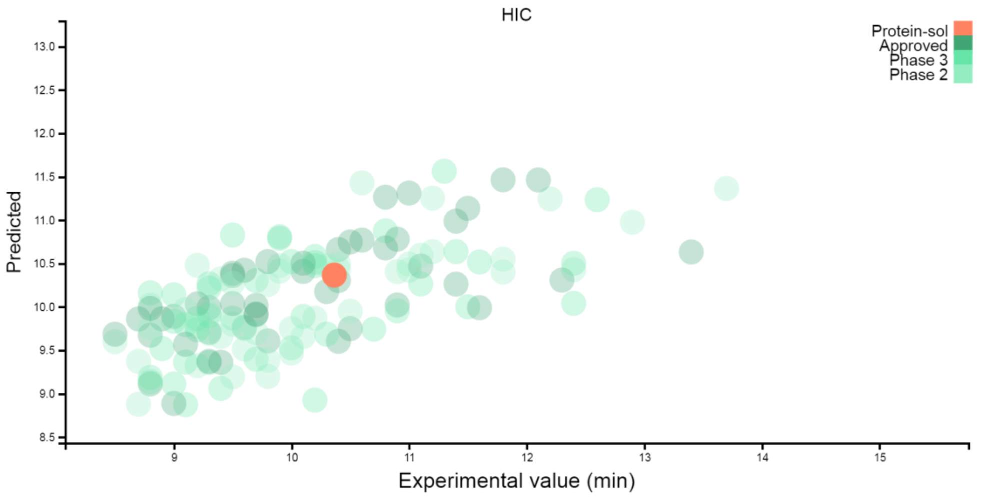


HIC PLOT FOR HEAVY CHAIN

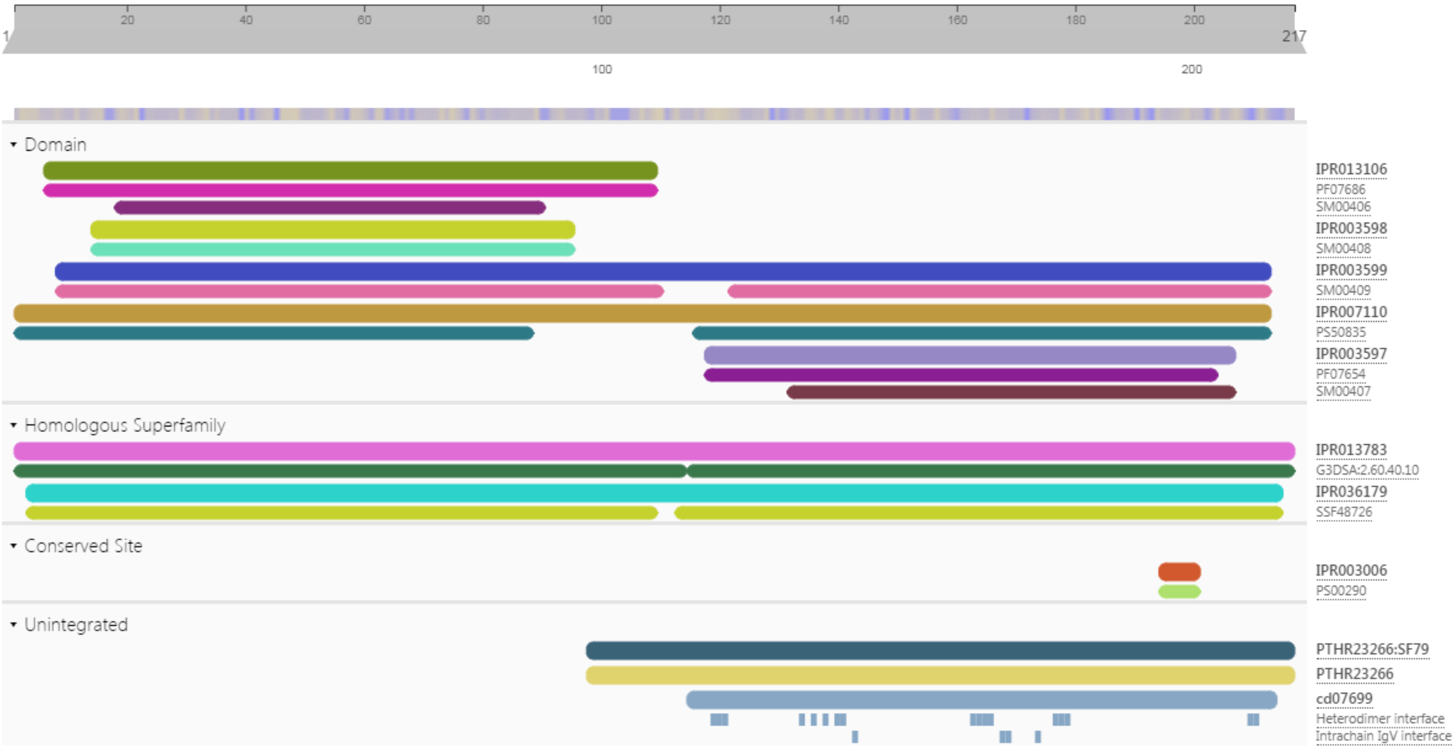


Light chain superfamily analysis

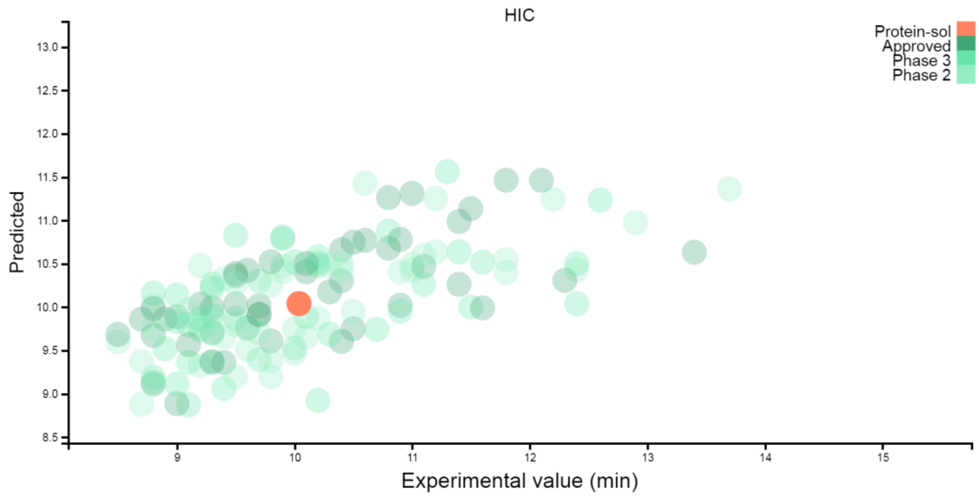


LIGHT CHAIN HIC PLOT

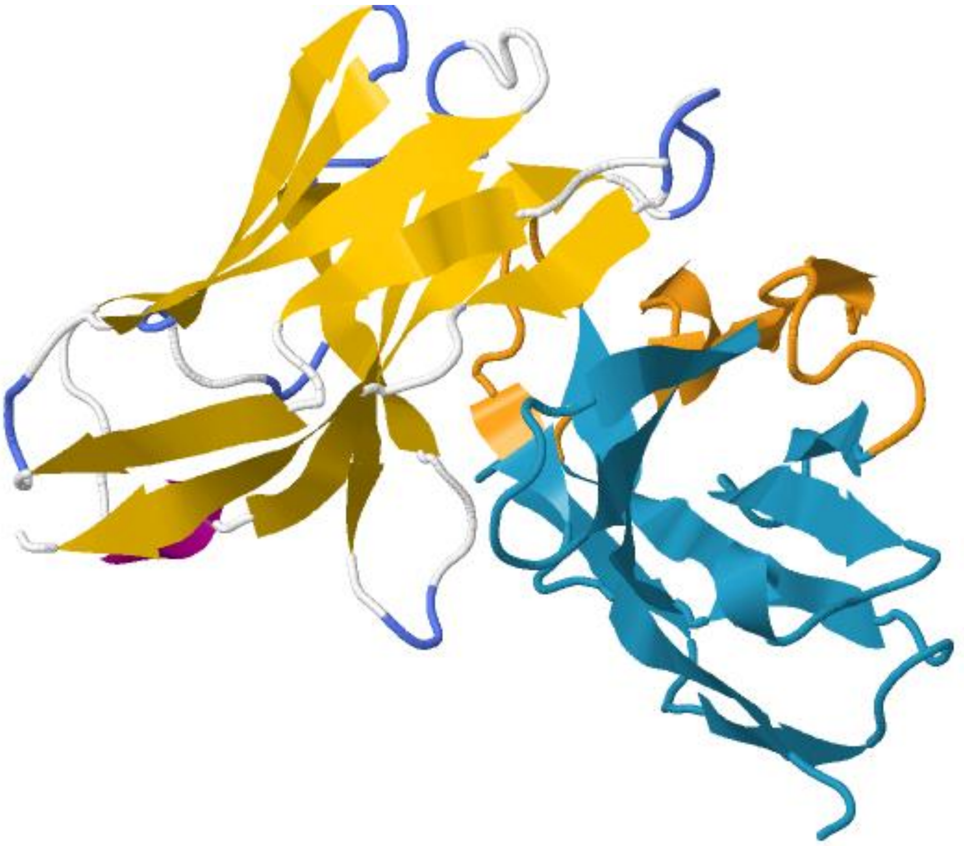

Final confirmed structure of FAB region of antibody

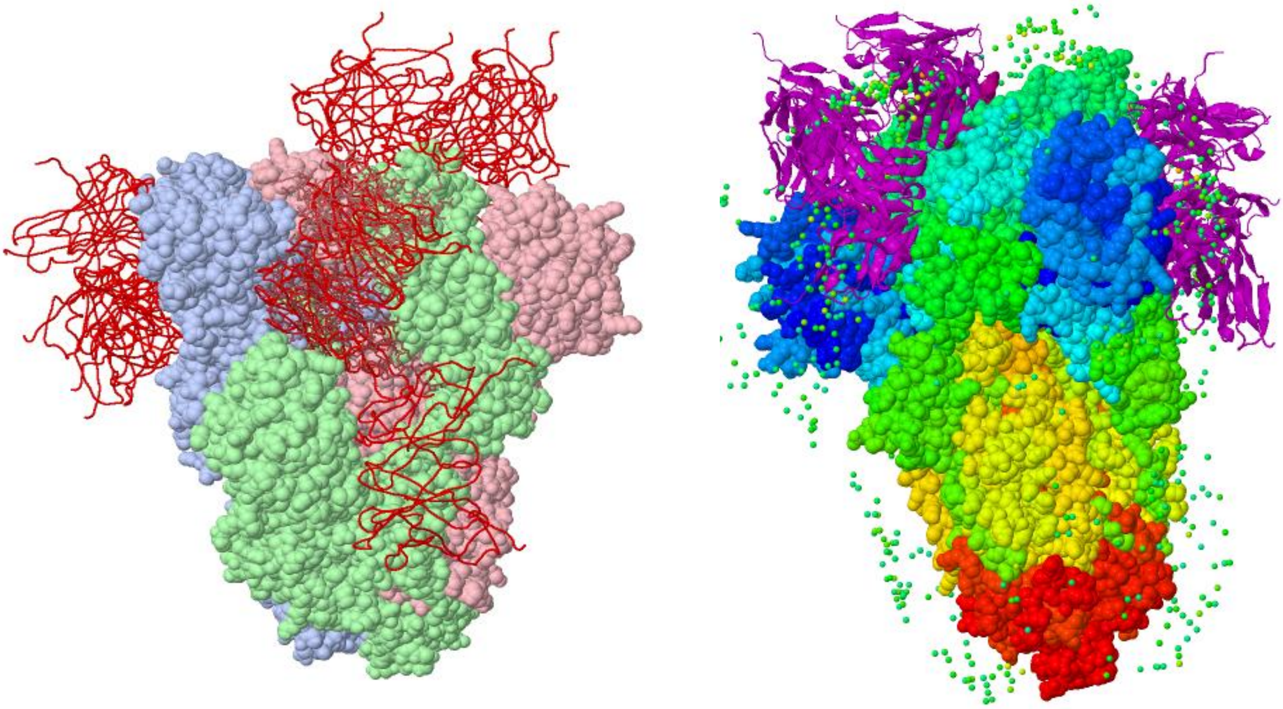

Polyspecific region binding of Fab- antibody to Spike glycoprotein of SARS-CoV-2

Closest reference gene and allele(s) from the IMGT V domain directory: *Homo sapiens* (human)

| Species | Gene and allele | Domain | Domain label | Smith-Waterman score | % identity | Overlap |
|---|---|---|---|---|---|---|
| Homo sapiens | IGHV1-46*01 | 1 | VH | 464 | 66.3 | 98 |
| Homo sapiens | IGHV1-46*03 | 1 | VH | 464 | 66.3 | 98 |
| Homo sapiens | IGHV1-46*02 | 1 | VH | 459 | 65.3 | 98 |
| Homo sapiens | IGHV1-46*04 | 1 | VH | 457 | 65.3 | 98 |
| Homo sapiens | IGHV1-3*01 | 1 | VH | 443 | 65.3 | 98 |

| Species | Gene and allele | Domain | Domain label | Smith-Waterman score | % identity | Overlap |
|---|---|---|---|---|---|---|
| Homo sapiens | IGHJ4*01 | 1 | | 95 | 86.7 | 15 |
| Homo sapiens | IGHJ4*02 | 1 | | 95 | 86.7 | 15 |
| Homo sapiens | IGHJ4*03 | 1 | | 95 | 86.7 | 15 |

Heavy chain IMGT

Light chain IMGT

Closest reference gene and allele(s) from the IMGT V domain directory: *Homo sapiens* (human)

| Species | Gene and allele | Domain | Domain label | Smith-Waterman score | % identity | Overlap |
|---|---|---|---|---|---|---|
| Homo sapiens | IGKV1-5*01 | 1 | V-KAPPA | 435 | 72.7 | 88 |
| Homo sapiens | IGKV1-5*02 | 1 | V-KAPPA | 433 | 72.7 | 88 |
| Homo sapiens | IGKV1-12*01 | 1 | V-KAPPA | 427 | 72.7 | 88 |
| Homo sapiens | IGKV1-12*02 | 1 | V-KAPPA | 427 | 72.7 | 88 |
| Homo sapiens | IGKV1D-12*01 | 1 | V-KAPPA | 427 | 72.7 | 88 |

| Species | Gene and allele | Domain | Domain label | Smith-Waterman score | % identity | Overlap |
|---|---|---|---|---|---|---|
| Homo sapiens | IGKJ4*01 | 1 | | 50 | 77.8 | 9 |
| Homo sapiens | IGKJ4*02 | 1 | | 50 | 77.8 | 9 |
| Homo sapiens | IGKJ2*01 | 1 | | 44 | 77.8 | 9 |
| Homo sapiens | IGKJ2*02 | 1 | | 44 | 77.8 | 9 |
| Homo sapiens | IGKJ5*01 | 1 | | 43 | 77.8 | 9 |

**Here is a list of examples some FDA-approved monoclonal antibody drugs in market as examples**

- abciximab (Reopro)
- adalimumab (Humira, Amjevita)
- alefacept (Amevive)
- alemtuzumab (Campath)
- basiliximab (Simulect)
- belimumab (Benlysta)
- bezlotoxumab (Zinplava)
- canakinumab (Ilaris)
